# Rapid identification of mosquito species, sex and age by mass spectrometric analysis

**DOI:** 10.1101/2022.03.07.483241

**Authors:** Iris Wagner, Linda Grigoraki, Peter Enevoldson, Michael Clarkson, Sam Jones, Jane L Hurst, Robert J Beynon, Hilary Ranson

## Abstract

**Background:** A rapid, accurate method to identify and to age grade mosquito populations would be a major advance in predicting the risk of pathogen transmission and evaluating the public health impact of vector control interventions. Whilst other chemometric methods show promise, current approaches rely on challenging morphological techniques or simple binary classifications that cannot identify the subset of the population old enough to be infectious. In this study, the ability of Rapid Evaporative Ionisation Mass Spectrometry (REIMS) to identify the species, sex and age of mosquitoes reared in the laboratory and derived from the wild was investigated.

**Results:** The accuracy of REIMS in identifying morphologically identical species of the *Anopheles gambiae* complex exceeded 97 % using principal component/linear discriminant analysis (PC-LDA) and 84 % based on random forest analysis. Age separation into 3 different age categories (1 d, 5-6 d, 14-15 d.) was achieved with 99 % (PC-LDA) and 94 % (random forest) accuracy, after including three species and storing samples at room temperature for up to 11 days. When tested on wild mosquitoes from the UK, REIMS data could determine the species and age of the specimens with accuracies of 91 and 90 % respectively.

**Conclusions:** The accuracy of REIMS to resolve the species and age of *Anopheles* mosquitoes is comparable to that achieved by chemometric approaches. The processing time and ease of use represent significant advantages over current, dissection-based methods. Importantly the accuracy was maintained when using wild mosquitoes reared under differing environmental conditions, and when mosquitoes were stored frozen or desiccated. This high throughput approach thus has potential to conduct rapid, real-time monitoring of vector populations, providing entomological evidence of the impact of alternative interventions.

## Introduction

Mosquito-borne diseases cause suffering and hundreds of thousands of deaths every year. Malaria remains a leading cause of mortality and morbidity in Africa, killing over 400,000 people annually [1] and arboviral diseases transmitted by *Aedes* mosquitoes, like dengue, chikungunya and Zika have placed more than half of the world’s population at risk. Vector control and in particular the use of insecticides in domestic environments (insecticide treated bed nets and indoor residual spraying) and mosquito breeding sites, remains the most effective tool in averting mosquito-borne infections [2]. However, increasing insecticide resistance [3, 4] is intensifying the search for new vector control tools [5, 6] and it becomes critical to have methods for evaluating the efficacy of mosquito control interventions. Epidemiological studies are considered the gold standard in evaluating the impact of vector control on disease transmission, but they are logistically and financially challenging. An alternative faster and cheaper approach could be the collection of robust entomological data that can directly reflect the risk of disease transmission.

Mosquito survival is a major determinant of their vectorial capacity. This is because after the ingestion of an infected blood meal the pathogens undergo a complex pathway of replication and dissemination to the salivary glands from where they can be transmitted further. The length of time that elapses between a mosquito ingesting a pathogen and becoming infectious is known as the extrinsic incubation period (EIP). For the malaria causing *Plasmodium* parasites, the EIP is at least 10 d [7] whereas for viruses it ranges from 6 – 15 d [8, 9].Thus, data on the age profile of a mosquito population would provide a much clearer indication of the risks of transmission (and the impact of interventions in reducing these risks) than simple counts of adult mosquito density. Indeed, the limited data available on daily survival rates of mosquitoes under natural conditions suggest that only a very small proportion survive beyond the EIP of most human pathogens [10–12].

Traditionally mosquito age has been estimated based on morphological changes observed in the ovaries of female mosquitoes. These changes can be used to assess whether a mosquito has laid eggs (parous); skilled technicians are also able to determine the number of gonotrophic cycles from examination of the ovaries. This age grading method avoids the need for specialized equipment and can be done with minimal cost under field settings, but it is labour-intensive and only provides an indirect measure of the mosquitoes’ physiological age. A range of additional methods have been explored for their potential use in mosquito age grading, including cuticular hydrocarbon analysis [13], gene expression [14, 15] and protein profiling [16–18], MALDI-TOF mass spectrometry [16, 19–21] and near infrared (NIRS) and mid infrared (MIRS) spectroscopy (recently reviewed in [22]). The chemometric methods NIRS and MIRS show the most promise, mainly due to the fast data acquisition they provide [23, 24]. However, NIRS has a high predictive power only when assigning mosquitoes into binary age categories (>7 and <7 days old) [25] and similarly, MIRS can clearly differentiate very old (> 15 days) and very young (1 day) mosquitoes, but has less accuracy in distinguishing intermediate ages [23].

Many mosquitoes belong to groups of cryptic species and, in some cases, only some members of these groups are disease vectors. For example, *Anopheles gambiae sensu lato* complex comprises at least seven sibling species of mosquitoes that are morphologically indistinguishable. Only three of these species (*An. gambiae s.s*., *An. coluzzii* and *An. arabiensis*) are important malaria vectors; each of these has differing ecology, behaviour, host-feeding preference and tolerance to insecticides [26]. Therefore, changes in species composition can have important implications for the selection of the most appropriate control tools. Multiple PCR assays have been developed to differentiate members of the *An. gambiae* species complex, but these can be challenging to perform at scale.

A method that could provide a rapid assessment of both the species and the age of mosquitoes, that is both robust to changes in the environment to which they have been exposed and insensitive to storage conditions post capture would greatly accelerate the evaluation of mosquito control strategies. In this study the potential of rapid evaporative ionisation mass spectrometry (REIMS) to predict the age and species of mosquitoes was evaluated. REIMS is based on combustion of the sample using diathermy, and subsequent ionisation and mass spectrometry of molecules present in the aerosol. It is extremely rapid (5 s per sample), generates richly detailed mass spectra and has been used in such diverse areas as surgery (where it was first developed as the iKnife - [27–29]), food fraud [30–32], in bacterial typing [33–35], monitoring of heterologous protein expression by bacteria [36] and in the analysis of rodent faecal material [37]. Most recently, we have completed a proof-of-concept series of experiments with laboratory reared *Drosophila spp* to demonstrate that REIMS can be used to discriminate species, sex and age in insects without any prior processing [38].

In this study, the ability of REIMS to provide information relevant to mosquitoes was explored. Initially, laboratory reared mosquitoes from the *An. gambiae* complex were tested as proof of principle under controlled conditions. To test the applicability of the method for field studies, REIMS was used to analyse multiple species of mosquitoes collected in North West England. We demonstrate the potential of REIMS for classification of species, sex and adult age, traits of key importance in profiling mosquito populations.

## Results

### REIMS spectra are a rich and informative data source

To generate a REIMS signal, the insect sample is combusted through application of a high frequency electric current. The resulting aerosol is collected by an airflow leading to the ion source of the instrument, where molecules are de-clustered/ionized and subsequently detected by the mass spectrometer [38]. In a single burn event, a small insect, such as an adult *An. gambiae* (adult dry weight, 250 µg, [39]) can be completely consumed in about 5-10 s (Supplementary video file 1). The mass spectra acquired at 1 Hz throughout the burn event are summed, processed and binned into 0.1 *m*/*z* bins. The negative ion mass spectra largely reflect the lipid profile of the insect and are richly informative in the range 50-1200 *m/z*. Moreover, different individuals from the same cohort yielded consistent mass spectra, but which also differed from the mass spectral patterns from other, different cohorts. The rapidity of data collection means that more than a hundred individuals can be analyzed per day. After data acquisition, the resultant data matrix, containing sample identification and binned *m*/*z* values (typically over 100 samples and over 11,000 *m/z* bins) is then subjected to multivariate analysis and analysed using machine learning and classification approaches (Figure 1).

**Figure 1.**
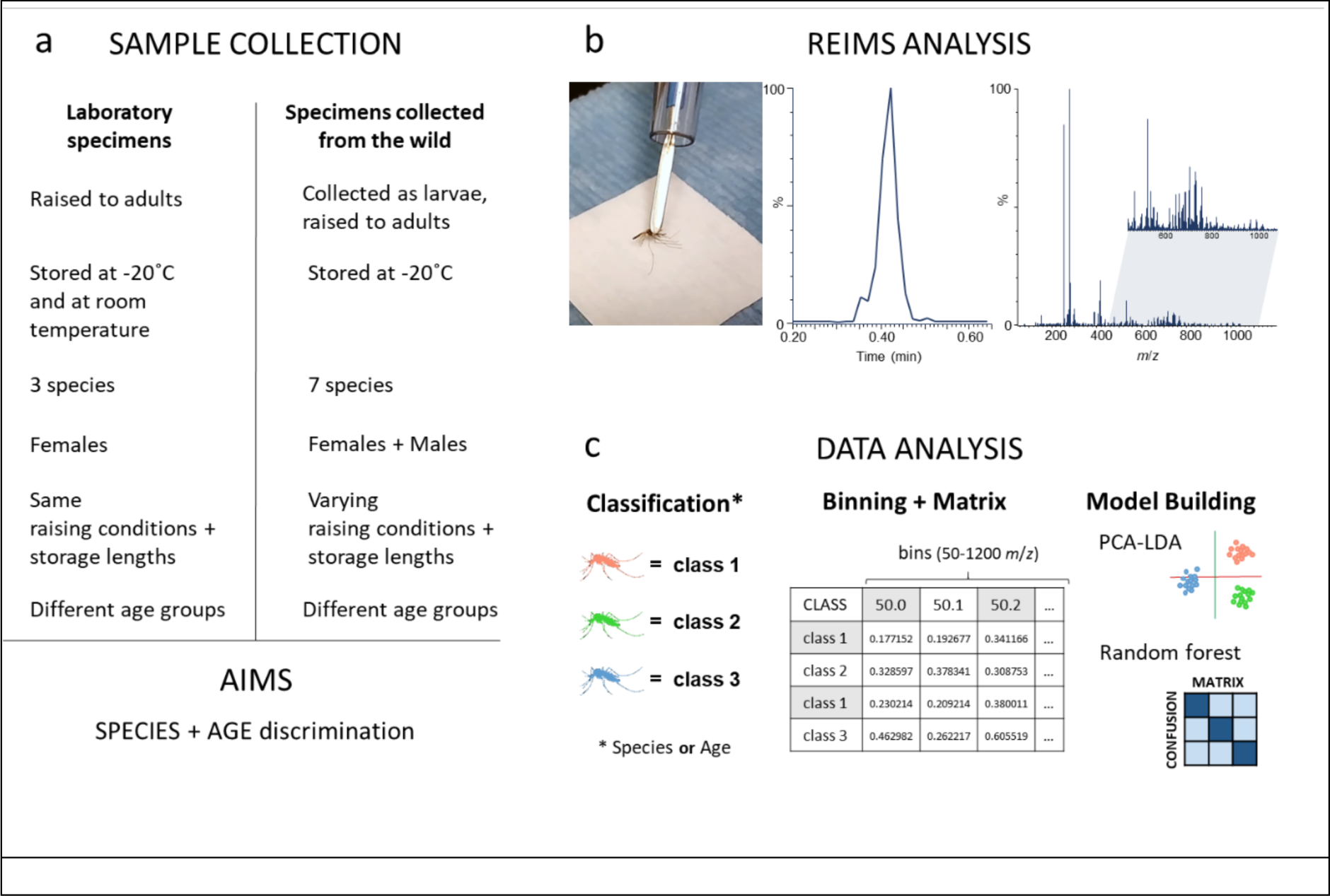
Overview of REIMS analysis of mosquito specimens. Panel a: General information about the mosquito samples used in this study. Details are listed in the methods section or mentioned with the corresponding results. Collection and treatment procedures were varied to explore the stability of the REIMS analysis. Panel b: Samples were combusted by the diathermy electric current and the resulting aerosol was evacuated through a tube and introduced to the mass spectrometer, where the ionized molecules were detected. One ‘burn event’ was generated for each mosquito and the corresponding mass spectrum was integrated over the duration of the event (typically, 5 s). Panel c: Mass spectral data were pretreated and collapsed into 0.1 m/z wide bins from 50 m/z to 1200 m/z. The resulting data matrix is then used to identify patterns through principal component and linear discriminant analysis (PCA-LDA) as well as random forest analysis, allowing sample classification and identification.

### Application of REIMS for species identification

#### Laboratory reared mosquitoes

First, we examined whether REIMS was able to discriminate three species of the *An. gambiae* species complex. We acquired REIMS mass spectra from 202 specimens, all 4-day old females comprising 54 specimens of *An. coluzzii* (strain Ngusso), 59 specimens of *An. gambiae s.s,* (strain Kisumu) and 89 *An. arabiensis* mosquitoes, (strain Moz). All mosquitoes were killed by freezing, stored at -20°C and analysed by REIMS in a randomized order. In all instances, the burn yielded a satisfactory mass spectrum. Prior to data analysis, mass spectral data were pre-processed in Offline Model Builder (Waters) in which the background signal was subtracted, spectra were mass corrected using a lock-mass standard (leucine enkephalin, 554.26 *m*/*z*) that was co-infused with each sample and finally, spectra were discretised by binning signals into 0.1 *m*/*z* wide bins.

Principal component analysis followed by linear discriminant analysis (PC-LDA), based on 90 principal components, using the Offline Model Builder software, resolved the three species effortlessly with only a single *An. coluzzii* individual being co-localized with the individuals from *An. gambiae s.s* (Figure 2, a). The first discriminant function was responsible for resolution of *An. arabiensis* from the other two *An. gambiae s.l* species whereas the second function yielded good resolution of *An. gambiae s.s* and *An. coluzzii*, a result which correlates with their genetic relatedness (Figure 2d). This finding is mirrored in the kernel density (Figure 2b) and scatter plots (Figure 2c) based on PC-LDA conducted in R. The mass spectra for the three species were very similar and exhibited only small pattern differences in the region of 600-1200 m/z between *An. arabiensis* and the other two *An. gambiae* s.I. species (the averaged spectra based on all individuals available for each species can be seen in Supplementary Figure 1). The subtlety of REIMS discrimination and the ability to resolve different species, even when they are closely related and morphologically identical (Figure 2d, 2e), aligns well with our prior observations on *Drosophila* species [38]. As a further test, the entire set of REIMS spectra from all individuals were randomly assigned to three pseudo-species classes - these could not be resolved (Supplementary Figure 2). The PCA-LDA model was also re-built using less variance (fewer principal components) and all models were cross-validated in Offline Model Builder (Supplementary Figures 3 and 4) Freezer storage might not always be feasible, especially when collecting field samples. To test whether dehydration, another frequently used approach to treat and store samples, is compatible with REIMS analysis, mosquitoes from all three species (*An. arabiensis*, *An. coluzzii*, *An. gambiae s.s*) were split into two sets: one set was killed by freezing and stored at -20°C, the other was killed through dehydration and stored with desiccant material at room temperature. Additionally, samples from each set were stored for different durations (0-10 weeks). Species separation was possible with frozen and desiccated specimens, as well as when both sample sets were combined (Supplementary Figure 5).

**Figure 2.**
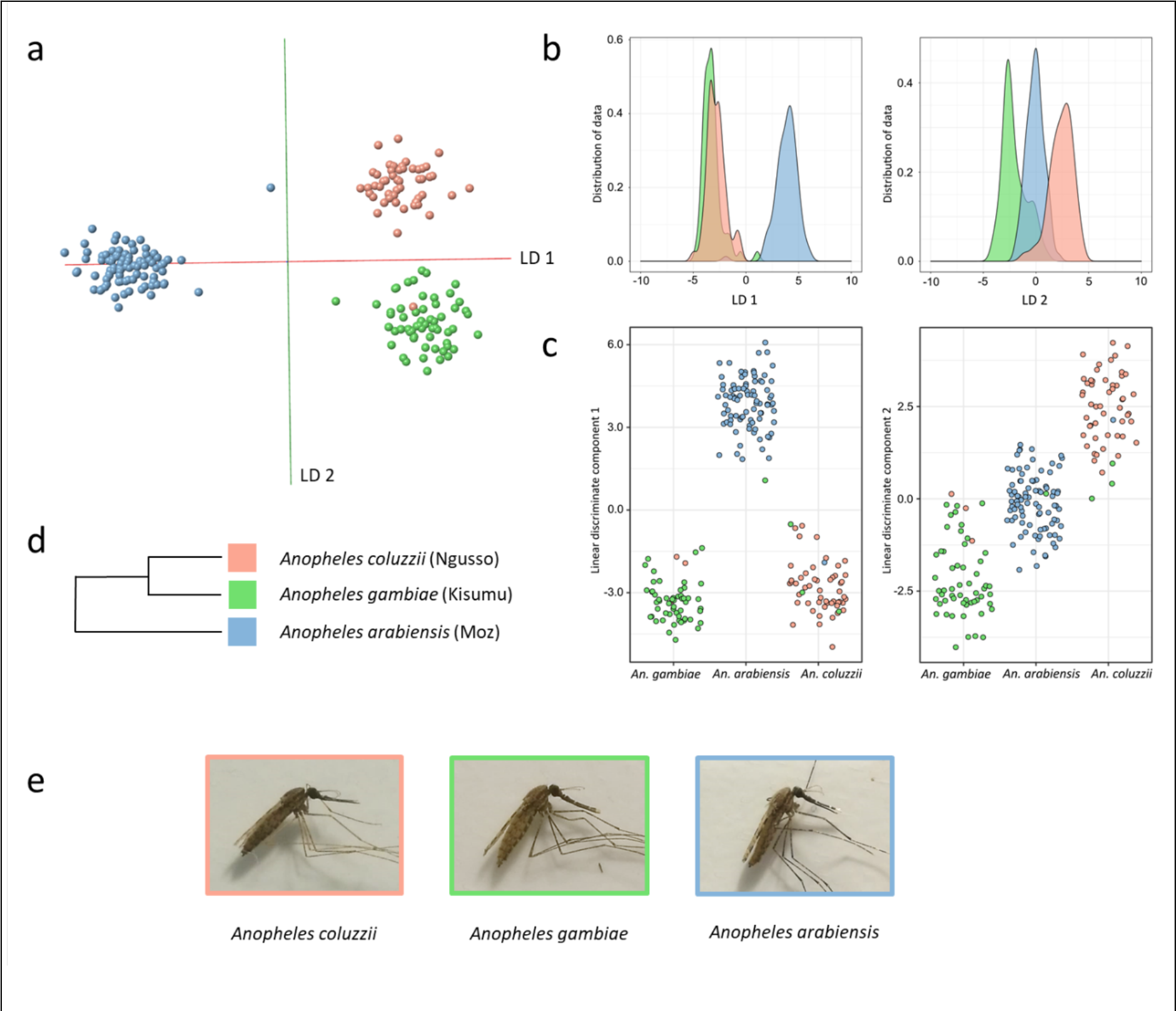
Analysis of three Anopheles mosquito species by REIMS. A total of 202 specimens from An. arabiensis (n=89), An. coluzzii (n=54) and An. gambiae s.s (n=59) were killed through freezing and stored at -20°C until REIMS analysis. All specimens were female and 4 d old. Principal component - linear discriminant (PC-LD) analysis of the REIMS data within the model building software Offline Model Builder led to a clear separation of the three species (panel a). After exporting the data matrix (incl. classifications and signal intensities after pre-processing) PC-LDA was repeated in R; results are displayed in form of kernel density (panel b) and scatter plots (panel c), shown for both linear discriminant 1 and 2. The group formation correlates with the genetic relatedness of the three groups (panel d, from [40]); the biggest variance (LD 1) supporting separation of An. arabiensis, followed by separation of An. Gambiae s.s and An. coluzzii via LD 2. Images of females from all three species are displayed in panel e.

**Figure 3.**
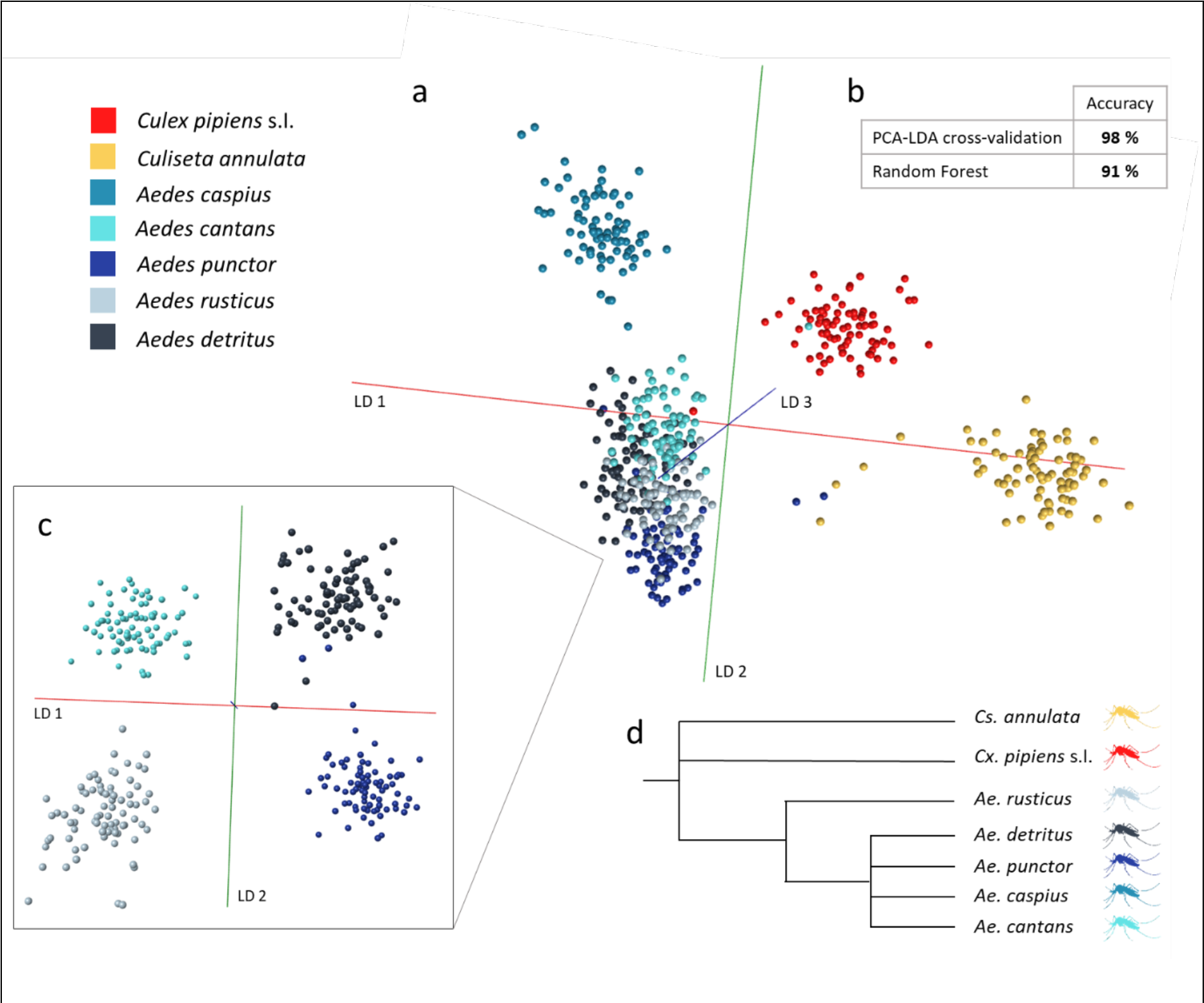
Resolution of local mosquito species using wild derived specimens. Mosquito larvae collected from the wild (Neston region, north west UK) were raised to adults (males and females, ages from 0-4 days) and identified by morphological examination, before being killed and stored at -20°C for varying lengths of time. Collection of larvae as well as REIMS analysis of stored adults occurred over several months. The data acquired for a total of seven species (80 individuals per species) was analysed in Offline Model Builder via PC-LDA using 100 PCs (section a). Visualisation was aided by removing clearly separated species groups (Cs. annulata, Cx. pipiens, Ae. caspius) from the model (section c) and axis rotation. The PC-LDA separation was resonant with the phylogenetic relationships of these species (section d). The OMB model was cross-validated (using the option ‘Leave 20 % out’ and a standard deviation of 5) and reached a classification accuracy of 98 %. Additonally, the data was analysed using a random forest algorithm, which produced a separation accuracy of 91 % (section b). Details of the cross-validation and random forest results can be found in Supplementary Figure 5.

**Figure 4.**
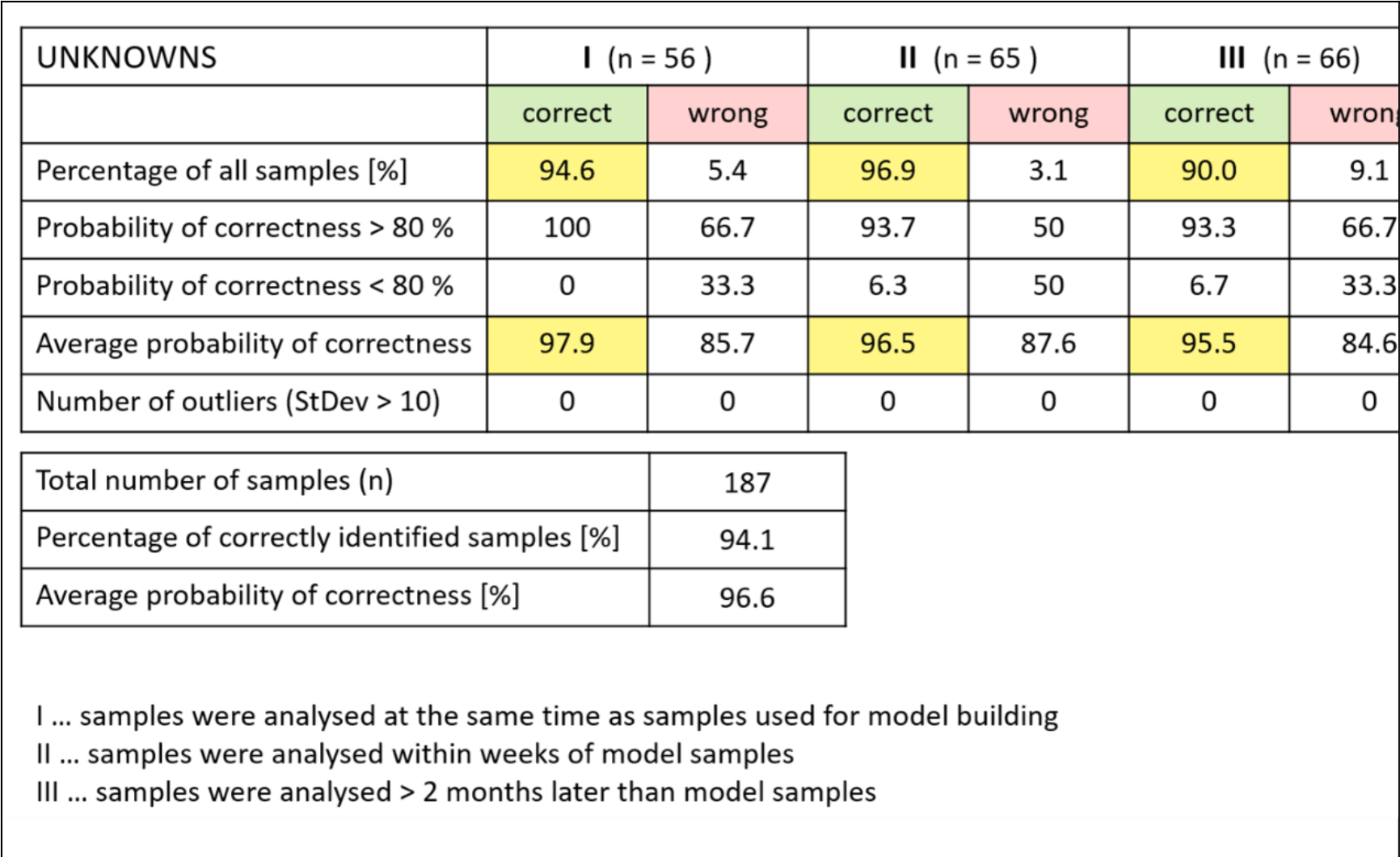
Species identification of unknown samples. The seven species PC-LDA model built in Offline Model Builder with 100 PCs was exported to the Recognition software and used to identify blind samples of unknown species (UNKNOWNS). Samples were categorised depending on their time of analysis: samples that had been analysed at the same time as the samples used for model building (I), samples analysed within weeks of model samples, (II) and samples that were analysed over two months later than the last samples included in the model building (III). The number of tested samples is stated in brackets. The percentage of correctly identified samples and the likelihood that the identification is correct are highlighted in yellow for easier comparison. Not all species were represented in every test group and identification success varied in some categories with the species; a more detailed list can be found in Supplementary Figure 11.

Whilst male and female *Anopheles* mosquitoes can be readily distinguished by morphological examination of the sexually dimorphic antennae, separation of the sexes was evaluated as a further test of the methodology. Males and females of *An. gambiae* s.s were frozen before being processed through REIMS; subsequent data analysis via PC-LDA and random forest provided a sex assignment accuracy of 98 % (in OMB) and 85 %. Detailed results and photographic examples of the sexual dimorphism are listed in Supplementary Figure 6.

#### Wild mosquito populations

Whilst laboratory specimens are reared under controlled conditions and feeding regimens, a critical test of the methodology arises when applied to specimens recovered from the natural environment. We therefore applied REIMS to the analysis of mosquitoes derived from the saltwater marshes and freshwater pools around the town of Neston on the Wirral peninsula (located in the Northwest of the U.K). This area and terrain support the proliferation of several different mosquito species that have been monitored over many years [41]. Larvae were collected and raised to adults of both sexes, which were identified to species level by morphological examination [42, 43]. In total, seven different species could be collected in sufficiently large numbers for REIMS analysis: *Culex pipiens, Culiseta annulata, Aedes caspius, Aedes punctor, Aedes rusticus, Aedes cantans and Aedes detritus.* Eighty individuals of each species were analysed in a formally randomized sequence.

Larvae were collected from their natural breeding pools and raised to adults in the same water as they were derived. Adult mosquitoes were raised throughout the year, killed by freezing and stored at -20°C for different lengths of time before being analysed by REIMS on different days spread over the course of several months. All speciemns were analysed in randomised order. The data, acquired from 0-4 d old female and male mosquitoes belonging to seven different species (80 individuals per species) were used to develop a species model (Figure 3). PC-LDA in Offline Model Builder (OMB) readily resolves *Cx. pipiens*, *Cs. annulata* and *Ae. caspius* from a tight cluster containing the other four species (Figure 3a). In fact, this four-species cluster was also resolved, as seen with a clipped and differentially projected location (Figure 3c). The tighter clustering of data points for *Ae. cantans, Ae. punctor, Ae.rusticus and Ae. detritus* is consistent with their phylogenetic proximity [44] (Figure 3d). The samples were also analysed through random forest with the resulting species determination model reaching an average accuracy of 91 % compared to 98 % for OMB (Figure 3b, Supplementary Figure 7). When species classifications are randomly assigned to samples, separation of classes fails (Supplementary Figure 8). The model proves that discriminant patterns can also be found within samples displaying natural variability, providing an important step towards the application of REIMS with field caught mosquitoes.

As the sample population of wild-derived mosquitoes consisted of both male and female individuals, classification of samples by sex was attempted. Males and females from the species *Aedes detritus* were resolved with a correct classification rate of 83 % for PC-LDA (cross-validation in OMB) and 82 % for random forest analysis (Supplementary Figure 9). To add more complexity to the process, a further sample set included males and females from four different species (*Ae. detritus*, *Ae. punctor*, *Ae. rusticus*, *Ae. cantans*). The increase in variance mildly impacted the separation process, with correct identification rates to 82 % (PC-LDA in OMB) and 77 % (random forest; Supplementary Figure 10).

### Blinded identification study

To test the utility of REIMS further, we used the seven-species model in Figure 3 as a predictive tool. A further 200 samples (collected as larvae, harvested as adults and assigned to species by independent morphological examination) were blinded to the analyst, and the REIMS data were submitted to the model for species prediction. Overall, 94 % accuracy was achieved for blinded samples (Figure 4).

Wild, trap-caught adults were also presented to the predictive model for identification. The samples used to build the predictive species model had been collected from the field as immatures and do not fully represent wild adult mosquitoes (no sugar or blood feeding). Nevertheless, an average of 68 % of samples were correctly identified; mosquitoes that had been analysed at the same time as the model samples were identified at even higher rates of 87 and 91 % (categories I and II, Supplementary Figure 12).

Analysis of mosquito larvae We also explored the ability of REIMS to resolve the immature (larval) stages of mosquitoes into species. While we did not establish the species of the larvae by morphology, many of the pools in the Neston area from which the mosquitoes were collected are essentially mosquito monocultures. Many larvae were harvested, a subset of which was immediately analysed with REIMS, while the remainder were allowed to develop into adults. The emerged adults were then used to predict the species heterogeneity of the larval population in each sampling pool and confirmed that the pools contained single species (100 % of the adults identified were the same species, either *Ae. puncto*r or *Ae. detritus;* detailed numbers in Supplementary Figure 13a).

The sample data was analysed through PC-LDA, in OMB and R. In both cases the separation of *Ae. detritus* and *Ae. punctor* larvae was very distinct with only 20 principal components (Figure 5, a+b). After using the data matrix to conduct random forest analysis (10 repeats), the test samples were almost always identified correctly, leading to correct identification rates of 100 % (*Ae. detritus*) and 99

**Figure 5.**
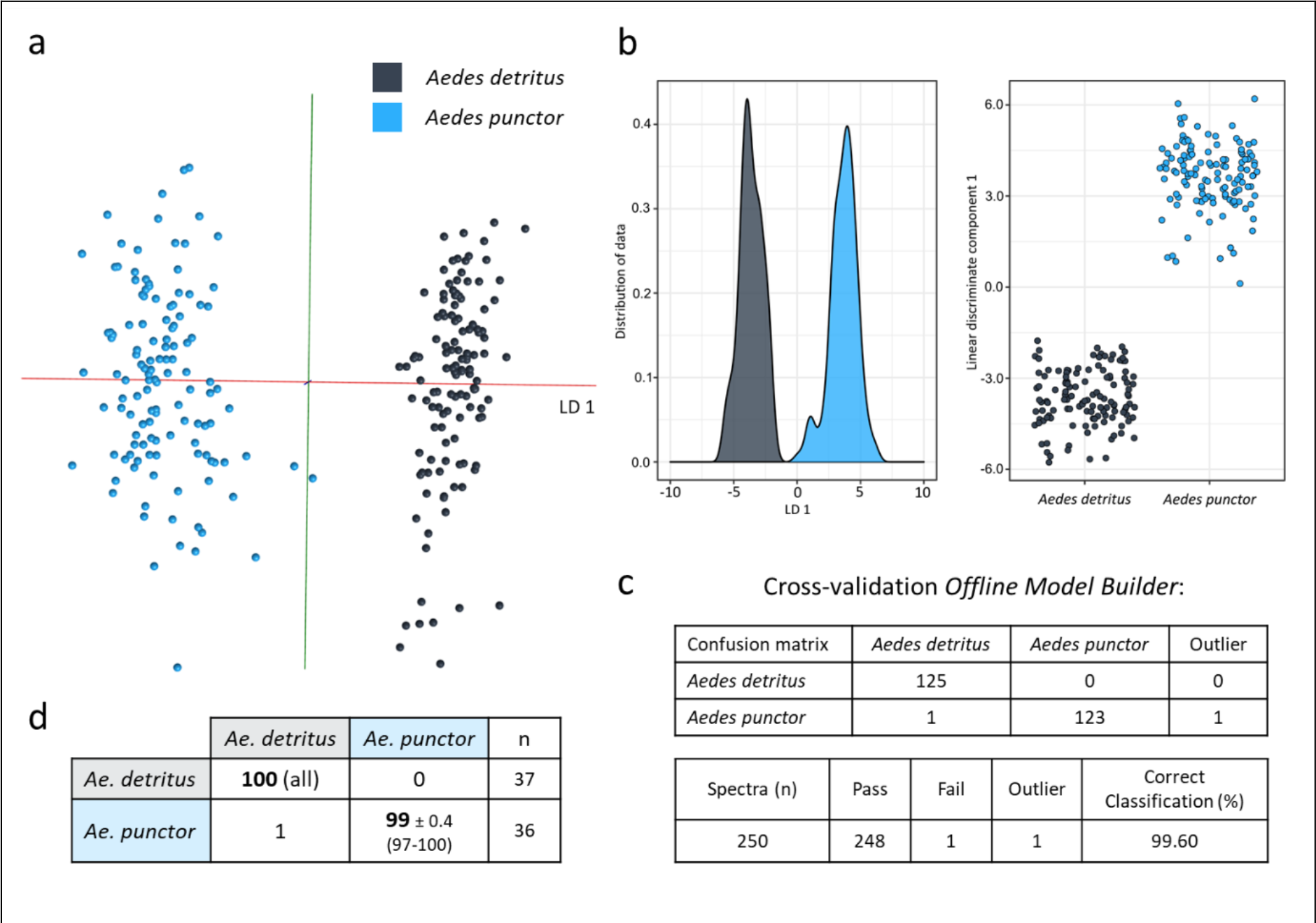
Species separation based on immature specimens. Immature mosquitoes were obtained by filtering water collected from different pools, followed by 2-3 rinsing steps before killing and storing them at -20°C. Samples, mostly 3^rd^ instar larvae were analysed with the same REIMS settings used for adult specimens. Their species was confirmed by sampling the same pool of larvae and raising them to adults before, during and after taking samples to be used for this model. The differences detected by PC-LDA are visualised in the form of an OMB model (a) and kernel density and scatterplots produced in R (b). The separation was put to test via cross-validation in OMB (c) and random forest analysis with training/test dataset split of 70 %/30 % (d). The separation of Ae. detritus (n=125) and Ae. punctor (n=125) larvae required little information/variance, reaching very distinct separation with only 20 principal components for PC-LDA. When testing the random forest model, samples were identified correctly most of the time leading to correct identification rates of 100 % (Ae. detritus) and 99 % (Ae. punctor).

% (*Ae. punctor*) (Figure 5, d). Cross-validation of the OMB model resulted in one misclassification and one outlier out of 250 samples (Figure 5, c). Seeing this distinct separation through PC-LDA, principal component analysis alone was performed in OMB, which resulted in samples clustering into species groups along principal component three (Supplementary Figure 13b). The larval-based species model was also re-built using randomly assigned classifications; the resulting model was devoid of any separation or sample clustering (Supplementary Figure 13c).

### Application of REIMS for age profiling

A characteristic of particular significance in vector control is that of female mosquito age. Establishing the age of individual mosquito specimens could, in combination with regular sampling and high-throughput analysis, allow determination of population age profiles and hence the risk of disease transmission. Age distribution would inform and support vector control strategies, particularly when in control validation stages, with a reduction in the median age of the population being a more reliable proxy of the public health value of a new tool than metrics based on population density. Indeed vector control trials frequently include measurements of the proportion of the population that are parous (having laid at least one egg batch) as a crude measure of the proportion of ‘older’ adults, in this context being mosquitoes ≥ three days [45]. The ability to rapidly and reliably separate mosquitoes into different age groups would be transformative for mosquito surveillance. We therefore established whether REIMS possessed the ability to resolve mosquitoes according to age.

#### Laboratory reared specimens

Reports of the lifespan of female *An. gambiae s.l* mosquitoes range from 2 to 43 days, with longevity under laboratory conditions typically greatly exceeding that in the field where predation, environment and disease all reduce survival [46]. For the initial tests, we included female *An. gambiae s.s* (Kisumu strain) raised for 0–1, 2, 3, 4 and 5 d under standard insectary conditions. Mosquitoes of all five age groups were killed by freezing and stored at -20°C the same day. After REIMS analysis of specimens of all ages, randomly sequenced, PC-LDA in Offline Model Builder led to a clear clustering according to age, with the location of each age group reflecting the progressive age of specimens (Figure 6, a). Further PC-LDA in R and visualization through 3D plots with different triads of the top four linear discriminants reveals a similar picture of age progression along LD 1. The values of LD 1 contain enough information to provide some distinction for all groups but with overlap; the second to fourth LDs add further resolution of specific groups, improving the overall separation.

**Figure 6.**
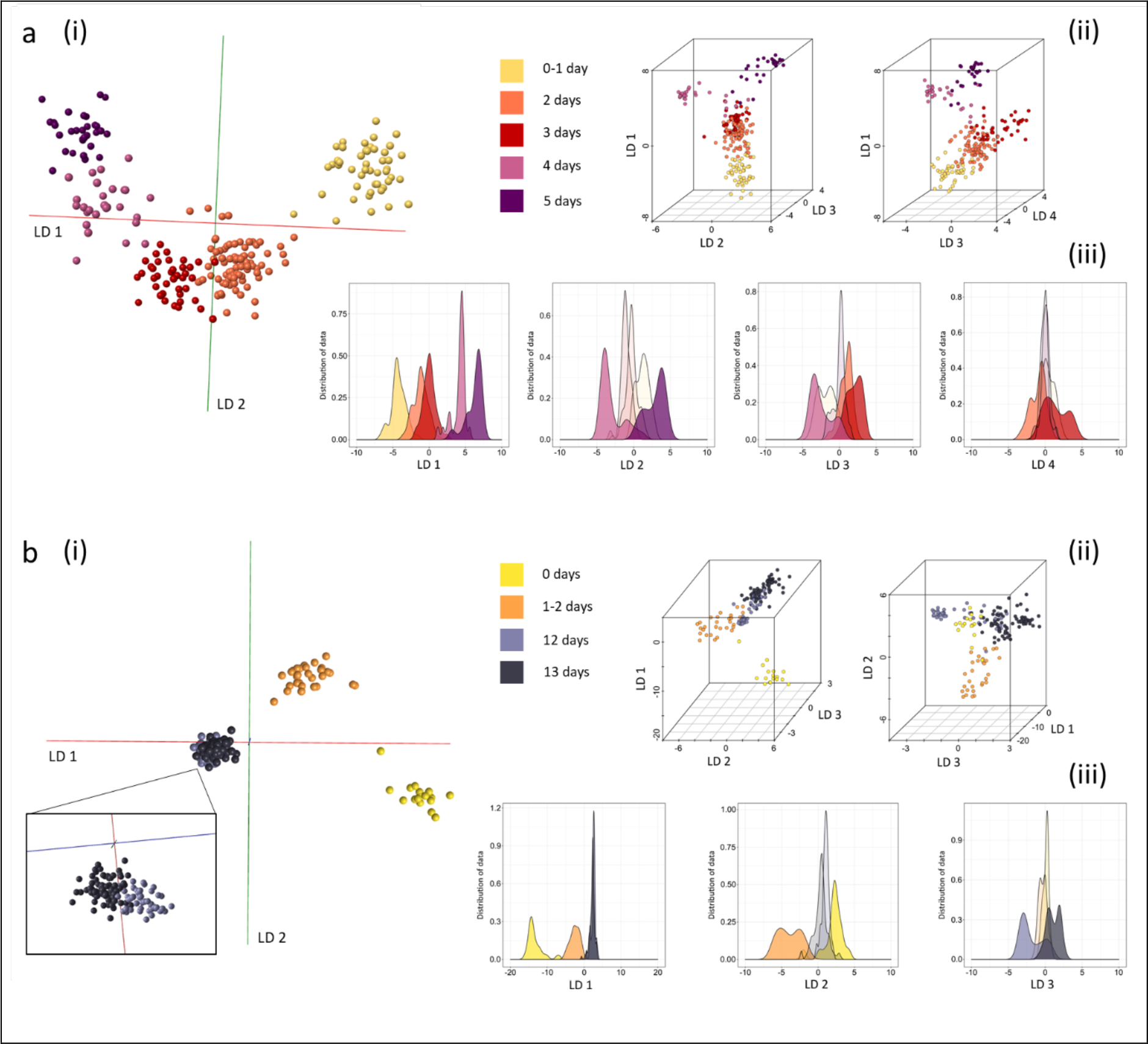
Discrimination of Anopheles gambiae mosquitoes by age. Two groups (panels a and b) of Anopheles gambiae s.s (Kisumu strain) specimens of different ages were killed by freezing before REIMS analysis. Mosquitoes were raised on sugar solution, regardless of age. Differences between age groups were explored by PC-LDA and visualized using OMB (i) as well as R, in form of 3D models (using different linear discriminant combinations) (ii) and kernel density plots for each LD. The difference between classes in model a (built with 100 PCs in OMB and R) is only 24 h, nonetheless a distinct group formation can be observed with chronological positioning within the 3D space, leading to a transition from younger to older samples. Model b (built with 88 PCs in OMB and 85 PCs in R), comprising young and old mosquitoes (2 groups each), revealed greater variance between the young classes than between the old. Sample numbers per class, model a: 0-1 day (n=47), 2 days (n=84), 3 days (n=39), 4 days (n=27), 5 days (n=30); model b: 0 days (n=17), 1-2 days (n=29), 12 days (n=46), 13 days (n=83).

A second set of *An. gambiae s.s* was analyzed, focusing on more broadly spaced age groups: very young (0 days or 1-2 d) and old (12 d, 13 d) (Figure 6b). Again, samples grouped according to their age class, however, with unequal separation between the three classes. The difference between mosquitoes that had just emerged (day 0) and those that are 1-2 days old was greater than the difference between adults at 12 d and 13 d, which could reflect metabolic and developmental changes in the first 24 h after emergence from pupae. This difference in variance can also be clearly seen in either scatter or kernel density plots; both LD 1 and 2 are adding to the separation of the young mosquitoes, whereas LD 3 was able to provide limited variance to distinguish between 12 and 13 d mosquitoes.

Both age models were also built with fewer principal components (Supplementary Figure 14), to test robustness, as well as with randomly assigned pseudo-classifications (Supplementary Figure 15) to confirm that separation is based on age-related differences between classes.

The ease with which age could be resolved and the clear separation of the 0-1 day old mosquitoes might reflect a major change in lipid deposits post-emergence. However, averaged mass spectra of 0-5 day old mosquitoes are visually similar and do not reveal a highly evident variation in their mass spectral patterns (Supplementary Figure 16). Nevertheless, sample clusters based on age related variance were formed, making age a REIMS accessible parameter. In contrast to species or sex, age is a continuous parameter and might benefit from conversion to discontinuous categorisation to help define class boundaries and increase separation and identification accuracy. A reduction in class numbers (from 5 to 3 and from 4 to 3), achieved by combining neighbouring classes, reduced the overlap between age groups and improved definition of class boundaries (Supplementary Figure 17) as well as increased correct classifications rates obtained through cross-validation of PC-LDA models in OMB (Supplementary Figure 18).

**Figure 7.**
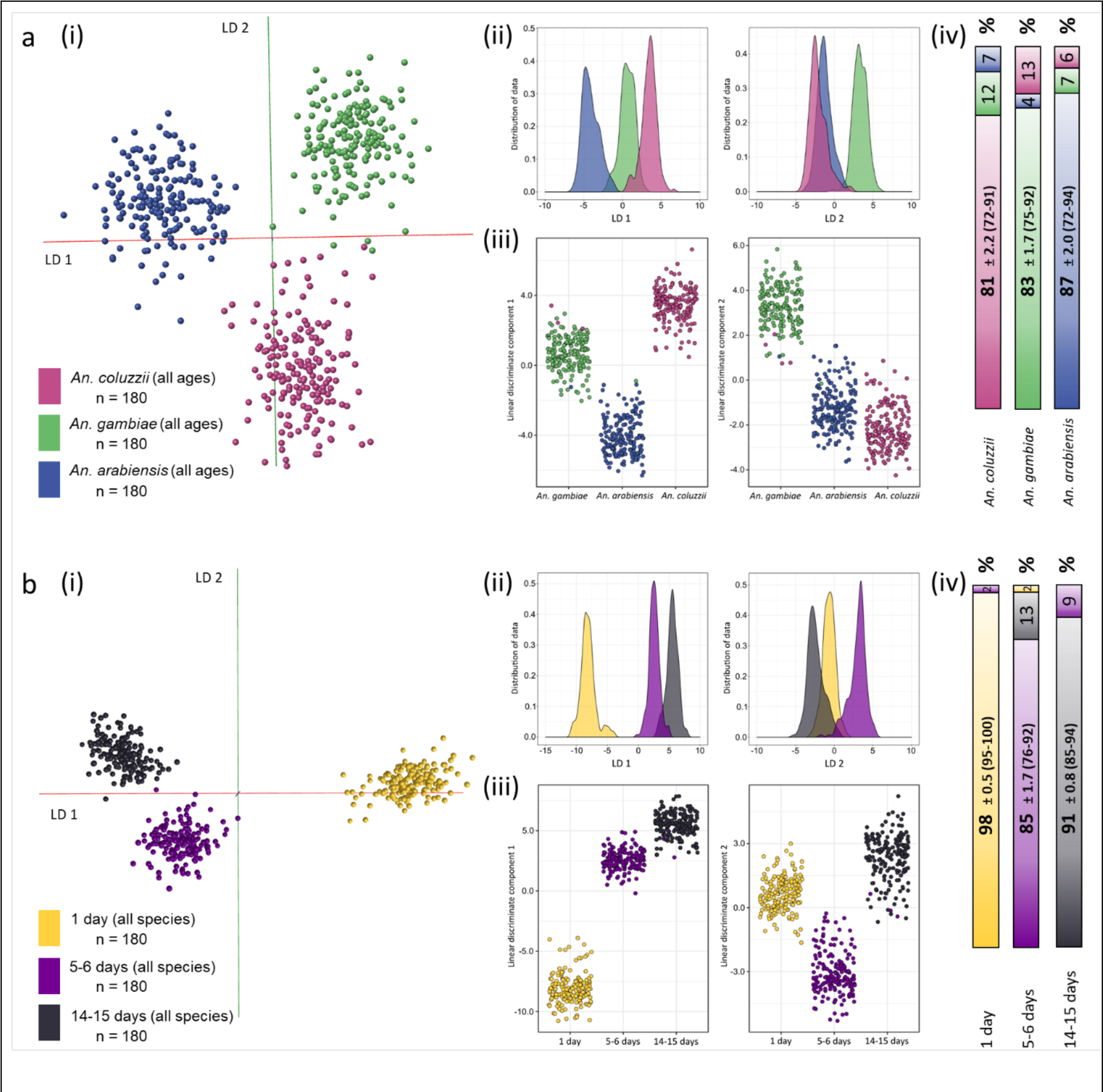
Separation of species and age by REIMS Mosquitoes (total n=540) were raised to three different age groups (1 day, 5-6 days, 14-15 days) for each of the three species (An. coluzzii, An. gambiae s.s, An arabiensis, see text). The specimens were killed by dehydration and stored at room temperature with desiccant for 1-1.5 weeks prior to analysis. The samples were used to build two models: one separating the three species (panel a) and one separating the three age groups (panel b). Data was processed using PC-LDA in Offline Model Builder (using 100 PCs) (i) and R (using 235 PCs), latter visualized for each linear discriminant (LD 1 and LD 2) separately in form of kernel density (ii) and scatter plots (iii). The data matrix, exported from the Offline Model Builder software, was additionally analyzed using the random forest algorithm in R; 70 % of the samples were used for model building, 30 % as test samples. The classification results are depicted as a bar graph, showing percentages of correctly and wrongly identified samples (iv). Depicted are the average values of 10 random forest repeat runs ± the standard error of the mean, with the range of accuracy values that were achieved in brackets.

**Figure 8.**
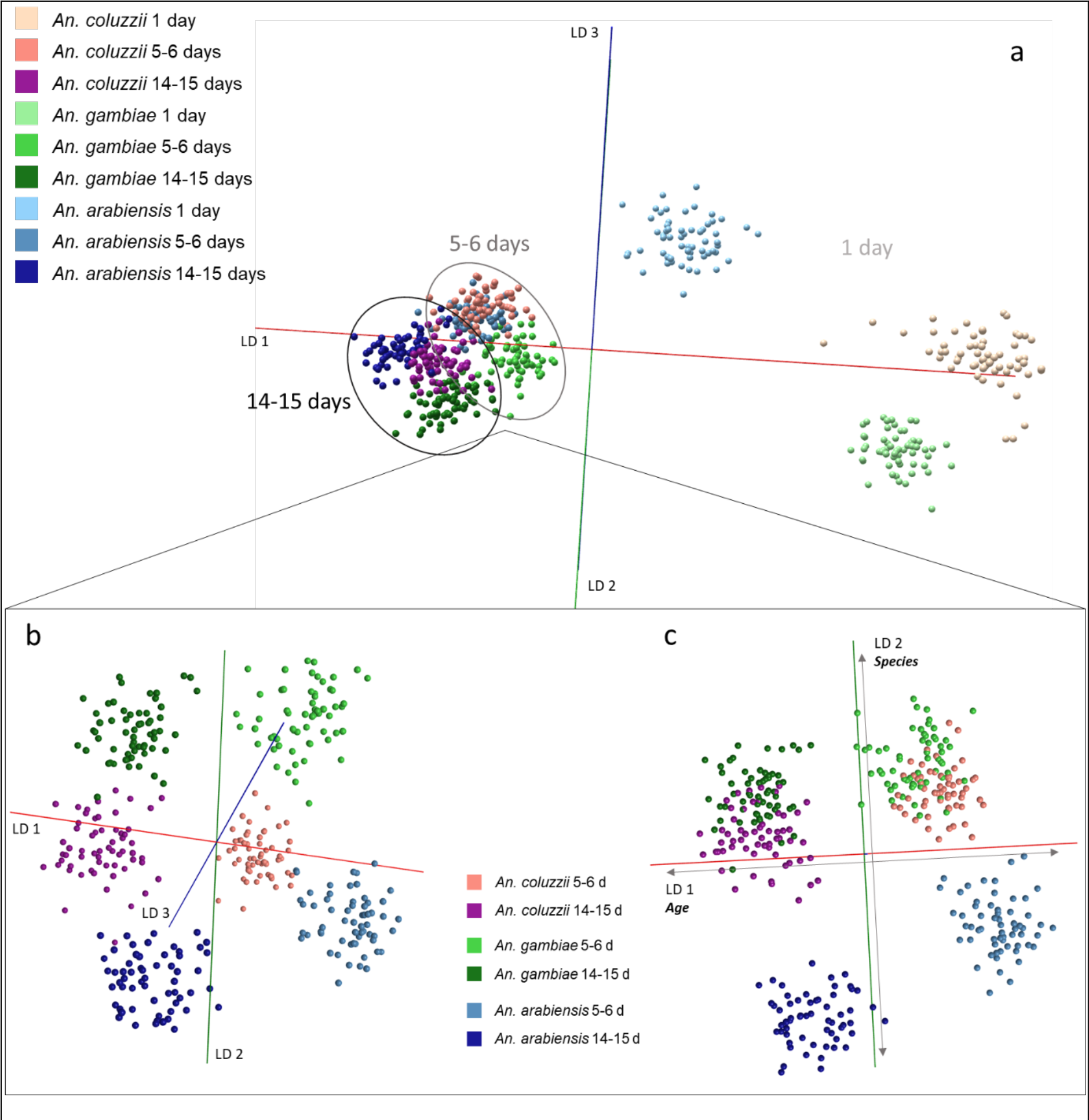
Two-factor model combining species and age information The same samples used to build the species and age models in Figure 7 were used to construct a two-factor model, comprising nine classes, each containing species and age information. Separation of all nine classes (60 specimens each) was attempted using PC-LDA in Offline Model Builder (based on 100 PCs) (section a). Due to wide dispersion of the 1 day old groups, spatial resolution of the 5-6 and 14-15 day old groups in the 3D space is hindered. To help visualize the separation, the 1 day old sample groups were removed (section b). The largest variance in the data set (LD 1) is correlated with age, followed by species separation of An. arabiensis enabled by LD 2 and lastly days separation of An. gambiae s.s from An. coluzzii days based on LD 3 (section c).

Following the successful separation of laboratory-reared mosquitoes according to their species or age, potentially confounding factors were introduced to the next sample set to further test the concept of mosquito characterization through REIMS, particularly in the context of field-caught samples, in which circumstance, freezing might not be an option. Female specimens from *An. arabiensis (*strain Moz), *An. gambiae* s.s (strain Kisumu) and *An. coluzzii* (strain Ngusso) were each raised and sampled into three different age groups: 1 d, 5-6 d and 14-15 d. The specimens were then killed through dehydration and stored at room temperature with desiccant material for 1-1.5 weeks before REIMS analysis. These data were used to build two different models: one to resolve species, the other to explore resolution according to age. Thus, mosquitoes of different ages are part of the species model (Figure 7a) and age separation was tested on aggregated data from all three species (Figure 7b).

Data obtained from 540 mosquitoes were first analyzed by PC-LDA in Offline Model Builder (i), followed by PC-LDA in R, depicted as kernel density (ii) and scatter plots (iii), and lastly used to conduct random forest analysis (iv). The random forest construction and analysis was repeated 10 times, using randomly selected model construction samples and test samples each time. The average performance statistics, and number of correct and incorrect classifications confirm a high level of discrimination (Figure 7). Despite increasing variability in the data set by inclusion of specimens of different ages, separation of species was still successful, even though samples within groups are slightly more scattered and overall group resolution was slightly reduced. The average accuracy achieved through random forest analysis was 87 % correct identification for *An. arabiensis*, 81 % for *An. coluzzii* and 83 % for *An. gambiae s.s*. The greatest degree of misclassification was clearly between *An. gambiae s.s* and *An. coluzzii* (12-13 %). Cross-validation of the PC-LDA model built in OMB resulted in an even higher correct classification rate of 98 % (Supplementary Figure 19).

**Figure 9.**
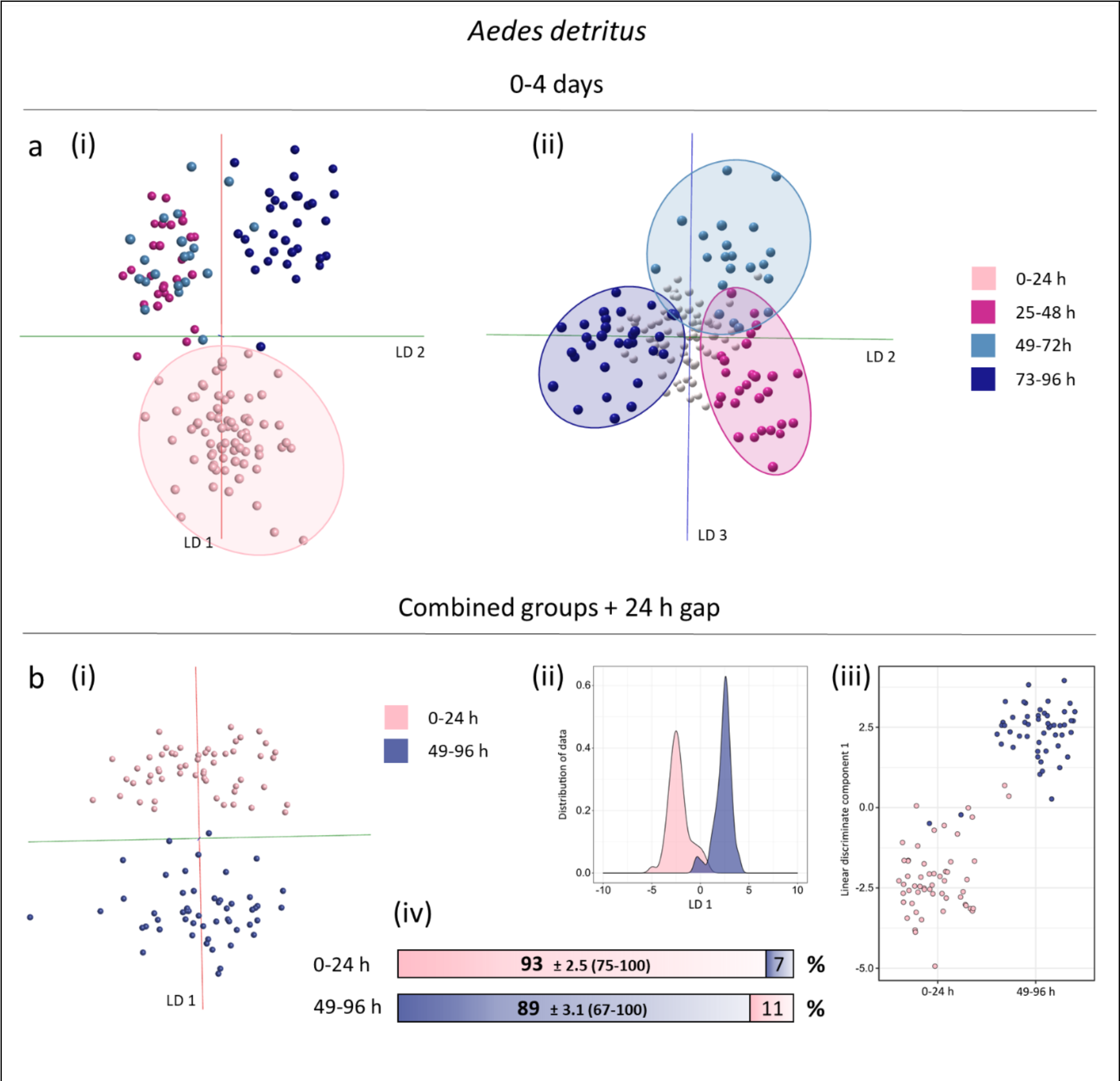
Discrimination of Aedes detritus mosquitoes by age. Ae. detritus mosquitoes, emerged from larvae collected from natural pools, were killed by freezing at different ages ranging from zero to four days (males and females). As a first step samples were sorted into four age groups for PC-LDA (based on 75 PCs), which resulted in a definitive grouping according to age (panel a), with linear discriminant 1 separating just emerged mosquitoes (0-24 h) (i) and linear discriminants 2 and 3 separating specimens which are between 25 and 96 h old (ii). To improve separation a 24 h gap was introduced and two groups merged (panel b). PC-LDA shows a clear reduction in class overlap in OMB (using 50 PCs)(i), which can also be observed in the kernel density (ii) and scatter plots (iii) produced in R (using 55 PCs). Random forest analysis, using a 70 %/30 % ratio for training and testing, resulted in identification accuracies of 93 % for 0-24 h old specimens and 89 % for 49-96 h mosquitoes (iv). Sample numbers per class for model in panel a: 0-24 h (71), 25- 48 h (28), 49-72 h (21), 73-96 h (31). Sample numbers per class for model in panel b: 0-24 h (55), 49-96 (52) - sample numbers from the 0-24 h class were reduced for random forest analysis.

The second model, separating mosquitoes according to their age, displays a tight clustering of samples within age groups and a very distinct separation between age groups. As previously observed, the very young mosquitoes (1 day old) resolved readily from older specimens and show the biggest separation. Both analytical approaches, PC-LDA and random forest, concur that the 1-day old specimens are easily separated from the rest, leaving a pronounced separation in all three LDA plots (b i-iii) and scoring the highest identification accuracy in random forest (98 %). Overall, the age model achieved an average accuracy of 91 %, making this age separation species independent. Cross-validation of the PC-LDA based model produced an average accuracy of 99 % (Supplementary Figure 19). Thus, although samples had been treated very differently, compared to the previous species and age models (Figure 2 and 6), REIMS analysis nevertheless resulted in information rich mass spectral data allowing for clear separation of species and highly accurate age discrimination. Both models (species and age) were also re-built in R with fewer principal components to confirm that classes can still be separated using less variance and potentially a smaller number of separating factors (Supplementary Figure 20).

For species resolution, the m/z bins contributing a high level of discrimination were different from those driving age discrimination (Supplementary Figure 21), suggesting that it should be possible to derive a two-dimensional model. Both characteristics, species and age, would be of interest when identifying trapped mosquitoes, leaving two options: acquire data and test with two separate models or build a two-factor model capable of providing both types of information at once. To test whether a two-factor model could be a viable option, the same data set used to build the species and age models (Figure 7) was split into nine classes, each representing two factors, species and age (Figure 8). Regardless of the added difficulty of splitting variance to enable separation of nine classes based on two properties each, samples were successfully grouped and separated. Surprisingly, separation was facilitated in ways similar to the separate species and age models. The average random forest accuracy of the species model was 84 %, lower than the mean accuracy of the age model (91 %). The two-factor model assigned more variance to the age separation than the separation of species, which is apparent in the clustering according to age rather than species, followed by the separation of the three clusters (1 day, 5-6 days, 14-15 days) along linear discriminant 1 (Figure 8a). From comparison of the 5-6 day old and the 14-15 day old classes, age is separated based on linear discriminant 1, followed by species resolution of *An. arabiensis* from the other two *Anopheles* species through LD 2 and further separation of *An. Gambiae s.s* from *An. coluzzii* via LD 3 (Figure 8b, c). This analysis augurs well for profiling analyses with wild caught specimens. Cross-validation following PC-LDA in OMB produced an average correct classification rate of 97 % (Supplementary Figure 22), random forest analysis was slightly less successful, identifying only 79 % of samples correctly (Supplementary Figure 23). PC-LDA was also performed in the R environment (Supplementary Figure 24) producing similar results and misplacing only 7 out of 360 samples.

Following random forest analysis, the variables driving the separation of the nine classes were aggregated and compared to m/z bins previously identified as important in the species and age models (Supplementary Figure 21). Three out of five variables had been identified previously, which confirms their importance for species and age separation respectively (Supplementary Figure 25). Lastly, the separate age and species models as well as the nine-class model were all rebuilt using randomly assigned classifications to ensure the separating patterns are not based on unrelated noise; separation failed for all three models (Supplementary Figure 26).

#### Wild-derived specimens

Larvae of *Ae. detritus* collected in the Neston area were allowed to develop to adults and were harvested at different ages. We used these semi-wild samples to establish whether age specific patterns could be retained in the presence of confounding factors, such as impact of the natural environment during larval development. PC-LDA of 0-4 day old *Ae. detritus* mosquitoes (males and females) resolved into four 24 h age classes with clear resolution of the four groups (Figure 9a). As expected for continous data, the classes are very close or slightly overlap, consistent with the previous analysis of 0-5 day old *An. gambiae (*Figure 6). The three age groups ranging from 25 to 96 h were separated by LD 2 and 3, whereas mosquitoes that had just emerged are separated by the largest variance component (LD 1, Figure 9a). This finding is particularly encouraging, as it matches previous age models based on laboratory raised specimens. In a subsequent analysis, the 25-48 h class was removed to introduce a gap in the continous age range. Additionally, 49-72 h adults were combined with 73-96 h adult to increase separation and to help increase sample numbers. Separation of the two new age classes was clearly improved when analysed through PC-LDA in Offline Model Builder, repeated PC-LDA in R replicated the result, showing only a few samples are being misclassified in the process (Figure 9b). The data matrix exported from OMB was also subjected to random forest analysis, based on 70% training data/30 % evaluation data. The 0-24 h old mosquitoes reached a correct identification rate of 93 %, of the 49-96 h mosquitoes 89 % were correctly identified (Figure 9b). Aggregated age classes, with at least a 24 h period between them, are clearly beneficial for class separation and the gain in performance more than offsets the loss of resolution.

The laboratory mosquito based age model proved to be stable even in the presence of multiple species. To test whether this also holds true for wild-derived specimens, mosquitoes from four different species (*Aedes detritus, Culiseta annulata, Aedes rusticus, Aedes punctor)* were used to build age models with the same classifications as seen in Figure 9 (Supplementary Figure 27). The average percentage of samples correctly identified through random forest dropped only slighly from 91 % (Figure 9b) to 88 % upon introduction of different species. As before, this analysis was performed with combined classes and a 24 gap between them. Reduction of classes alone can increase separation performance signigicantly, which was tested using 0-48 h and 49-96 h classes. A comparison of PC-LDA models, built using different sets of classes, show the step-wise improvement of separation for the single-species as well as the multi-species model (Supplementary Figures 28 and 30). All models were cross-validated in Offline Model Builder to prove the increase in classification performance (Supplementary Figures 29 and 31).

After successful separation of age groups using wild-derived mosquitoes, which were unfed and killed by freezing, more samples were added to the exisiting 0-24/49-96 h model to test the stability of pattern-based age grading (Figure 10). A broader range of samples was included in this extended model, purposely adding variance and potentially confounding factors. A third age group was introduced to represent older mosquitoes, ranging from 168 to 240 h (7-10 days). While no intervention took place between emergence and killing when raising the previously used wild-derived mosquitoes (Figure 3 and 9), these adults were raised in three different conditions: kept dry, kept with water or fed with sucrose solution. Furthermore, samples from different species were added; four species (*Ae. detritus, Ae. rusticus, Ae. punctor and Cs. annulata*) are represented in the first two age classes (0-24 h and 49-96 h) and 2 species (*Ae. detritus and Ae. caspius*) are part of the older age class (168-240 h). Whereas all 0- 24 h old specimens had been killed by freezing, some mosquitoes of the older age classes had died naturally before collection. They were nonetheless added to the sample pool to add further variance to the model. Linear discriminant analysis in OMB, based on 100 principal components, produced distinct separation of the three age groups (Figure 10a). Similar results can be seen when conducting LD analysis (based on 95 PCs) in R, the scatterplots show that only a few samples are confused between the 0-24 and 49-96 h classes, and that all 168-240 h old mosquitoes are correctly located (Figure 10b). The latter can be explained when viewing the variance distribution in the kernel density plots; the oldest group is separated clearly via LD 1, whereas separation of 0-24 h and 49-96 h is mostly based on LD 2, meaning there is sligthly more variance benefitting the distinction of 7-10 day old specimens (Figure 10c). In random forest analysis, the 0-24 h old test samples scored the highest identification accuracy with 94 %, followed by the 168-240 h old class with 92 % of samples corrently identified and the 49-96 h old group with 84 % correct identification (Figure 10d).

**Figure 10:**
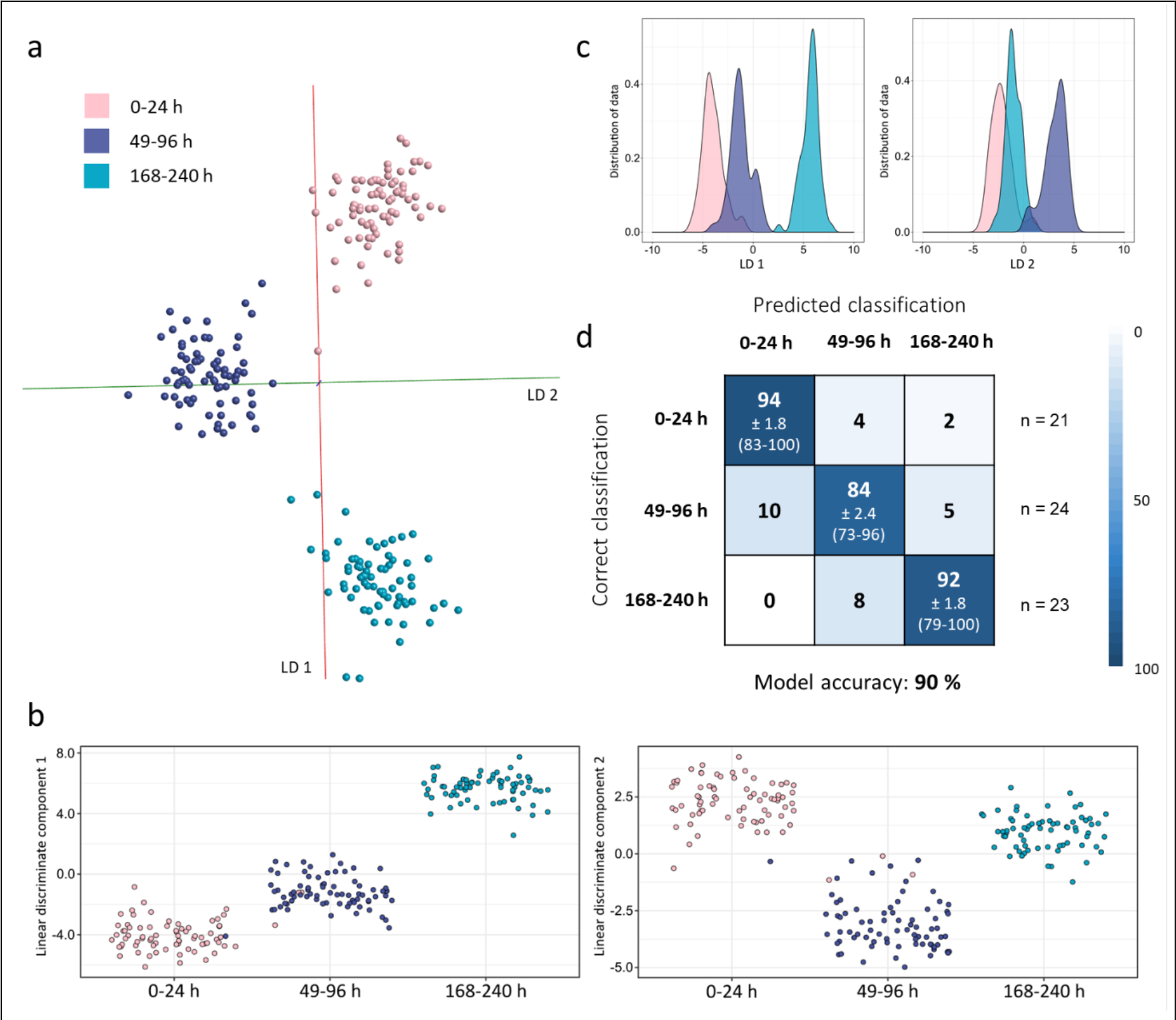
Species independent separation of age groups using wild-derived mosquitoes. In addition to the Ae. detritus samples used in Figure 9, samples from four other species and another age class (168-240 h, 7-10 days) were introduced, including adult mosquitoes that were kept with or without water or fed sucrose solution. A portion of specimens was killed by freezing, some died due to other reasons shortly before being collected; all were stored at -20°C before analysis. PC-LDA achieved separation of the three age groups (0-24 h, 49-96 h, 168-240 h), as can be seen in the OMB model (based on 100 PCs) (section a) as well as the scatterplots (section b) and kernel density distributions (section c) created in R (based on 95 PCs). An overall 90 % of test samples were correctly identified during random forest analysis with group specific accuracies of 94 % for newly emerged specimens (0-1 day old), 84 % for 2-4 day old mosquitoes and 92 % for the 7-10 day old group (section d). Sample numbers per class: 0-24 h (75), 49-96 h (75), 168-240 h (69).

All three age models for wild-derived specimens were re-built in OMB after randomly assigning them age classifications to ensure that previous separations had been achieved through age-related variance (Supplementary Figure 32). A comparison of cross-validation results of models with correct or randomly assigned classifications underpins the presence of age-related differences in the collected data; classifications rates for models with random sample assignment are drastically lower (Supplementary Figure 33). Even reducing the number of principal components and therefore variance used for LD analysis did not hinder class separation, it merely reduced the extent of separation and class resolution (Supplementary Figure 34).

The ion bins used to age-grade the semi-wild mosquitoes during random forest analysis differ from the bins identified for the age model based on laboratory-raised specimens (Supplementary Figure 35). This could be caused by environmental effects during their immature stages, i.e. the difference between laboratory and field pools, but could also be related to other factors, such as different raising conditions during adulthood, differing storage conditions (room temperature and freezer) or perhaps the ageing pattern simply varies for different mosquito species. While these experiments prove that age-grading through REIMS can be extended beyond mosquitoes raised in controlled laboratory environments, it is yet unknown whether models based on laboratory-raised or semi-wild mosquitoes (derived from the wild as immatures) are capable of age-grading fully-wild mosquitoes, which will require further exploration.

## Discussion

A mass spectrometric method such as REIMS has compelling advantages in the context of field studies. First, because there is zero sample preparation, the generation of the aerosol can be applied to a broad range of biological material, including tissues, [27–29], foodstuffs [30–32, 47], faecal pellets [37], bacterial colonies [33–35] and plant materials [48]. Within a few seconds, the burn event generates a complex, richly detailed mass spectrum. Atypically, there is no attempt to interpret this spectrum in terms of the molecular species that are introduced into the instrument. Rather, it is the pattern of ions that are generated that create a complex vector, and the matrix of samples and m/z bins are suitable for multivariate data analysis whether supervised or unsupervised.

We have primarily explored the ability of REIMS to serve as a rapid method to create species and age profiles of mosquitoes using supervised machine learning approaches such as PC-LDA and random forest. This information on mosquito demography is key for implementing and assessing the impact of vector control strategies, as even a small change in the lifespan of mosquito species that are vectors of disease can have a big impact on pathogen transmission. We have shown that models generated using REIMS data can discriminate the three main vectors of malaria in the *An. gambiae* species complex: *An. coluzzii*, *An. gambiae s.s* and *An. arabiensis*, with an accuracy of 81-87%, when using lab-raised individuals of different ages. In addition, seven species belonging to the Culicidae family were distinguished with an accuracy of 91 %, using male and female wild derived mosquitoes, marking a major milestone in testing REIMS as an identification tool for insects. This species identification accuracy is slightly higher than, for example, that reported by the MIRS method coupled with machine learning (∼76% for lab-raised *An. gambiae* and *An. arabiensis*, [23]) and similar to recently developed deep-learning MIRS models (93% for lab-raised *Anopheles* mosquitoes and 95% for wild derived females, [49]).

REIMS could also discriminate three age groups of lab-raised *Anopheles* mosquitoes: very young (1 day old), middle (5-6 days old) and old (14-15 days old) with a mean accuracy of 91 %; and three age groups of wild derived *Culicine* mosquitoes (0-1 day, 2-4 days and 7-10 days) with an accuracy of 90 %. By comparison, NIRS can place lab-reared *An. gambiae s.s* into binary groups (less than and greater than 7 days old) with an accuracy of 80 % [25], but this predictive ability is lost after day 7 and drops to a maximum of 69.6 % when analysing wild derived mosquitoes [50]. Likewise, the accuracy of the MIRS method in predicting chronological age is highly variable (average for lab-raised *An. gambiae s.s.* ranging from 15 %-97 %), with the very young (1 day) and very old (15 days) mosquitoes being more accurately predicted than the intermediate ages. A deep-learning MIRS model achieved an accuracy of 89 % for resolution of wild-derived *Anopheles* mosquitoes into three age classes (1-4, 5-10 and 11-17 days old). REIMS has species and age predictive accuracy that is equal to, if not higher than other chemometric methods. REIMS retained this performance for field derived samples and is robust to field collection and storage, a critical step for its translational development.

In addition to the high predictive accuracy, there are several other factors that need to be addressed for an age-grading methodology to become a valuable tool. First, the methodology must be compatible with samples that are collected under field conditions – our ability to resolve desiccated rather than frozen samples is a significant gain. Secondly, age predictions must not be unduly influenced by temperature and humidity, diet, intraspecific phenotypic variation or physiological status (bloodfed, gravid etc.). Whilst results from *Culicines* sampled from the UK are encouraging, further studies of the robustness of the predictions under field conditions are needed. Thirdly, although the mass spectrometry data does not require interpretation or molecular assignment, in the current configuration, the REIMS source is coupled to a relatively sophisticated mass spectrometer; to reduce the capital expenditure it would be desirable to explore instrumentation providing less resolution. Currently, data is binned in 0.1 m/z increments to reduce complexity prior to multivariate analysis. If extended to 1 m/z wide bins, the resulting data matrix can still be used to separate species as well as age classes (Supplementary Figure 36). This opens the possibility of coupling the REIMS source to a lower resolution instrument. Alternatively, a more focused analysis could be created that emphasizes specific ions, for example by isolation of precursor ions, fragmentation and quantification of product ions (selected reaction monitoring). Both approaches would improve compatibility with more routine instrumentation, such as single or triple quadrupole instruments, that would have a substantially lower price tag. Finally, although acquisition and recognition are extremely rapid, there is scope for evolution of multivariate data methods that might increase confidence and the speed of profiling. The combination of a simple, rapid ionization method with real-time data acquisition and data reduction makes REIMS worthy of further consideration.

## Acknowledgments

This work was funded by grants from the Low Carbon EcoInnovatory programme and BBSRC (BB/L014793/1) to RJB and JLH. LG thanks the Wellcome Trust for a Postdoctoral Fellowship, 215894/Z/19/Z. We are grateful to Dr Philip Brownridge for exceptional instrument support and to Joscelyn Sarsby and Natalie Koch for their help with the REIMS set-up and data analysis. We are pleased to acknowledge Marion Morris and Henrietta Carrington-Yates for assistance with rearing mosquitos.

## Methods

Laboratory raised mosquitoes: Three *Anopheles* mosquito strains maintained at the Liverpool School of Tropical Medicine were used in this study: an *An. gambiae* s.s strain called Kisumu (Kenya, 1975), an *An. coluzzii* strain called N’gusso (Cameroon, 2006) and an *An. arabiensis* strain called Moz (Mozambique, 2009). All three strains were reared at 26 ± 2 °C and a relative humidity (RH) of 80 ± 10 % under a L12:D12 h light:dark cycle with a 1-h dawn and dusk. All stages of larvae were reared in distilled water and fed on ground fish food (Tetramin tropical flakes, Tetra, Blacksburg, VA, USA). Adults were provided with 10% sucrose solution *ad libitum*.

*Anopheles* laboratory-raised mosquitoes were collected as pupae and placed in paper buckets for a 24h emergence period (day 0). Thereafter non emerged pupae were removed or kept for an additional 24h (in cases where two consecutive age groups are reported). Age profiling samples were collected at different days post emergence, as depicted in the results section. Females were separated from males based on clear morphological differences (sexual dimorphism of the antenna) and aspirated into paper buckets; males were discarded (exception: sex separation experiment). Mosquitoes were killed either by freezing at -20°C, in which case samples were stored in the freezer until the day of analysis, or through dehydration by placing the buckets at 36-38°C overnight without a water source. In the latter case samples were stored the next day in plastic tubes with silica gel at room temperature until the day of analysis.

Wild-caught mosquitoes: *Ae. detritus* and *Ae. caspius* larvae were collected from pools in the Dee estuary, as outlined in [41]. Other species were collected as larvae from pools at locations in the Neston area (summarised in Supplementary Figure 37). Specimens that were analysed in their larval stage were filtered and rinsed with MilliQ water 2-3 times before being killed by freezing. Specimens that were collected as larvae and then analysed in their adult stage were reared in glass jars in their original water and fed intermittently with yeast (Zipvit Brewer’s yeast). After emerging as adults, they were captured in nets attached to the original container that were emptied every 24 hours. Adults were killed by freezing, and their species and sex identified morphologically according to standard keys [42, 43]. The pool of larval origin, date of larval collection, date of emergence and date of killing (and thus, adult age at death), sex and species were recorded for each individual. All samples were then stored at -20 °C until analysis. Wild adult mosquitoes (almost exclusively females) were captured in traps (Mosquito Magnet, Lancaster, PA, USA) using carbon dioxide and octenol as attractants. Trapping occurred over a two day period every week between March and November 2019 in up to 4 sites in the Neston area [41], Figure 1a).

REIMS analysis: Samples were analysed via a rapid evaporative source (REIMS, Waters, Wilmslow, UK) attached to a Synapt G2Si instrument ion mobility equipped quadrupole time of flight mass spectrometer (Waters, UK). The specimens were burned/evaporated using a monopolar electrosurgical pencil (Erbe Medical UK Ltd, Leeds, UK), which was connected to a VIO 50 C electrosurgical generator (Erbe Medical UK Ltd, Leeds, UK), providing electrical current, and to the source inlet via plastic tubing. A black conductive rubber mat, placed underneath the samples, acted as a counter electrode and facilitated the flow of electric current. To avoid inhalation of fumes during analysis, the burning process was performed within a fume box (Air Science, Lydiate, Merseyside, UK). Insects were analysed using a 40 W setting on the generator and the cutting option of the pencil. To increase conductivity and to protect the counter electrode during analysis, specimens were placed on a piece of glass microfibre paper (GFP, GE Healthcare Whatman) on top of a wet paper surface (moistened with MilliQ water).

While burning the entire biomass of single specimens, the aerosol was aspirated through the pencil and the attached 3m long tubing into the REIMS source, using a nitrogen powered venturi valve on the source inlet. To increase the aerosol capture, a wide bore piece of plastic tubing was additionally placed over the tip of the electrosurgical pencil. A whistle incorporated into the Venturi tube guided the aerosol as well as a lock mass solution of leucine enkephalin (Waters, UK) in propan-2-ol (CHROMASOLV, Honeywell Riedel-de-Haën) into the source. This also filters the incoming aerosol to prevent larger particles from entering the inlet capillary. Inside the source, the ionised particles were declustered through contact with a heated impactor (Kanthal metal coil at 900 °C).

Mass spectra were acquired in negative ion mode at a rate of 1 scan per second between 50 *m/z* and 1200 *m/z*. The sample cone and heater bias were set to 60 V. Instrument calibration was performed daily in resolution mode using a 0.5 mM solution of sodium formate (flow rate 50 µl/min). The lock mass solution (0.4 µg/ml, leucine enkephalin in IPA) was continuously introduced during sample analysis at 30 µl/min. For each experiment, samples were analysed in a formally randomised order on one or more days.

## Data analysis

The raw mass spectra were imported into the model building software package Offline Model Builder (OMB-1.1.28; Waters Research Centre, Hungary), which allows separation of sample groups (classifications) based on principal component analysis (PCA) and linear discriminant analysis (LDA). Data were additionally analysed using R (version 3.6.1) [51] and the R Studio environment [52], by PCA and LDA, as well as random forest analysis.

For Offline Model Builder, the burn events of the analysed specimens were defined individually, summing all the MS scans within each chosen area. The option to create one spectrum per sample was selected. Other pre-processing parameters included the intensity threshold, which was set between 4e5 and 9e5 (depending on the background baseline), spectra correction using the lock mass (leucine enkephalin, 554.26 m/z) and background subtraction. To reduce the complexity of the mass spectral data, all acquired data points from 50 to 1200 m/z were combined into mass bins, each 0.1 m/z wide. Subsequent model calculation was based on principal component and linear discriminant analysis (PCA- LDA).

The models built by Offline Model Builder were cross-validated (leaving out 20 % of data, for outliers the standard deviation multiplier was set to 5) to obtain the correct classification rate, as well as the number of failures and outliers and a matrix displaying the number of correctly and incorrectly identified samples of each classification. To additionally test discrimination, sample classifications were randomised and the data were re-analysed, the expectation being a random distribution of samples and the inability to achieve separation.

For further analysis with R, the data matrix of each model was exported as a .csv file from Offline Model Builder, containing information about classification and the relative intensities for every mass bin. The matrices were used to perform random forest analysis in R using the package ‘randomForest’ [53]. The data sets were randomly split into a training set (approx. 70 % of the data) and a test set (approx. 30 % of the data). Random forest results are displayed in form of confusion matrices. Trees were conducted 10 times for every model (using a different, randomly selected subset of samples for training and testing every time); the numbers of correctly identified and confused samples were turned into percentages and averaged. The optimal number of trees and *mtry* value were determined during the first analysis of each model and kept the same for each repeated analysis. A second R package, called ‘randomForestExplainer’ [54], was used to identify the most informative bins/ions that were driving class separation. PCA-LDA was also performed within the R environment, using the in-built package ‘stats’ and the package ‘MASS’ [55] and results visualized in form of kernel density plots and 2D- and 3D-scatter plots created using ‘ggplot2’ [56] and ‘scatterplot3d’ [57]. PC-LDA based models were built with the number of principal components (PCs) giving the best separation before overfitting the model (that number can slightly vary between OMB and R). To ensure separation of classes is still possible when using less variance, models were also built using only a quarter of the maximum number of PCs possible for each model.

All raw data files (comprising 2,920 Waters MassLynx raw files) are freely available in the University of Liverpool data repository (http://dx.doi.org/10.17638/datacat.liverpool.ac.uk/1565).

## Supplementary Material

**Supplemental Figure 1:**
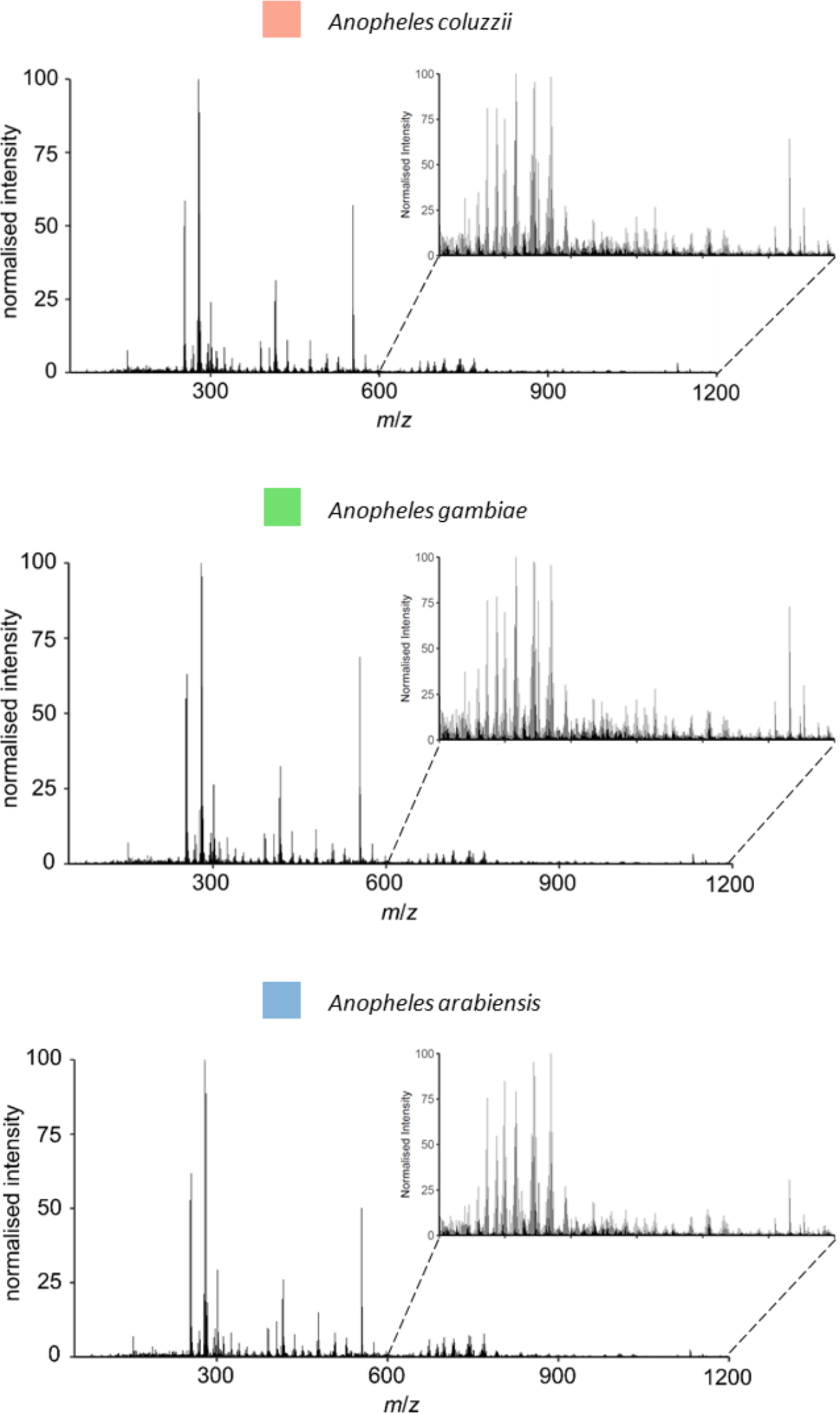
The data matrix, obtained after processing and binning the mass spectral data in Offline Model Builder, was used to create averaged mass spectra for all three species. Each mass spectrum represents an average of all samples available for each species (Anopheles coluzzii n=54, Anopheles gambiae n=59, Anopheles arabiensis n=89).

**Supplemental Figure 2:**
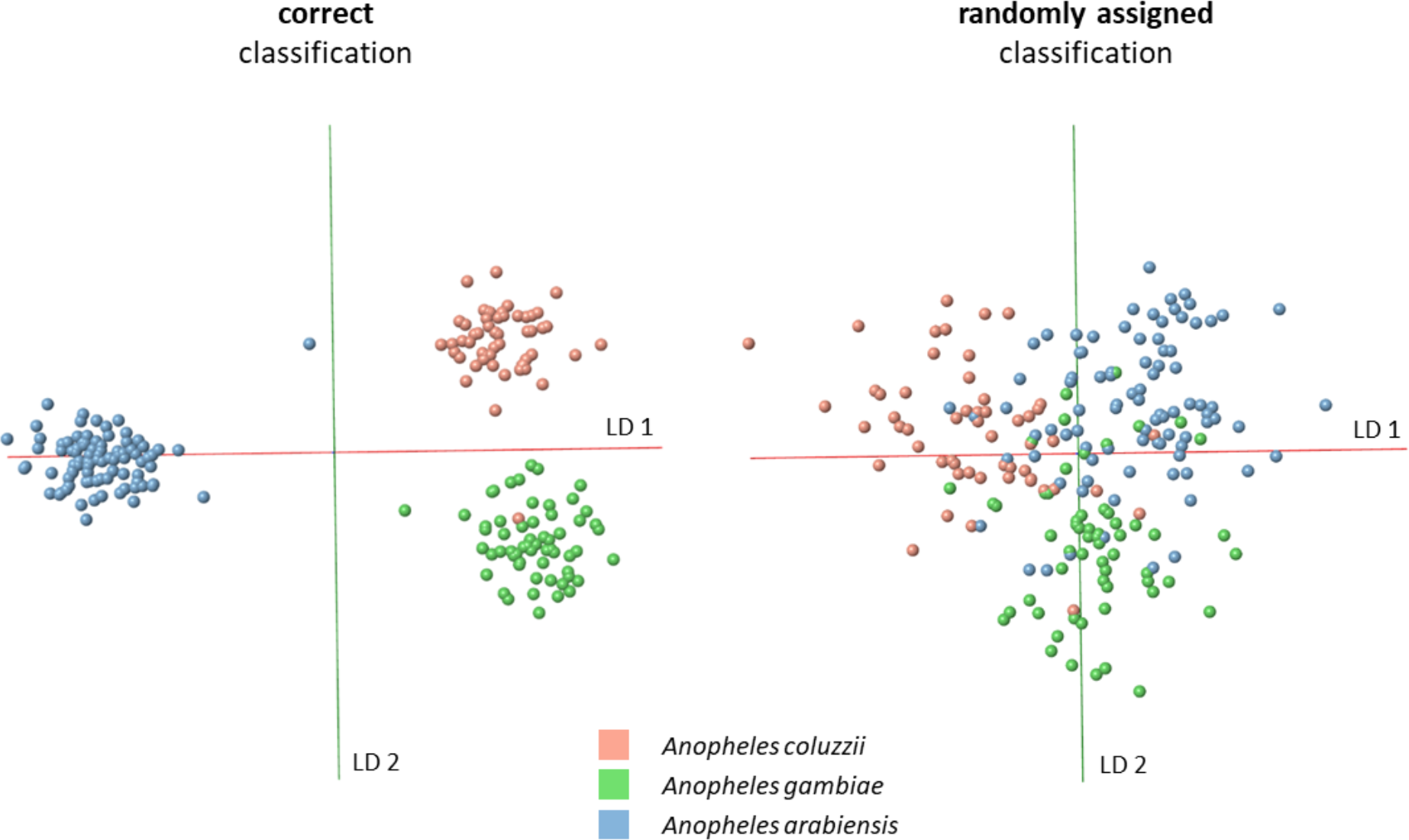
The PC-LDA model separating An. coluzzii, An. gambiae and An. arabiensis, built in Offline Model Builder using 90 PCs (left), was re-built after randomly assigning classifications to samples (right). The random classification model, also based on 90 PCs, displays no separation of the three species; samples are widely dispersed and groups strongly overlap.

**Supplemental Figure 3:**
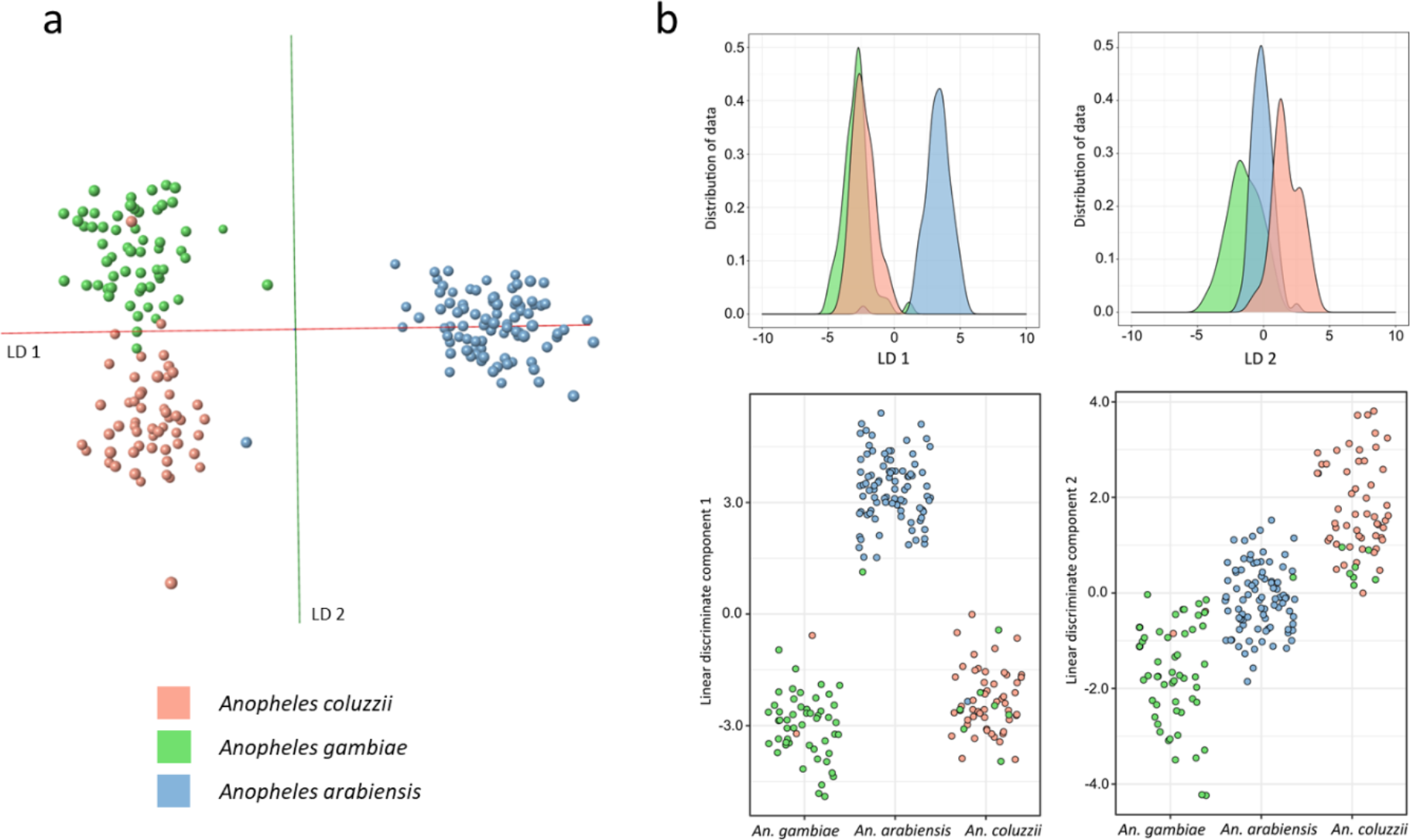
The model separating An. coluzzii, An. gambiae and An. arabiensis was re-built using a lower number of principal components. PC number was decreased to 50, which is ¼ of the maximum number possible. As can be seen in the OMB model (a) as well as the kernel density- and scatter plots (b), reduced variance in the model still resulted in a clear separation of all three species.

**Supplemental Figure 4:**
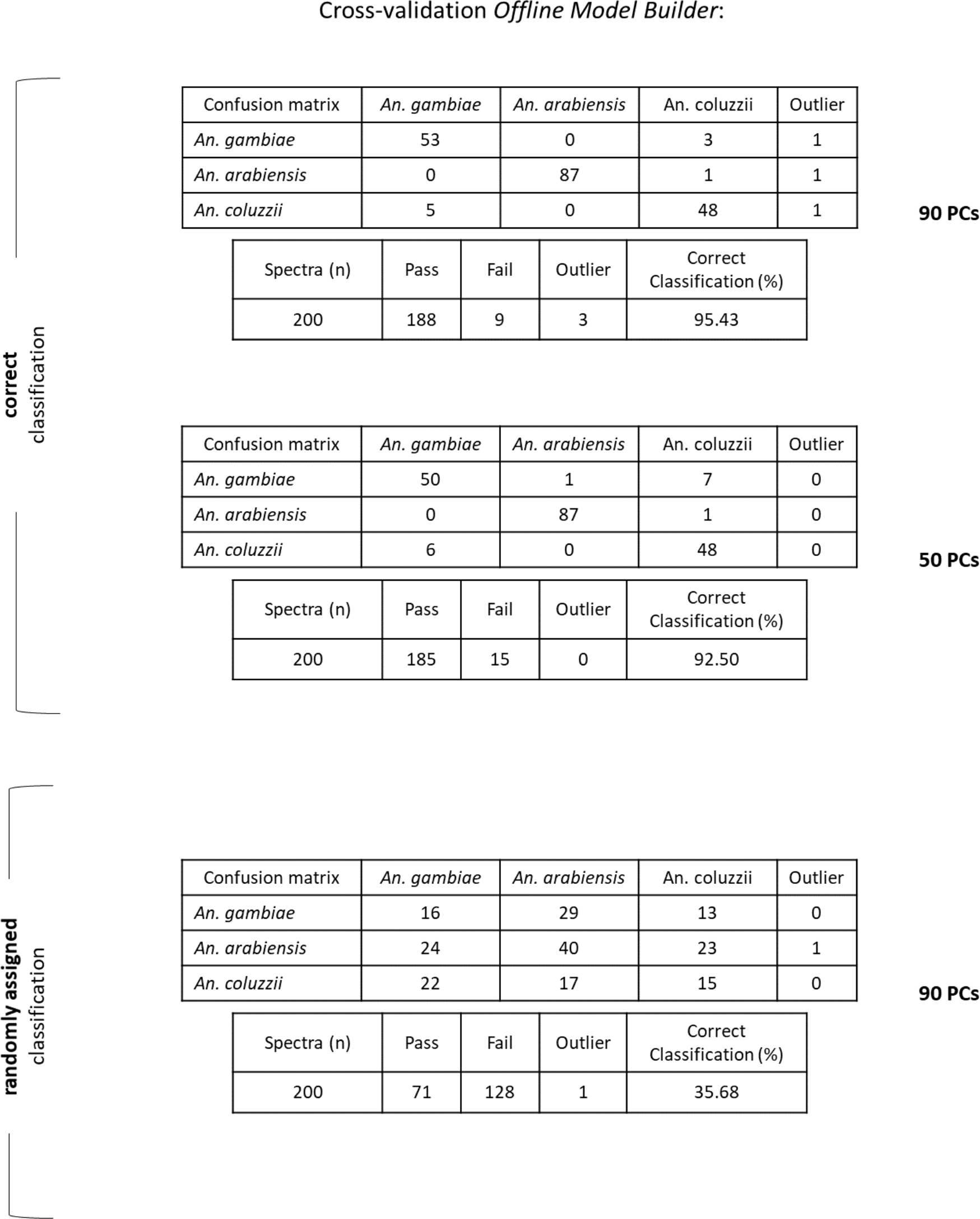
The PCA-LDA based species models (with correct classification, built with 90 and 50 PCs; with randomly assigned classes, built with 90 PCs) were cross-validated within Offline Model Builder using the setting ‘Leave 20 % out’ and a standard deviation of 5. During cross-validations two samples of the species An. gambiae and/or An. arabiensis were not tested as 20 % of 202 results in a fractional number.

**Supplemental Figure 5:**
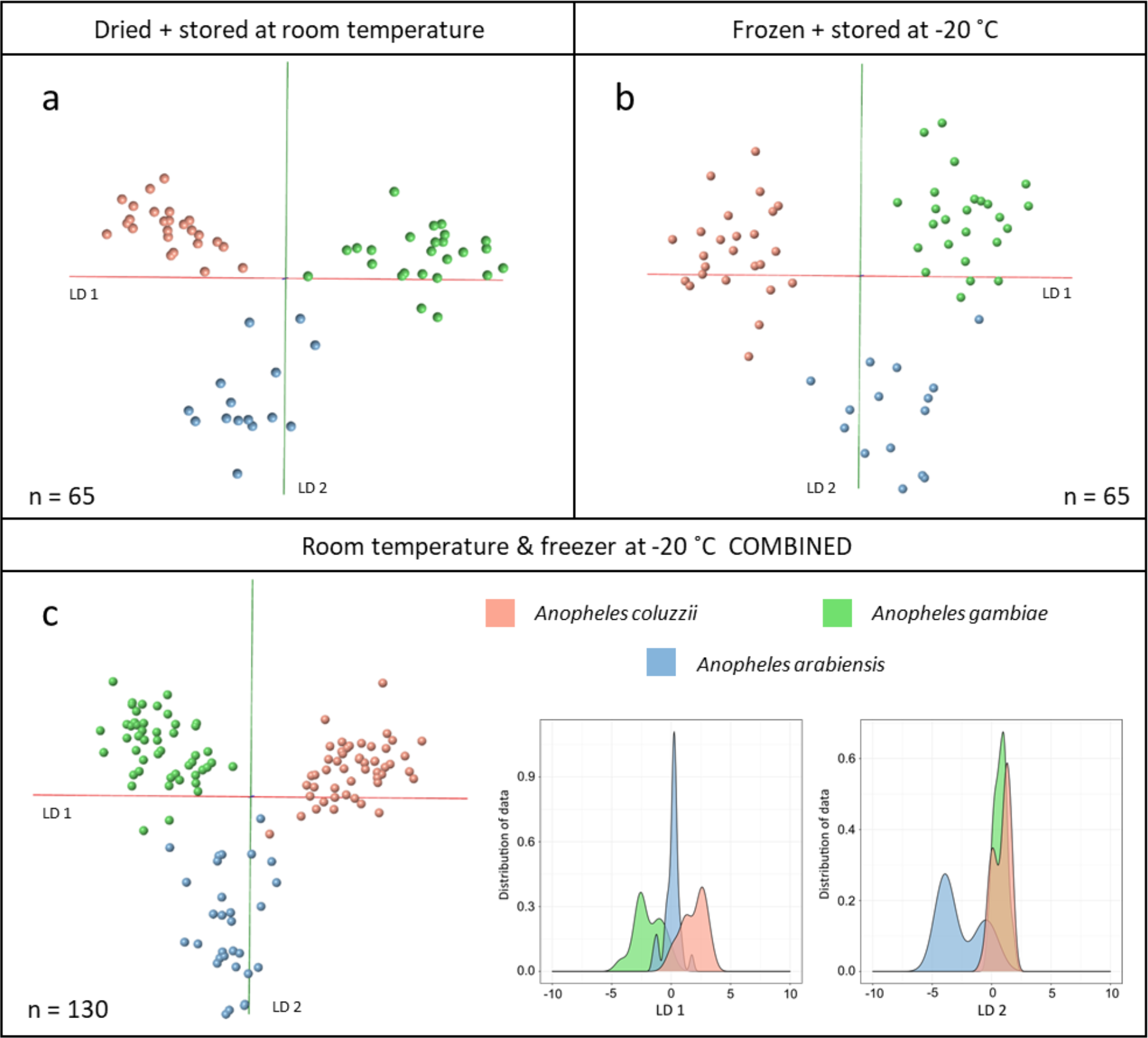
Female specimens from the three species An. arabiensis, An. gambiae and An. coluzzii were equally split into two groups: one group was killed through dehydration and stored at room temperature with desiccant material, the other group was killed by freezing and stored at -20°C in falcon tubes. Within each group samples were additionally split to be analysed at five different time points: immediately after killing (no storage) and after storage for 1, 2, 4 and 10 weeks. For every combination of storage type and length 5 An. coluzzii, 5 An. gambiae and 3 An. arabiensis mosquitoes were analysed. For both storage conditions (desiccated and frozen) samples from all storage time points were combined to build PC-LDA based species models in OMB (based on 35 and 30 PCs). Both storage types, dry at room temperature (panel a) and frozen at -20°C (panel b) led to informative REIMS spectra allowing differentiation of species through PC-LD analysis. Following successful separation in individual models, all samples were combined (n=130) for species classification (c). First, PC-LDA was attempted in Offline Model Builder (left) before exporting the data matrix and conducting the analysis in R (right); both were based on 70 PCs. Despite the large amount of variability in the sample set specimens were clustered into their respective species group.

**Supplemental Figure 6:**
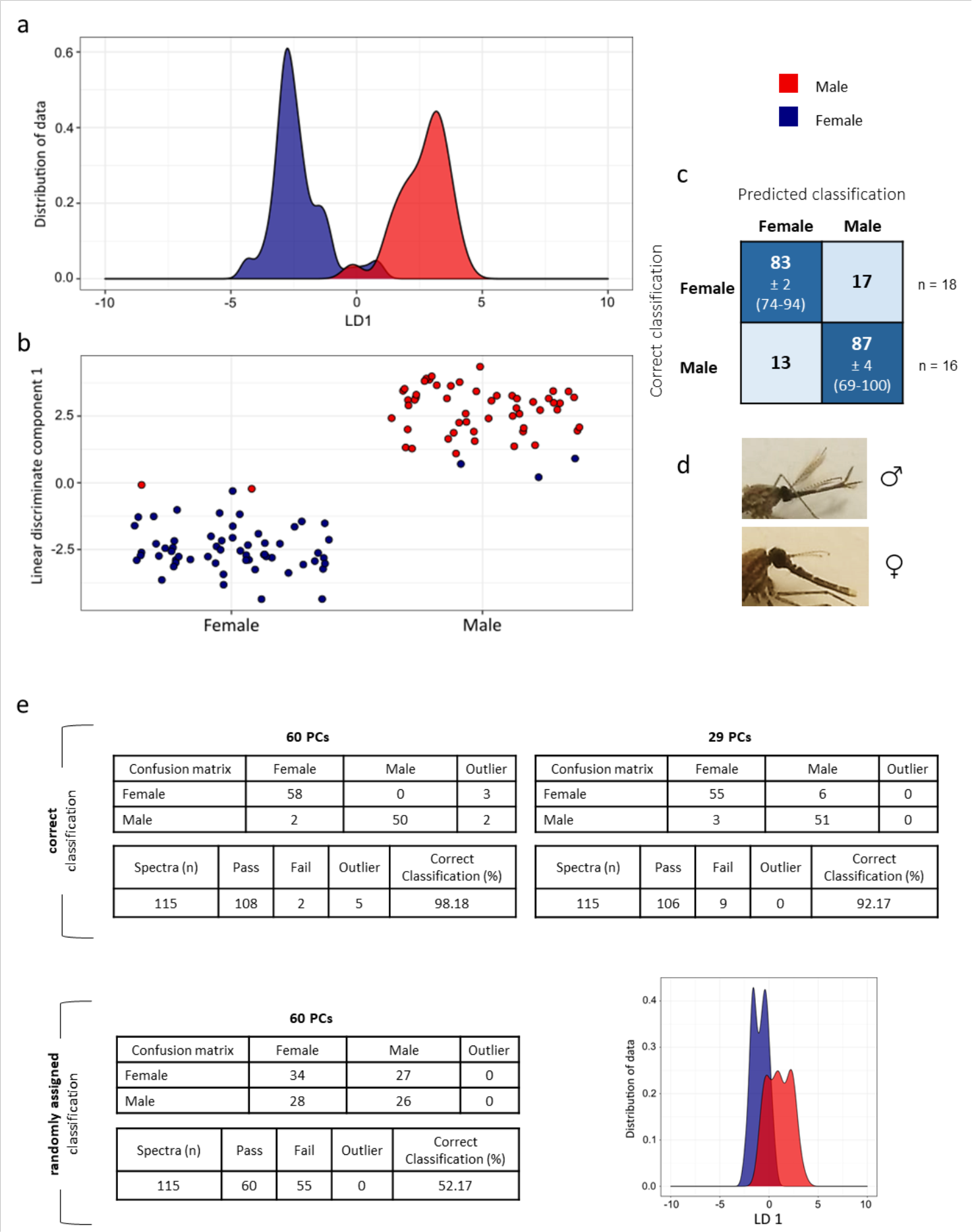
Laboratory raised Anopheles gambiae mosquitoes were separated into male (n=54) and female (n=61) classes, based on the sexual dimorphism of their antennae (d), before being analysed through REIMS. Data were analysed in the Offline Model Builder software (PC-LDA) as well as the R environment (PC-LDA, random forest). The results of PC-LDA (conducted in R, based on 60 PCs) are visualised in form of kernel density (a) and scatter plots (b), which both depict good separation of the sexes with only a small amount of samples (5) overlapping. Additionally, random forest analysis was conducted, using a 70 %/30 % split for model training and testing; the analysis was repeated 10 times with different samples in the training and testing category each time (randomly selected). The averaged results are listed as percentages in the confusion matrix (c) with SEM ± and the range of achieved accuracies (min-max) stated for the correct classification percentages. The average number of samples used for testing are listed at the end of the class rows (n=x). PC-LDA models, based on 60 PCs and 29 PCs, were cross-validated within OMB using the setting ‘Leave 20 % out’ and a standard deviation of 5 (e, upper panel). Furthermore, sample classifications (male, female) were randomly assigned to samples and the model was re-built using the initial principal component number (60 PCs) (e, bottom panel). The resulting separation is noticeably worse with nearly half of the male samples completely overlapping with the female class and a correct classification rate of only 52 %.

**Supplemental Figure 7:**
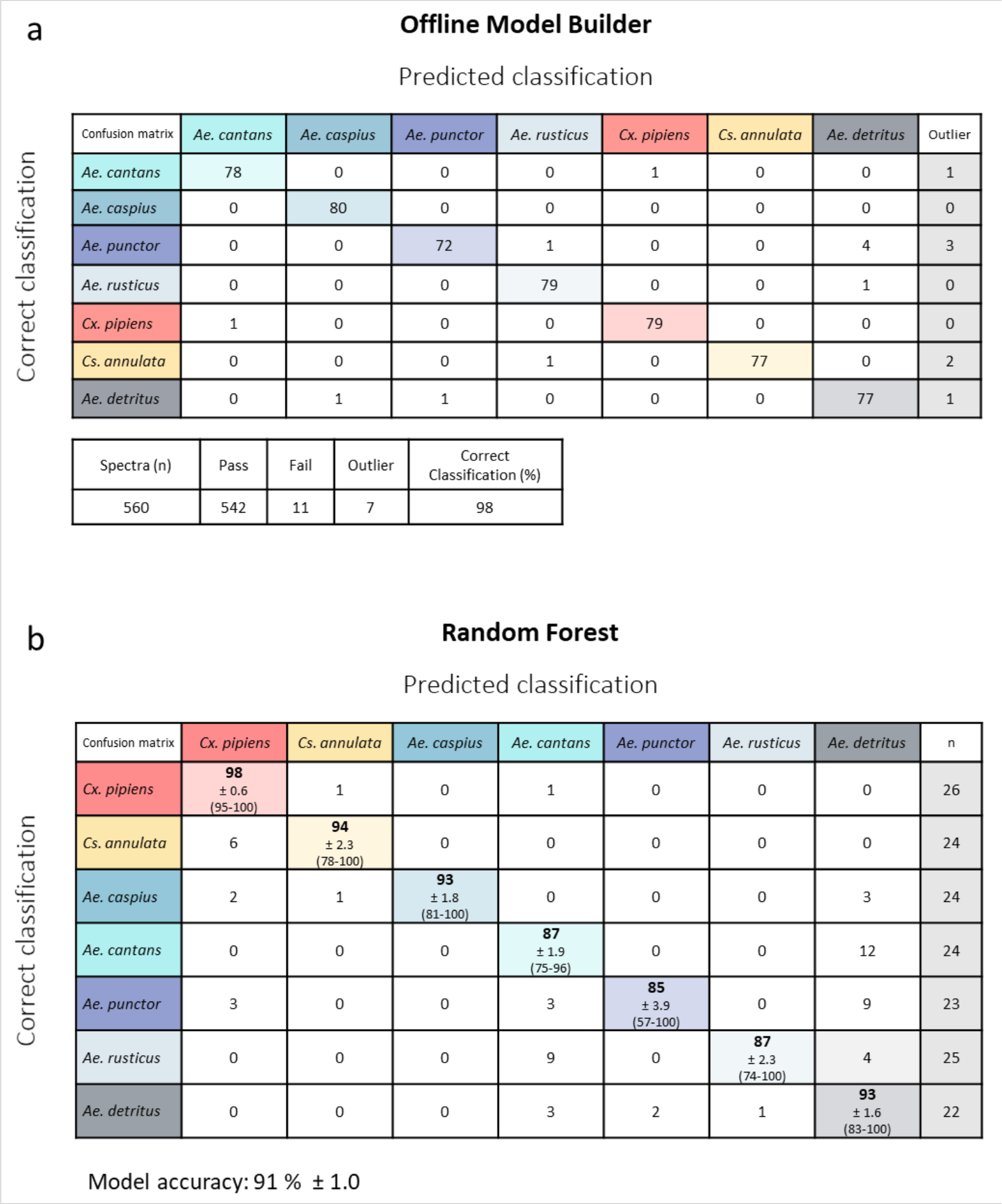
Cross-validation results for the seven species model built using 100 PCs (panel a). Cross-validation was performed within OMB using the option ‘Leave 20 % out’ and a standard deviation of 5. Results are listed in form of a confusion matrix containing the numbers of samples which have been either correctly or wrongly classified, as well as the number of outliers per classifications. The summary underneath contains the total number of spectra (samples) used for validation, the number of passed and failed samples, total number of outliers and the calculated correct classification rate (%) of the model. Random forest analysis of the seven species data set was repeated 10 times, using a different set of samples for model training (70 % of data) and testing (30 % of data) each time. The resulting confusion matrices, containing the numbers of correctly and wrongly classified samples, were turned into percentages and averaged over the 10 runs. The averaged correct classification accuracies (in %) plus SEM (±) and the range of achieved accuracies over 10 repeats (min and max) are listed in the coloured cells (panel b). The column on the right (n) states the average number of samples used for testing for each class. In total, the model achieved a classification accuracy of 91 %; meaning 91 out of 100 test samples would be identified correctly.

**Supplemental Figure 8:**
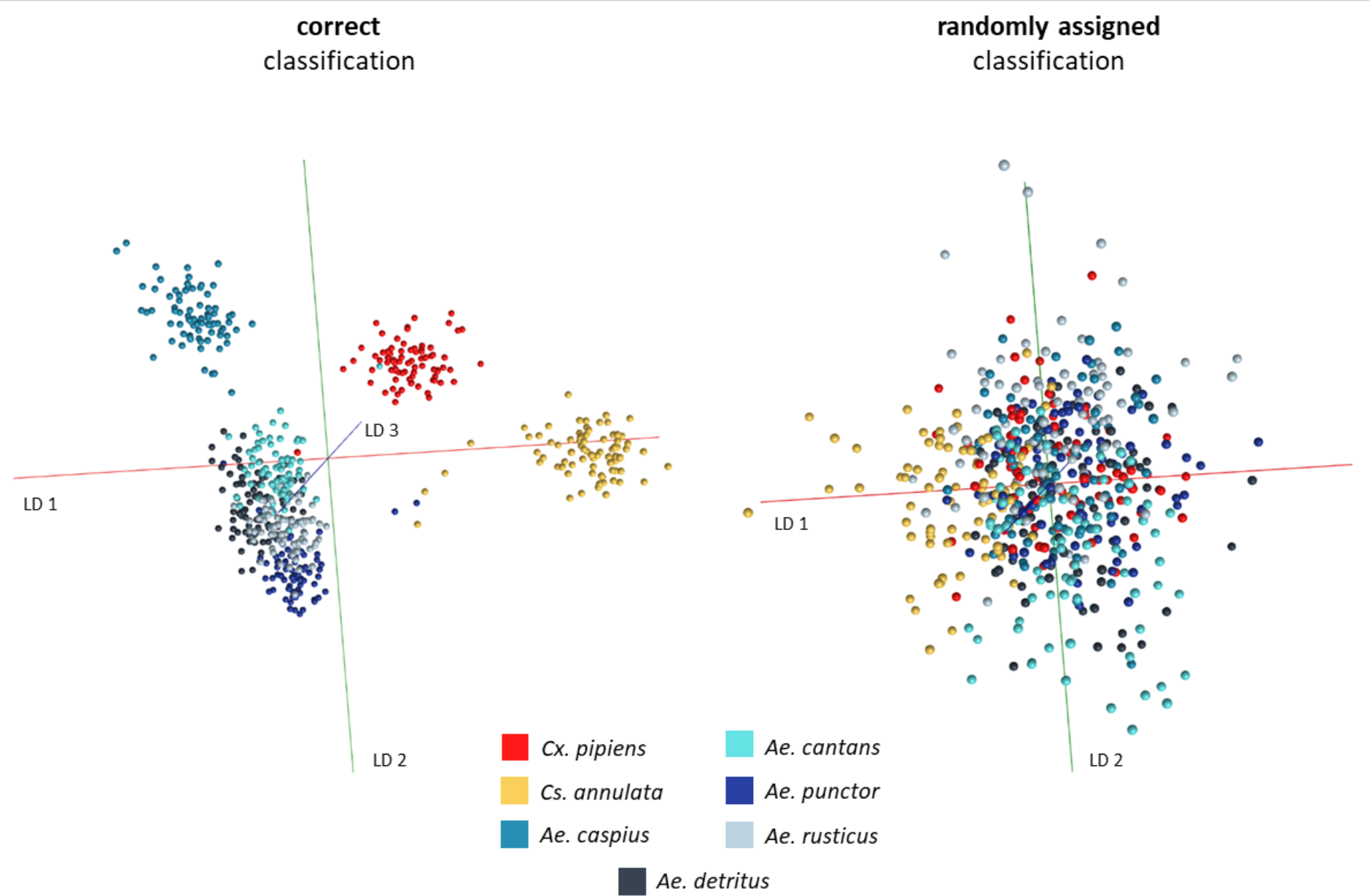
Comparison of the PC-LDA based 7-species model built with correct sample classifications (left) and randomly assigned classifications (right). Both models were built with the same settings in Offline Model Builder, using 100 principal components.

**Supplemental Figure 9:**
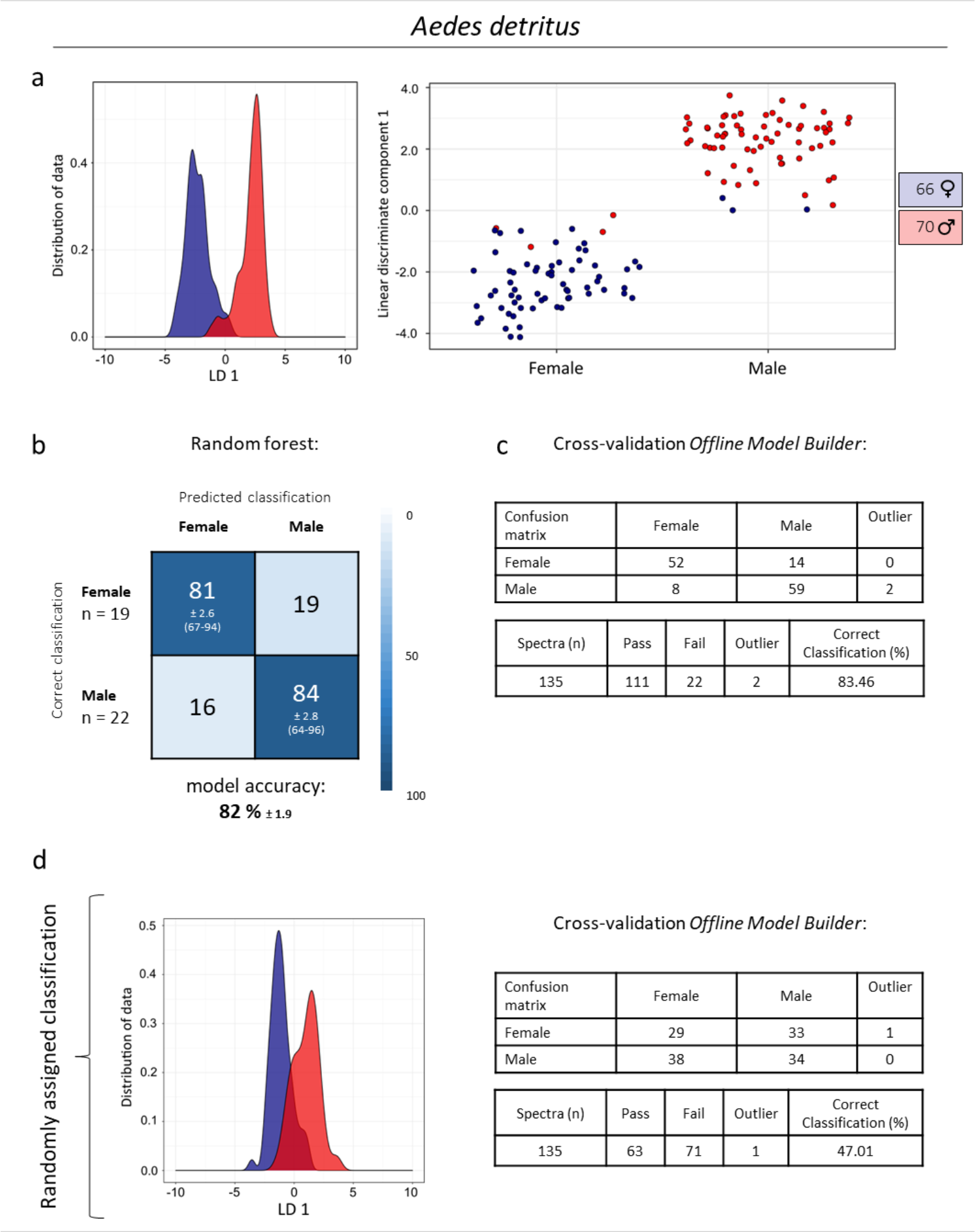
Separation of male (66) and female (70) Aedes detritus specimens based on principal component-linear discriminant analysis, displayed in form of a smoothed histogram (a, left) and a scatterplot (a, right) using 80 principal components. The separation was further examined through cross-validation (c) in OMB (‘Leave out 20%’, standard deviation of 5) as well as random forest analysis (b) in R (70 % training/30 % testing, repeated 10 times). The cross-validated OMB model was based on 70 PCs, one sample was left out during validation as 20 % of 136 samples results in a fractional number that was rounded to the nearest integer. The results of the random forest analysis are depicted in a confusion matrix containing information about the percentages of samples, which had been either correctly or wrongly classified, including the standard error of the mean (±) and the range of accuracies achieved (min and max) for the correct classifications. The average number of samples (n) tested from each class is listed on the left-hand side of the table and the average model accuracy (plus SEM) underneath. Additionally, the PC-LDA based model was re-built with randomly assigned classifications (d, left) and again tested via cross-validation in OMB (d, right), which resulted in the expected decrease of classification accuracy and confirmed that separation (with correct classifications) is based on sex related variance.

**Supplemental Figure 10:**
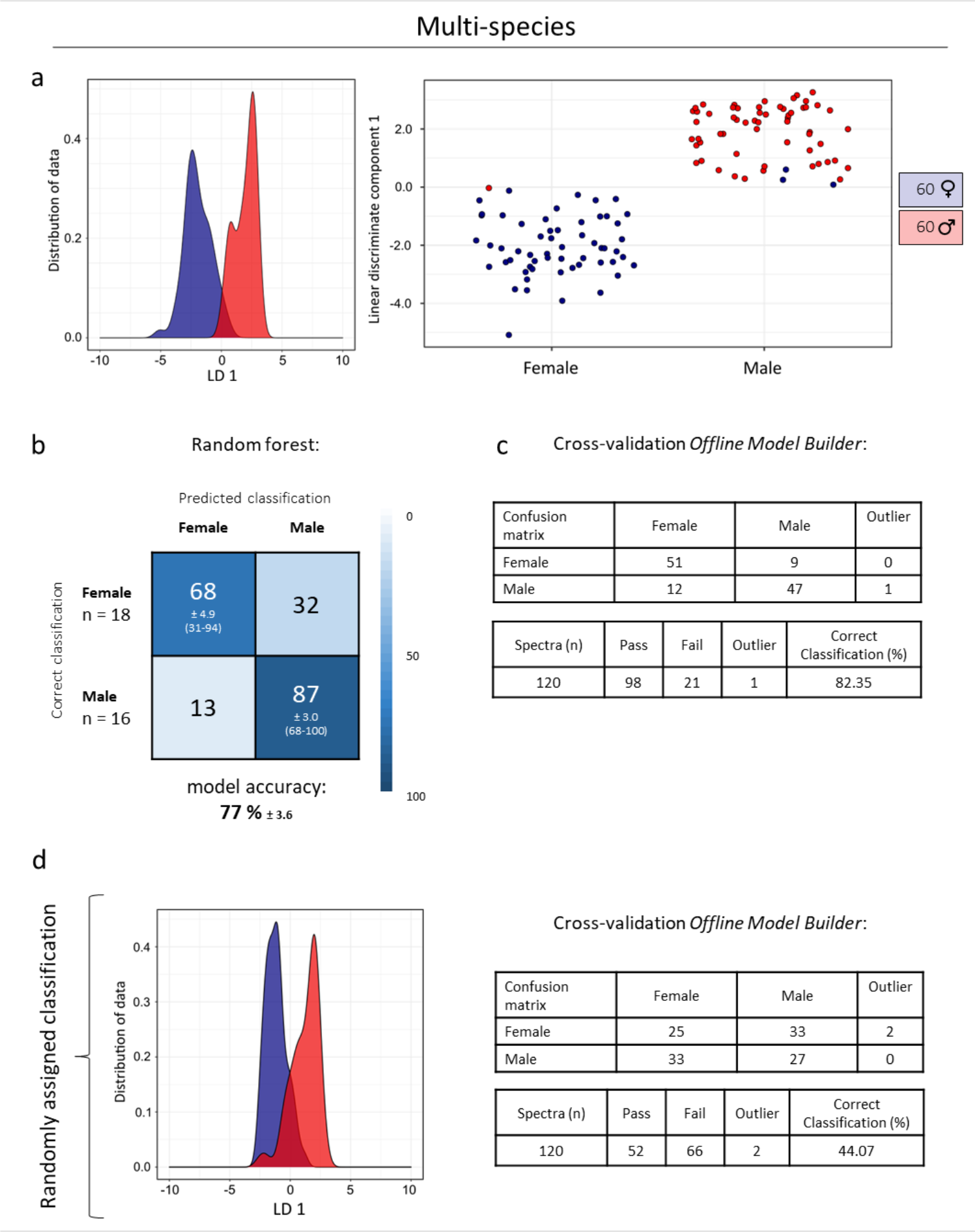
Male (60) and female (60) specimens were equally selected from four different mosquito species (Aedes detritus, Aedes punctor, Aedes rusticus, Aedes cantans) to test for species independent separation of sexes. The separation is based on principal component-linear discriminant analysis (70 principal components) and displayed in form of a smoothed histogram (a, left) and a scatterplot (a, right). The separation was further examined through cross-validation (c) in OMB (‘Leave out 20%’, standard deviation of 5) as well as random forest analysis (b) in R (70 % training/30 % testing, repeated 10 times). The cross-validated OMB model was also based on 70 PCs. The results of the random forest analysis are depicted in a confusion matrix containing information about the percentages of samples, which had been either correctly or wrongly classified, including the standard error of the mean (±) and the range of accuracies achieved (min and max) for the correct classifications. The average number of samples (n) tested from each class is listed on the left-hand side of the table and the average model accuracy (plus SEM) underneath. Additionally, the PC-LDA based model was re-built with randomly assigned classifications (d, left) and again tested via cross-validation in OMB (d, right), which confirmed that separation (with correct classifications) is based on sex related variance, despite including multiple species.

**Supplemental Figure 11:**
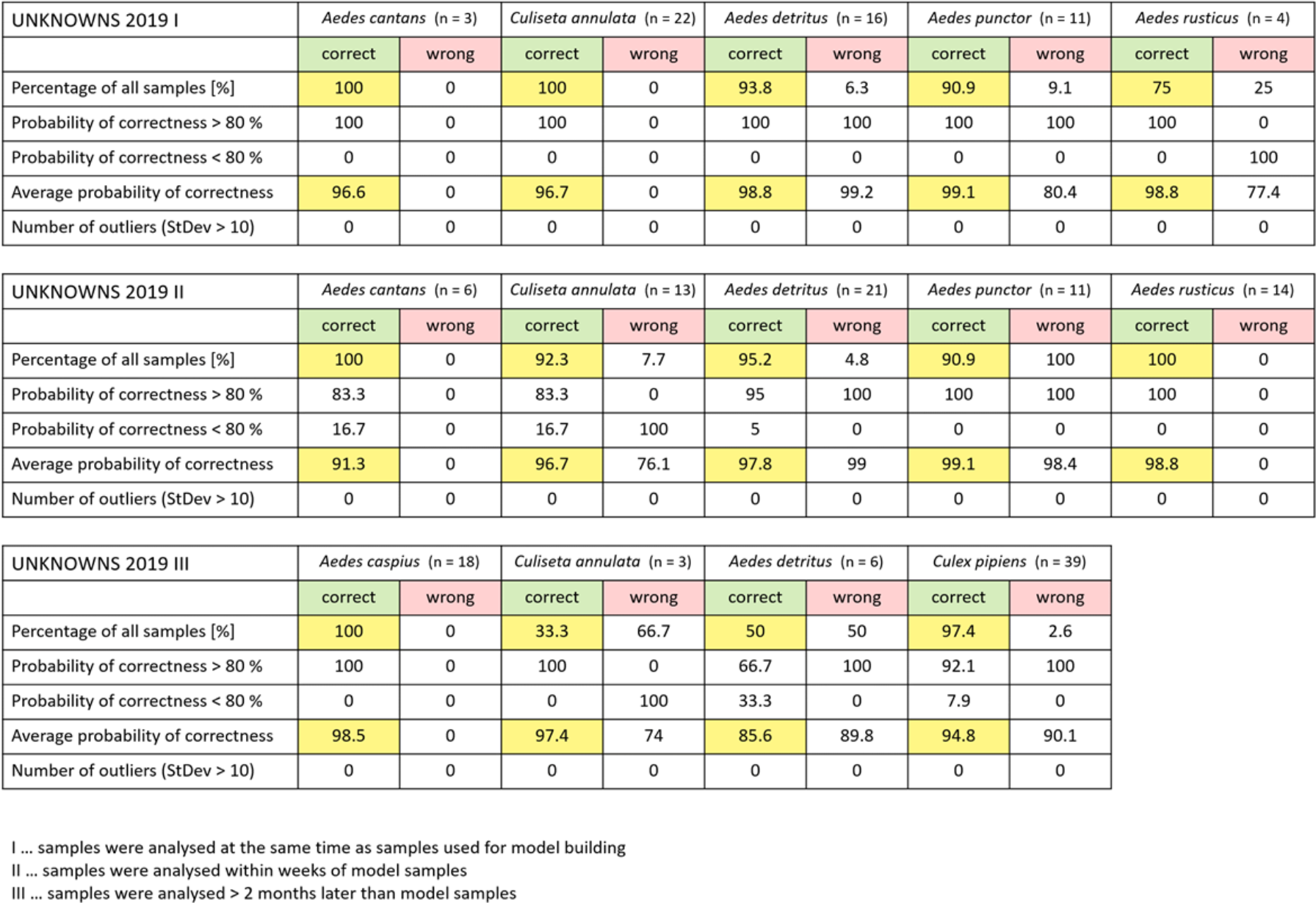
Identifications results of samples (raised) analysed in the same year as samples used for model building, listed for each species. The percentage of correctly identified samples and the probability that the identification is correct are highlighted in yellow for easier comparison.

**Supplemental Figure 12:**
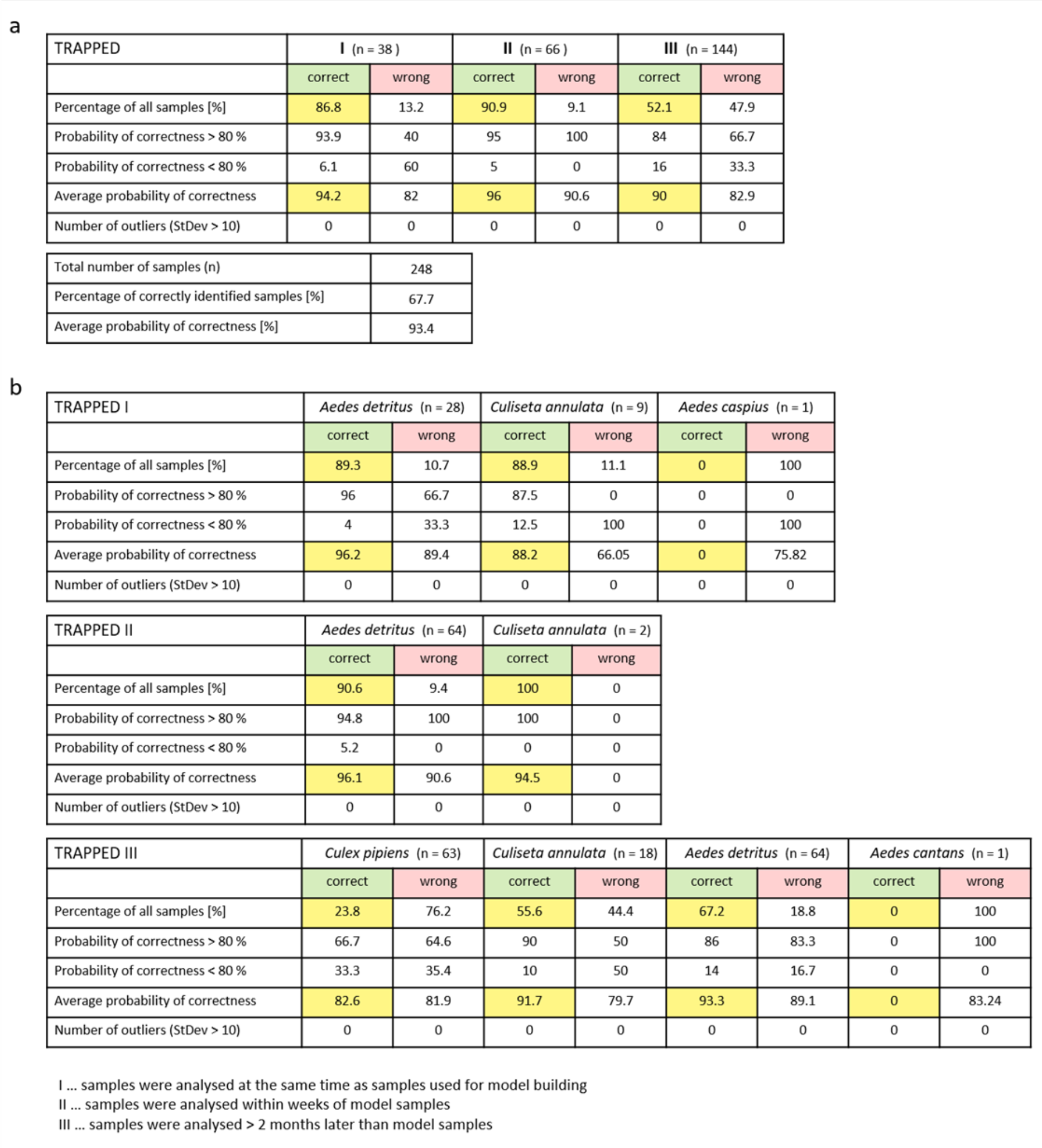
The seven species PC-LDA model built in Offline Model Builder with 100 PCs and exported to the Recognition software, was used to identify wild mosquitoes caught in traps (TRAPPED). Samples were categorised depending on their time point of analysis : samples which had been analysed at the same time as samples used for model building (I), samples analysed on other days than model samples, but same time period (II) and samples which were analysed over two months later than the last samples included in the model building (a). The number of tested samples is stated in brackets. The percentage of correctly identified samples and the likelihood that the identification is correct are highlighted in yellow for easier comparison. A more detailed view of the identification results and number of samples per species is listed under b.

**Supplemental Figure 13:**
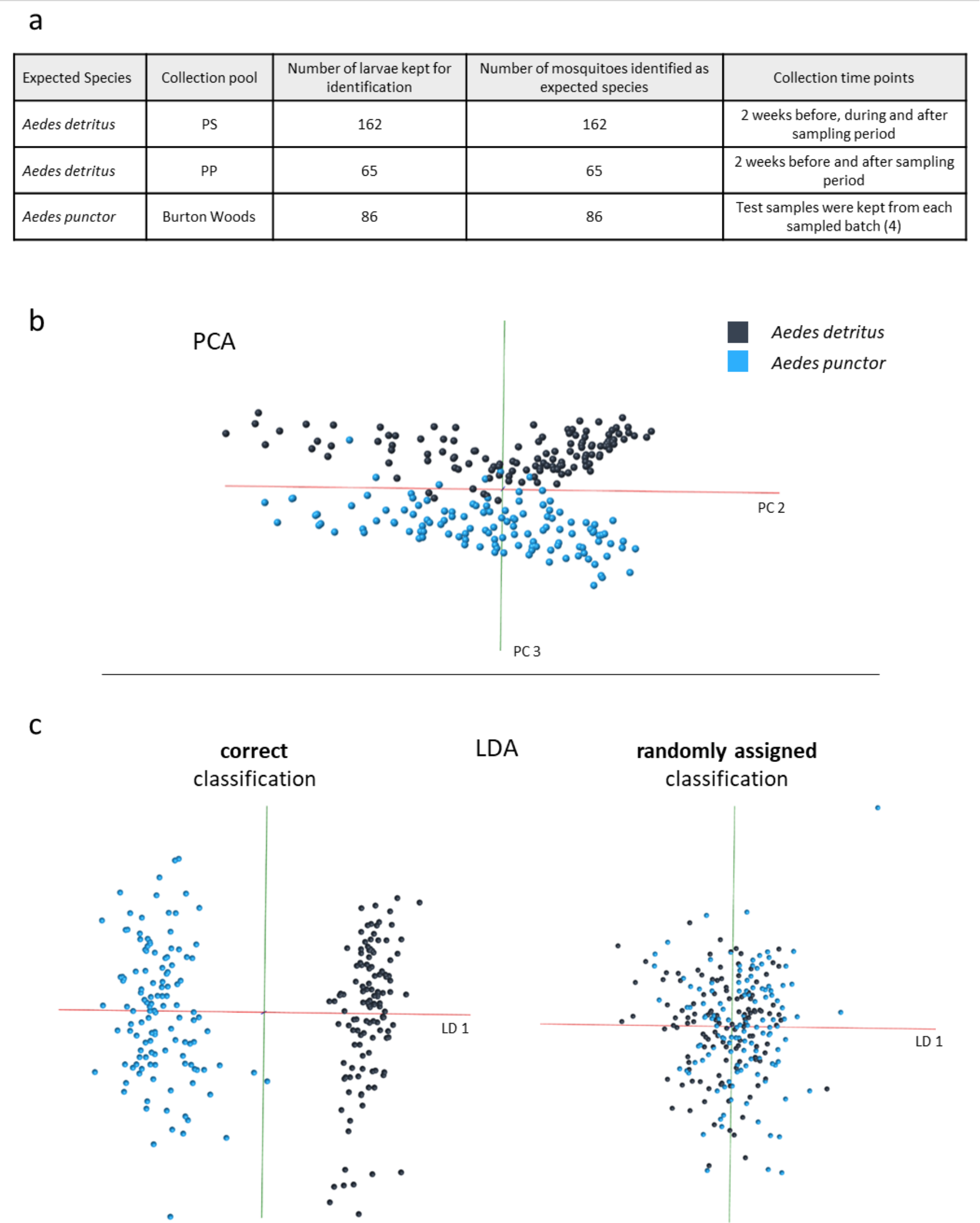
Larvae were collected from three pools for REIMS analysis as well as identification purposes. For species identification larvae were raised to adults; all larvae emerged as the expected species (a). The difference between the larvae of Aedes detritus and Aedes punctor is adequate to provide separation even when using unsupervised methods such as PCA. The individual differences are represented by the principal components 1 and 2. The variance in component 3, however, supports a clear clustering of samples into their respective species (b). To test separation the PC-LDA model was also built with classifications randomly assigned to samples. A comparison of the larval species model with and without correctly assigned classes can be seen in panel c.

**Supplemental Figure 14:**
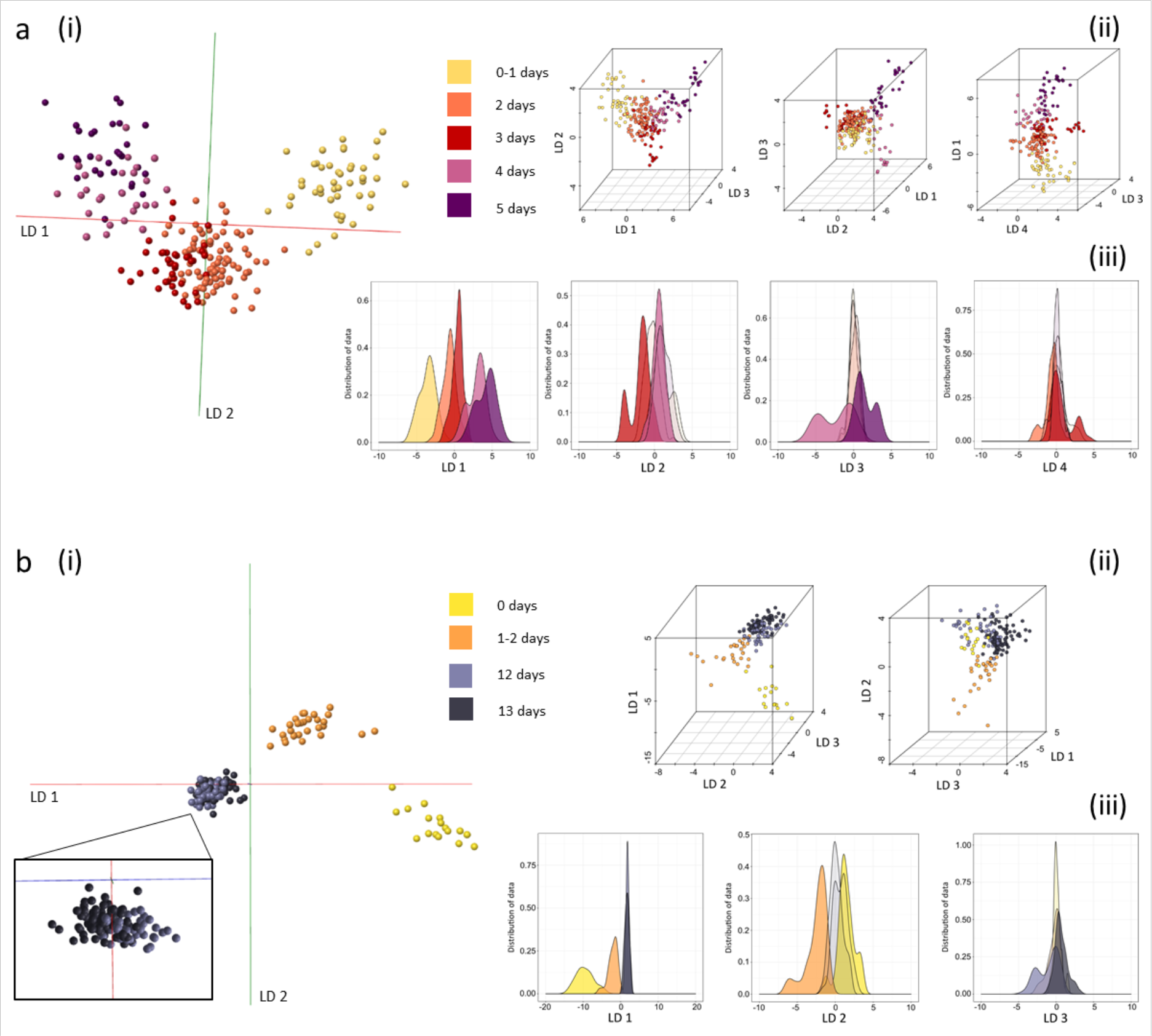
Age groups were separated by PC-LD analysis and visualized using OMB (i) as well as R , in form of 3D models (using different linear discriminant combinations) (ii) and kernel density plots for each LD) using only a quarter of principal components possible. The difference between classes in model a (based on 56 PCs) decreased with the lower PC number. This is especially noticeable between groups 2 and 3, which now strongly overlap and groups 4 and 5, where samples are clustered only loosely without clear group boundaries. The young groups in model b (based on 44 PCs), moved closer to each other due to the reduction in PC numbers, however, separation is still very distinct. The main portion of the older sample classes (12+13 days) are now completely overlaid.

**Supplemental Figure 15:**
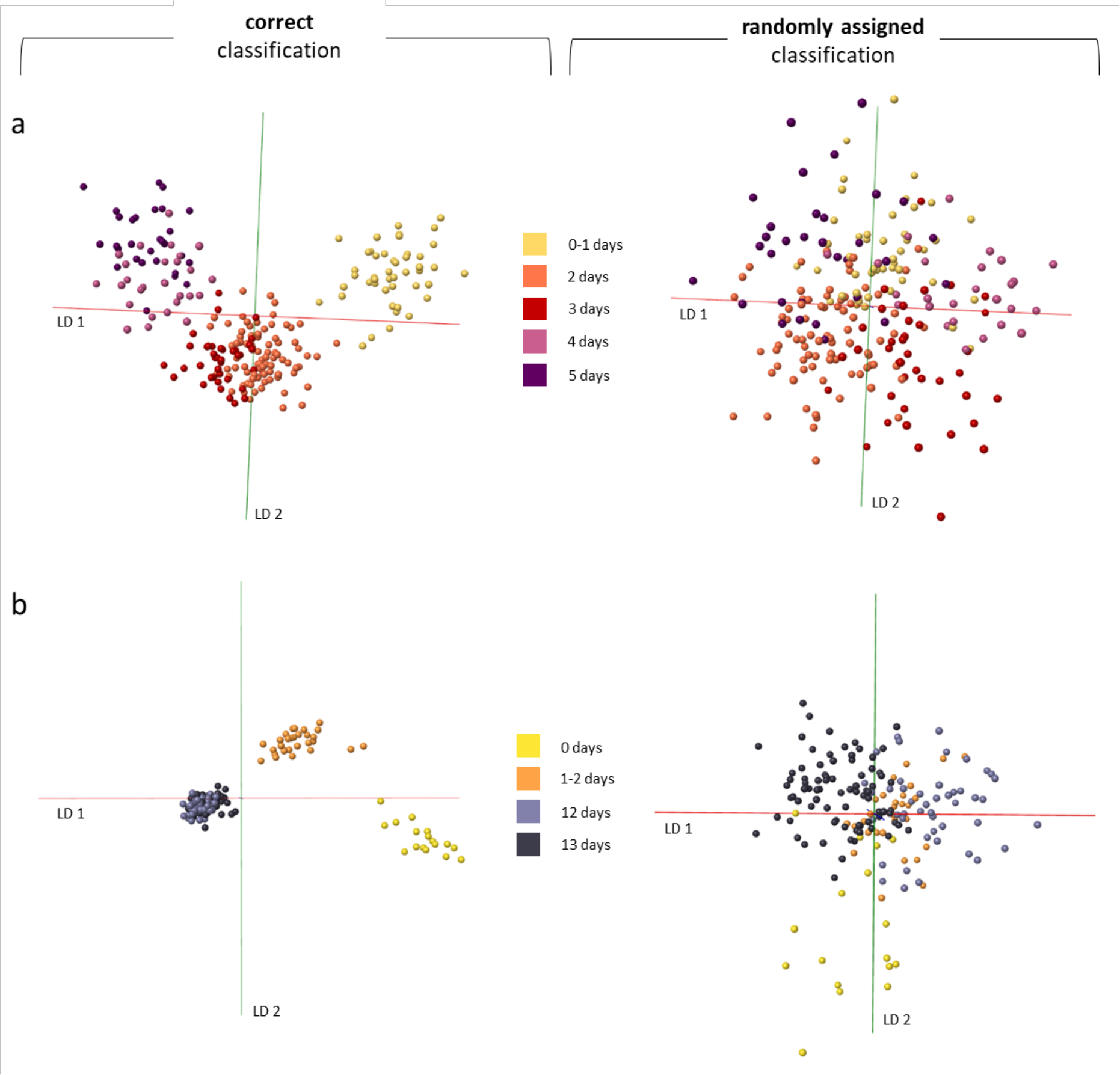
To test the separation principle of model a (based on 100 PCs) and model b (based on 88 PCs) in Figure 6, classifications were randomly assigned to samples before rebuilding the models in Offline Model Builder. The original separations (left panel) can be directly compared to the randomly assigned classification models (right panel). For both models the separation following the randomisation is significantly worse with samples from the same class clustering only very loosely compared to previous grouping and significant overlap of groups.

**Supplemental Figure 16:**
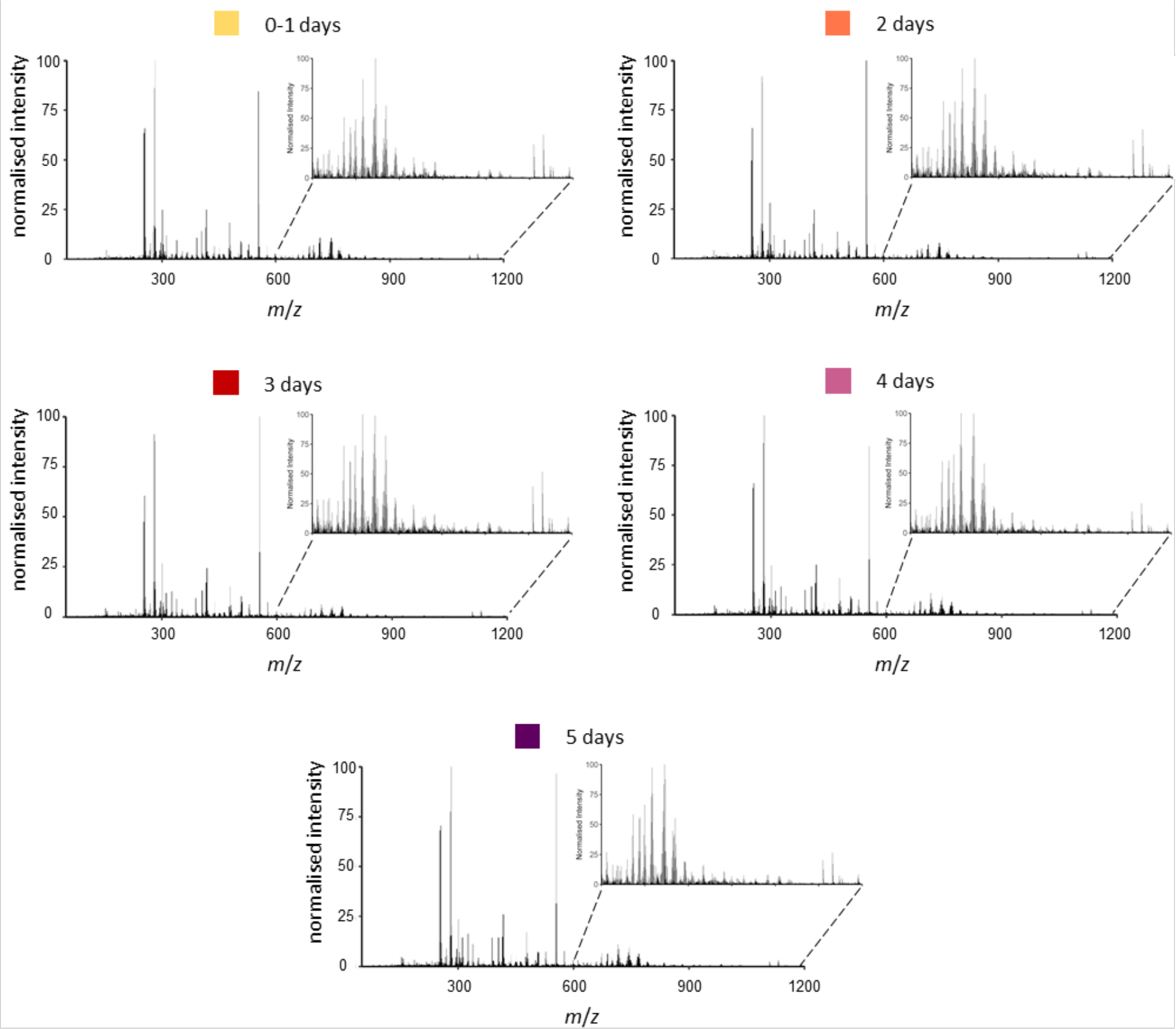
The data matrix, obtained after processing and binning the mass spectral data in Offline Model Builder, was used to create averaged mass spectra for all age classes from 0-5 days. Each mass spectrum represents an average of all samples available for each age group: 0-1 day (n=47), 2 days (n=84), 3 days (n=39), 4 days (n=27), 5 days (n=30).

**Supplemental Figure 17:**
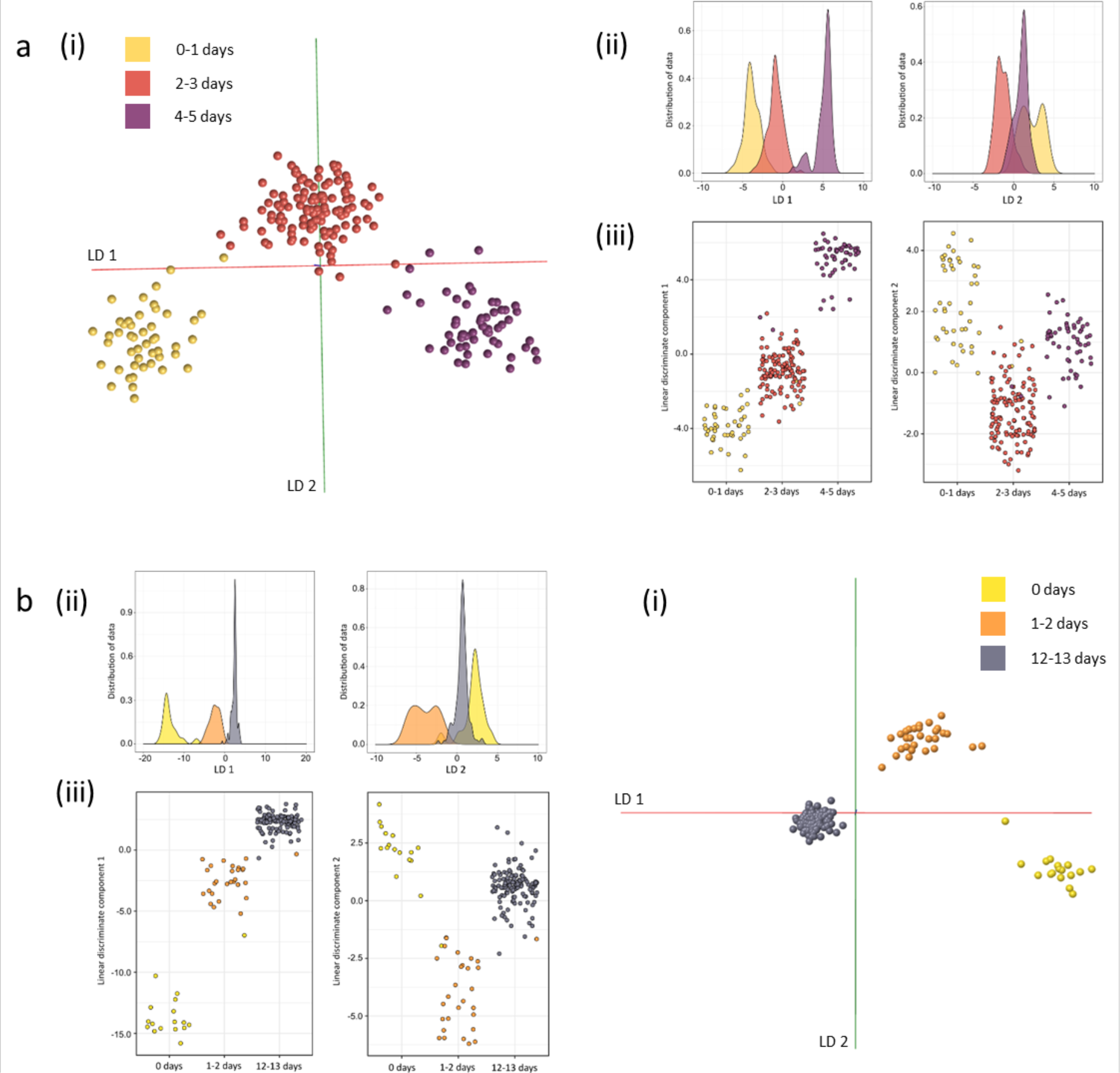
To improve separation of the individual age classes for both models some age groups were combined into one class. For the model in panel a the 2 and 3 day old mosquitoes, as well as the 4 and 5 day old specimens, were combined into one group each, reducing the overall number of classes from 5 to 3 (panel a). As mosquitoes which have just emerged and 1 day old mosquitoes can be readily distinguished, only the 12 and 13 day old mosquitoes were combined into one group for the model in panel b. As with the previous age models, PC-LD analysis was conducted first in Offline Model Builder (i) to extract the data matrix, before repeating analysis in R to visualise separation results through kernel density histograms (ii) and 2D scatter plots (iii). Principal component numbers were the same as used for the previous model (model a: 100 PCs, model b: 88 and 85 PCs) to solely observe the effect of class reduction. For both models all age groups are now separated along linear discriminant one; LD 2 merely contributes additional variance to increase separation of the younger groups. There are now distinct gaps between all age classes.

**Supplemental Figure 18:**
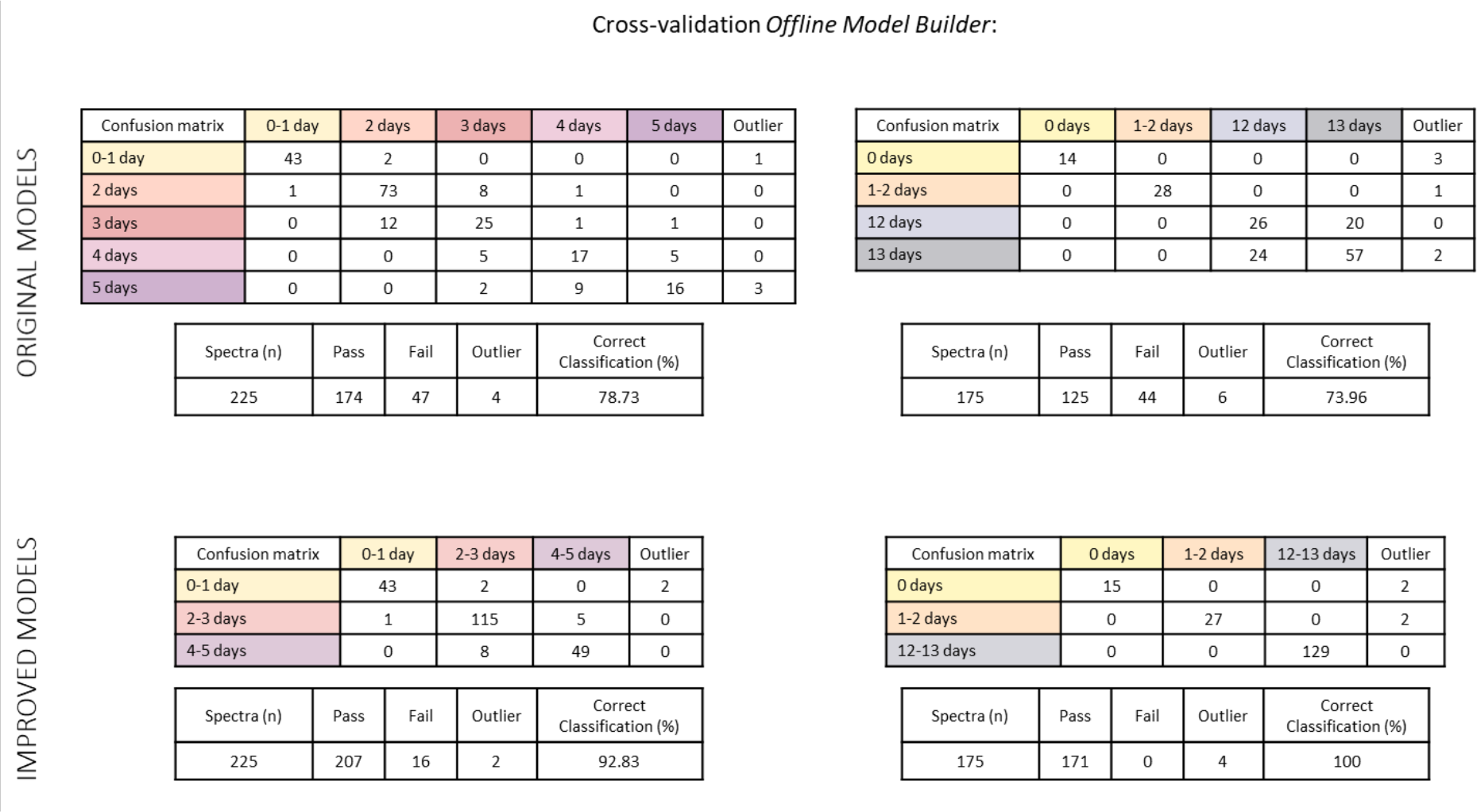
The PCA-LDA based age models, one comprising 5 classes (0-5 days) and the other 4 classes (0-13 days), were cross-validated within Offline Model Builder using the setting ‘Leave 20 % out’ and a standard deviation of 5 (top panel, original models)). The models comprised of combined age classes were also cross-validated using the same settings (bottom panel, improved models). Combining the age classes clearly improved separation accuracy from 79 to 93 % (for the model on the left) and 74 to 100 % (for the model on the right). Some samples were not tested as 20 % of the total sample number resulted in a fractional number.

**Supplemental Figure 19:**
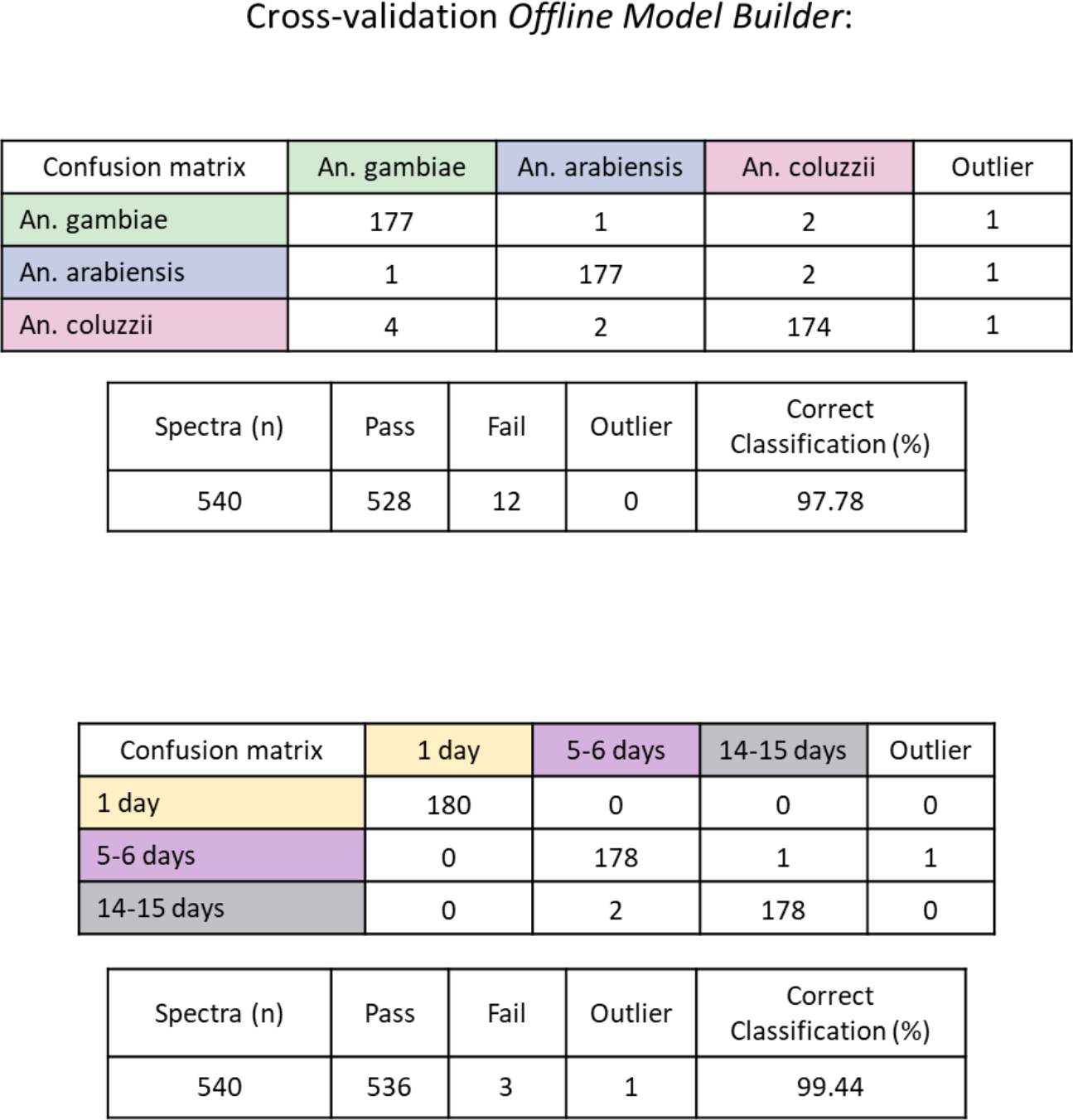
The Anopheles species model and age model (both based on 100 PCs) were cross-validated within Offline Model Builder using the setting ‘Leave 20 % out’ and a standard deviation of 5.

**Supplemental Figure 20:**
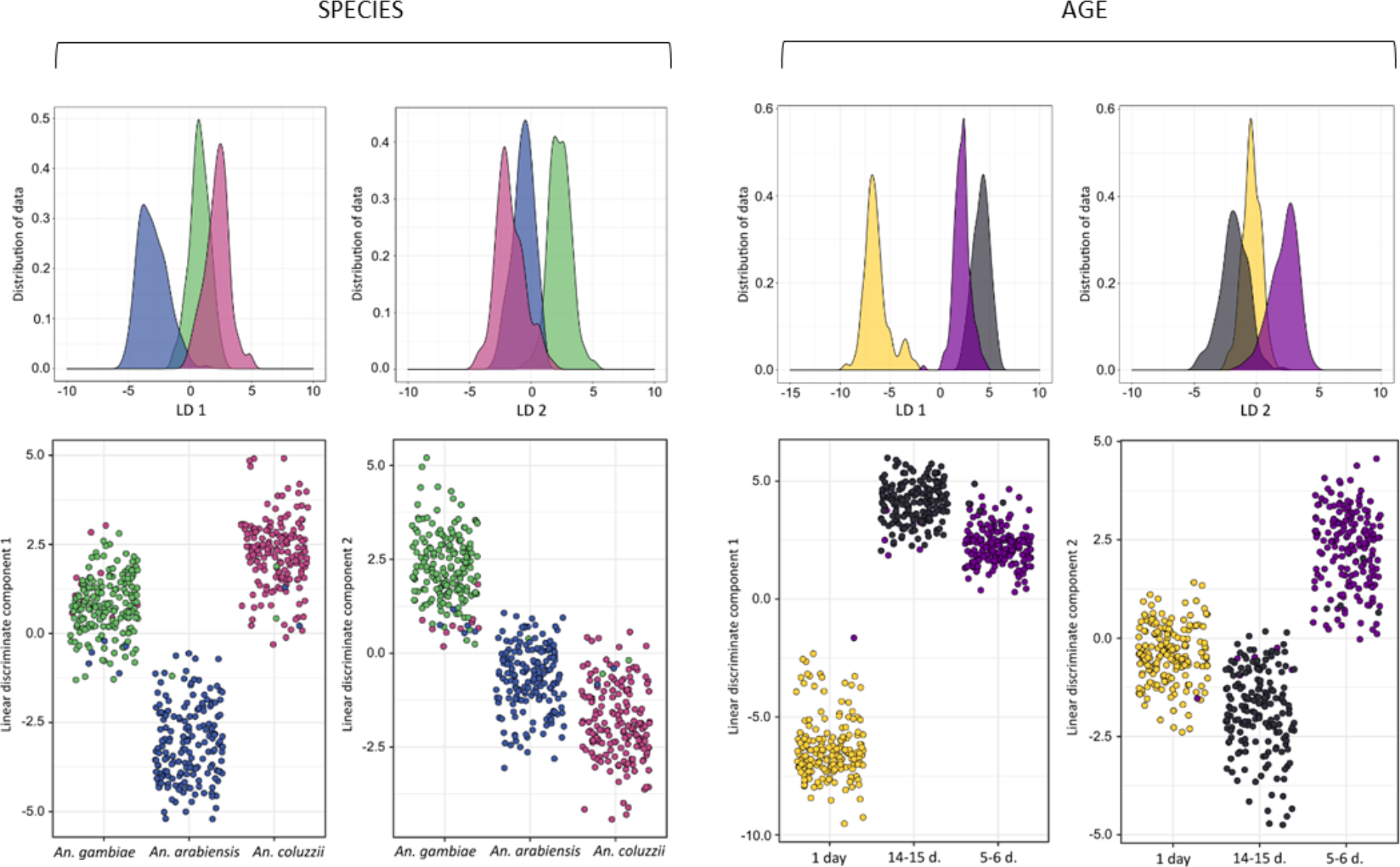
The PCA-LDA models separating Anopheles mosquitoes by species and age were re-built in R using a lower number of principal components. The separation depicted in the kernel density histograms and scatter plots is based on 135 PCs (¼ of max) for both models.

**Supplemental Figure 21:**
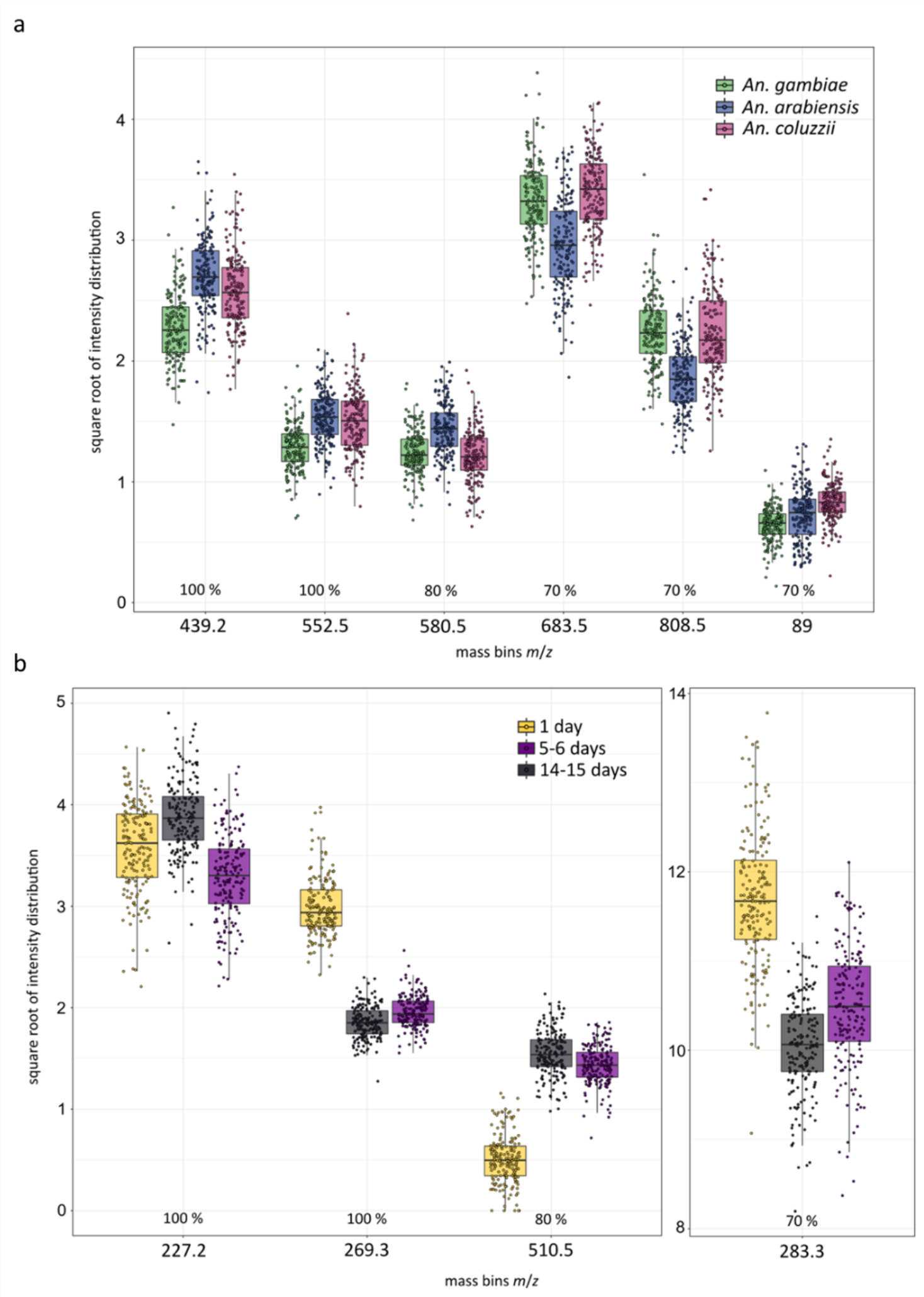
A list of the variables identified as important for the random forest based separation of the three mosquito species An. gambiae, An. arabiensis and An. coluzzii (a) as well as the three age groups 1 day, 5-6 days and 14-15 days (b). Only the intensities of ion bins, which have been in the Top 10 variables list in at least 7 out of 10 random forest runs, are plotted. Although some of the ion bins had not been identified as very important in every run, they nevertheless play an important role in the separation process. The m/z bins 580.5, 683.5 and 808.5 appear to support the separation of Moz from the other two classes, which was not achieved with the variables identified in every run (100 %). The ion bins which are driving age separation 100 % of the time, however, already provide enough variance to separate all three groups; the other two bins (510.5 and 283.3) merely add further variance to the process.

**Supplemental Figure 22:**
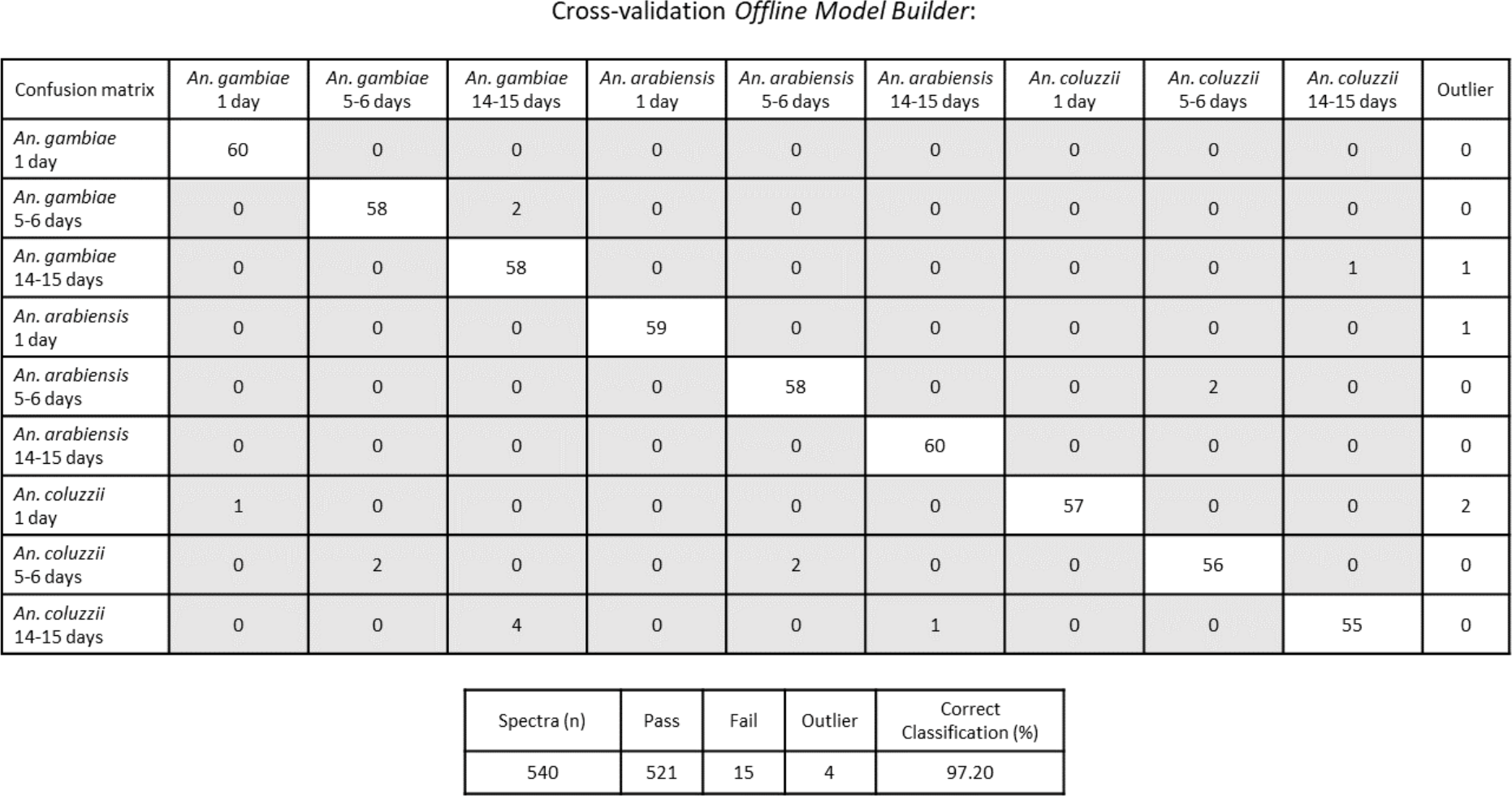
The nine-class species/age model (LDA based on 100 PCs) was cross-validated within Offline Model Builder using the setting ‘Leave 20 % out’ and a standard deviation of 5.

**Supplemental Figure 23:**
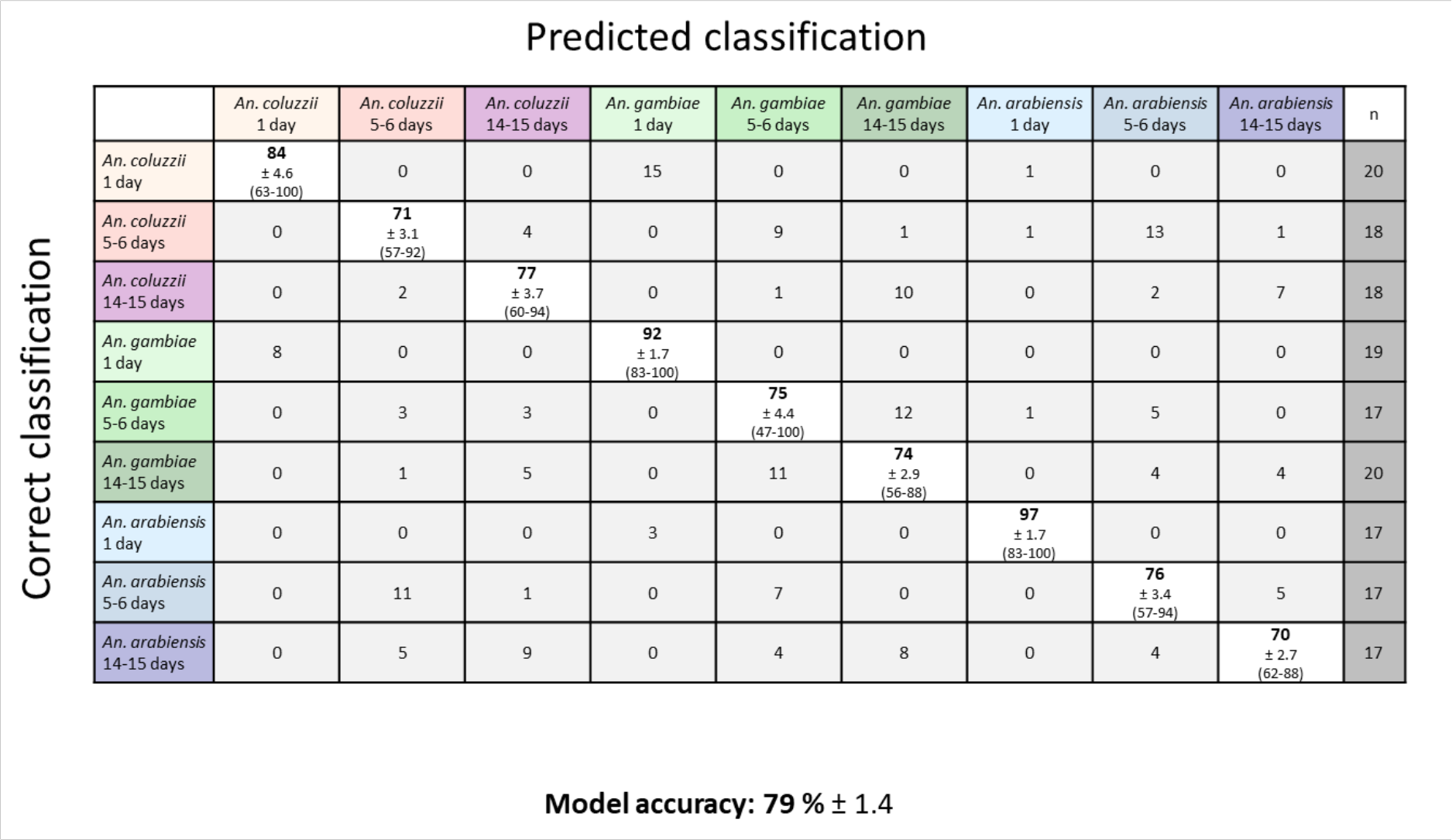
The data matrix from the nine-class species/age model was used for random forest analysis, which was repeated 10 times, using different randomly selected training (70 % of the data) and test (30 % of the data) data sets. The confusion matrix contains the mean percentages of correctly identified and misidentified samples for every species as well as the standard error of the mean. The range of classification accuracy achieved for each of the 10 models (lowest and highest percentage) is listed in parentheses below the standard error of the mean. The average number of samples per class used for testing the model are listed on the side (n = x). The overall model accuracy was 79 ± 1.4 % (mean ± SEM).

**Supplemental Figure 24:**
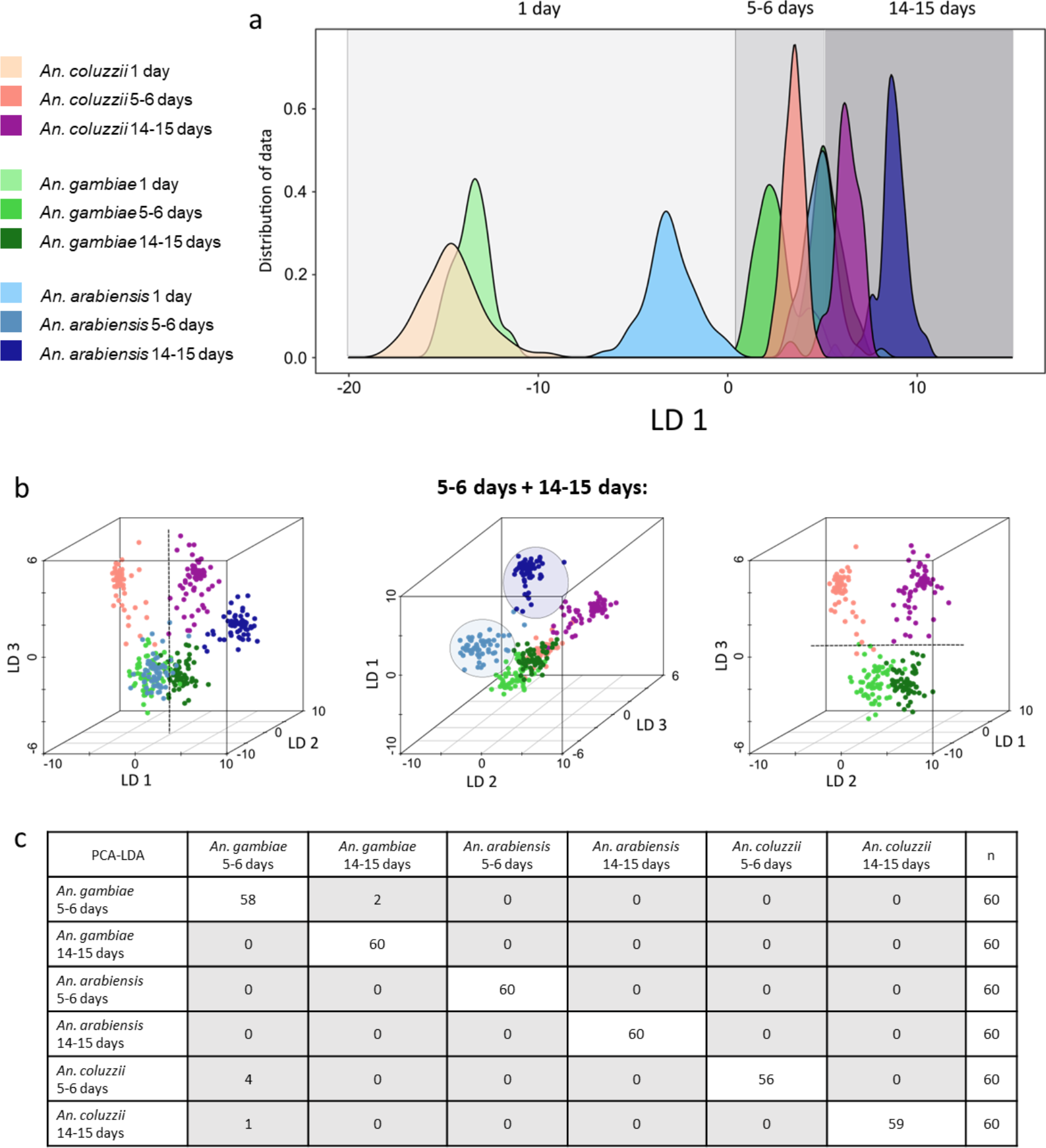
After building the nine-class model in Offline Model Builder, the data matrix was exported to repeat PC-LD analysis in R. The kernel density plot (LDA was based on 235 PCs) demonstrates an age related separation along LD1 (a), quite similar to what was observed in the Offline Model Builder result. Again, to simplify visualisation, the 1 day old mosquitoes were removed from the data set and the PC-LDA (180 PCs) separation process further examined using 3D scatter plots (b). While having a similar distribution of variance across the linear discriminants as seen in the Offline Model Builder model, the separation seems less defined and the class ‘An. arabiensis 14-15 days’ is now separated along LD1, together with the age clusters, instead of LD 2. Nevertheless, separation of groups due to age seems to happen along LD1, An. arabiensis is separated along LD1 and LD 2 and to distinguish An. coluzzii and An. gambiae groups LD 3 is needed. Interestingly, separation of differently aged mosquitoes is easier with An. coluzzii than with An. gambiae specimens. Plotting the outcomes of PC-LD analysis as a matrix table reveals that the six classes are very well separated (c).

**Supplemental Figure 25:**
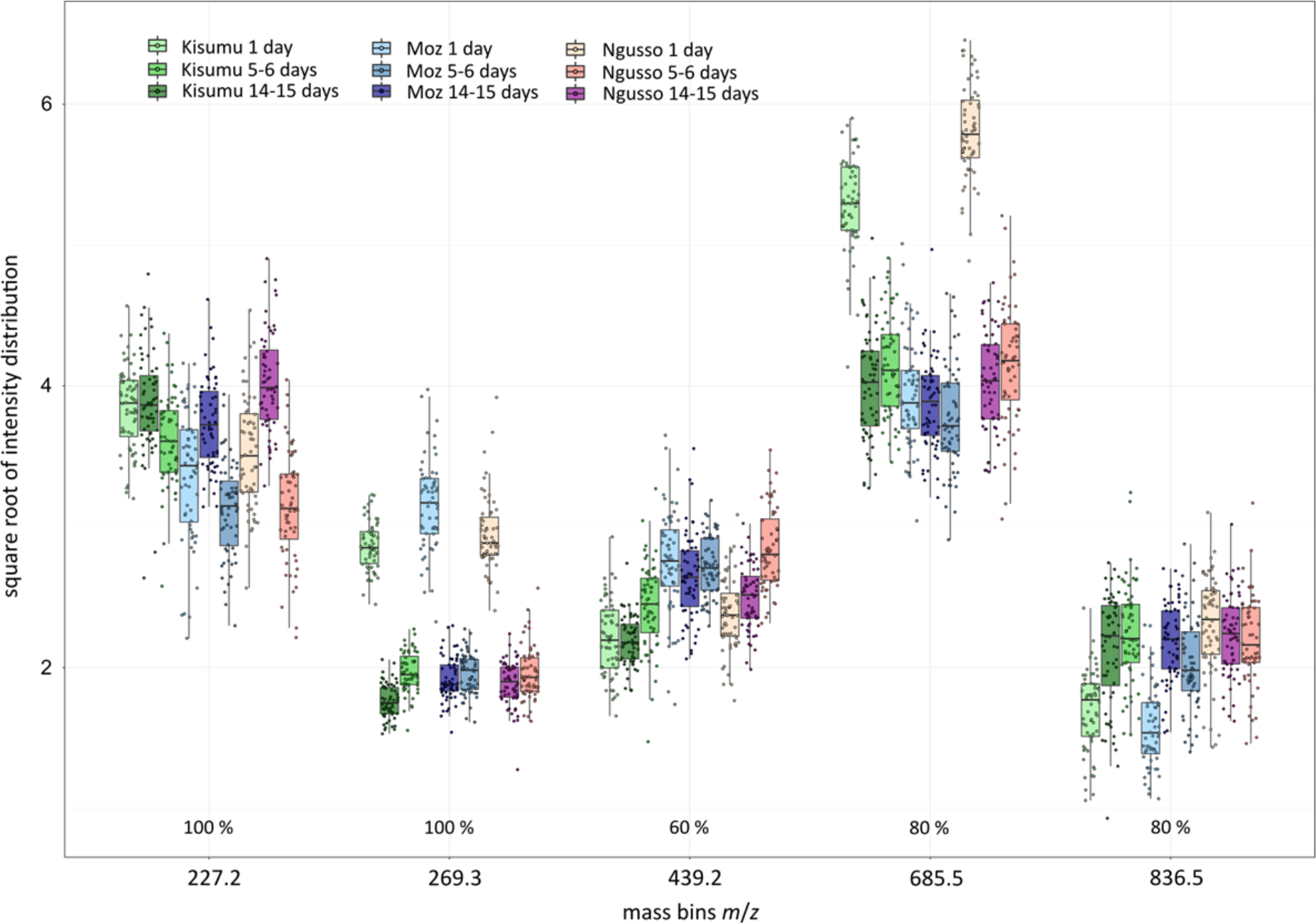
The top 10 most important variables were collated from ten repeated random forest analyses of the two-factor species/age model. Variables which had been identified as separation drivers in more than half the runs were selected to have their intensities plotted. The first two variables, identified 100 % of the time, m/z 227.2 and 269.3 had also been identified in the age model as important separators. The fact that they have also been identified in the nine-class model, in all 10 runs, confirms their importance for age separation. One of the two main separators of the species model, m/z 439.2, also features in this model’s variable list. The other two variables 685.5 and 836.5 have not been identified and seem to be uniquely important for this two-factor model, separating the 1-day old classes of the three species.

**Supplemental Figure 26:**
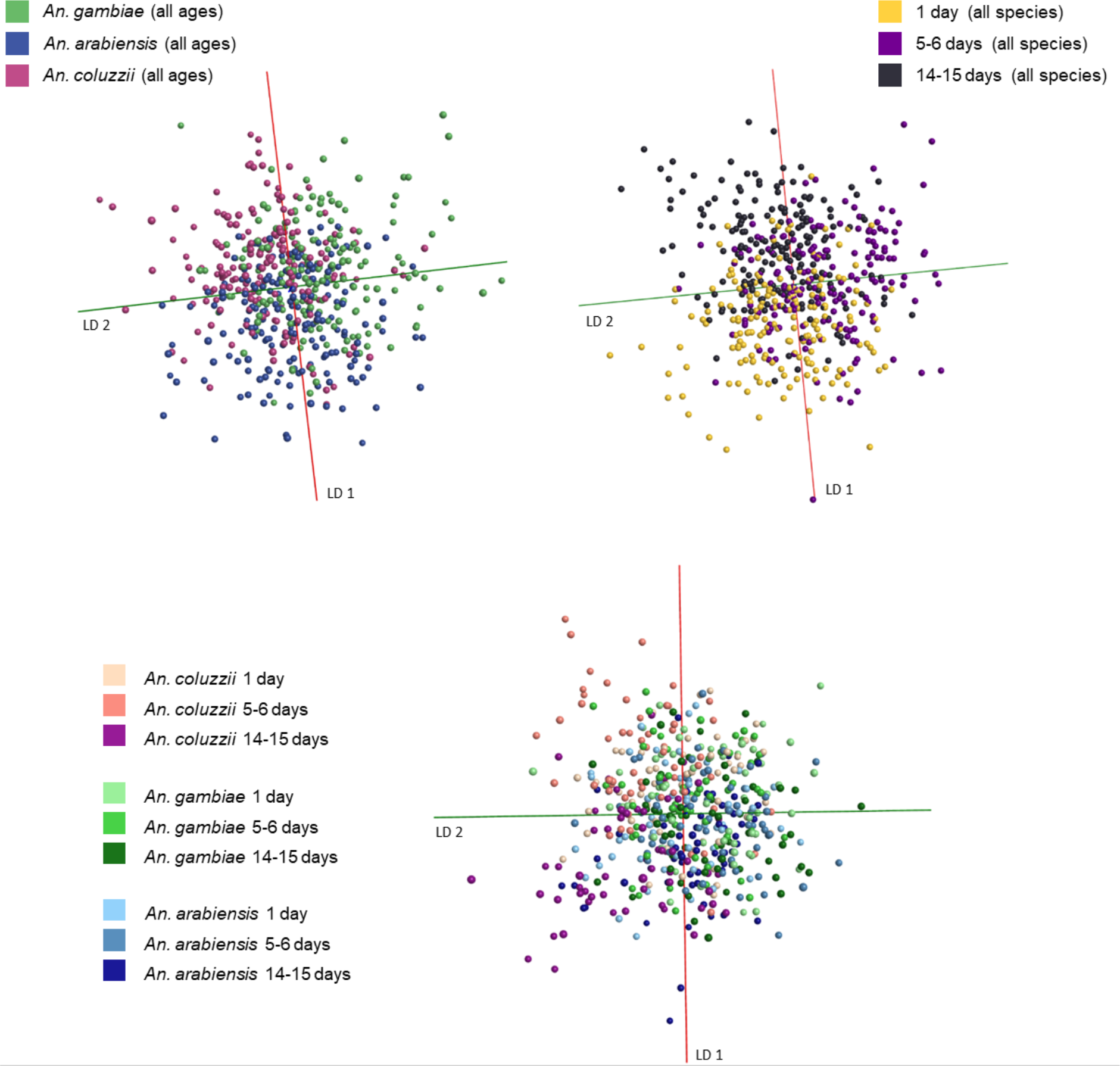
After using 540 Anopheles mosquito specimens, from three species and three age groups each, to build models separating species, age as well as both properties at once, models were re-built with randomly assigned classifications. When re-building the PC-LDA models in Offline Model Builder with classes randomly assigned to samples, separation failed for all three models.

**Supplemental Figure 27:**
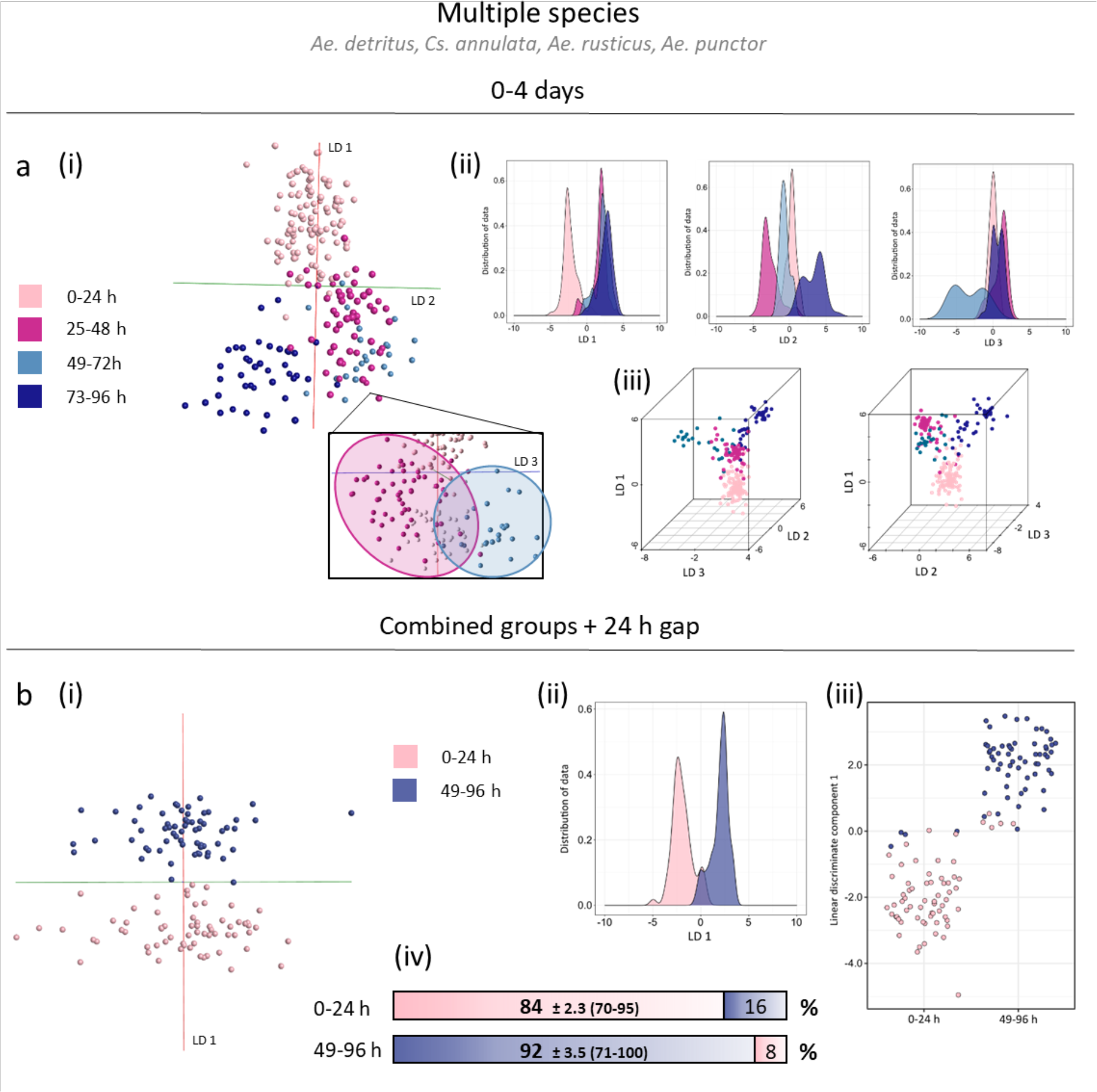
Samples from 4 species (Aedes detritus, Culiseta annulata, Aedes rusticus, Aedes punctor) are included in these age models, separating age groups between 0 and 4 days. Separation is demonstrated using four adjacent age groups (panel a), as well as 2 groups separated by a 24 h gap (panel b). First models were built within OMB using PC-LDA (i), before exporting the matrix and conducting PC-LDA in R, depicted in form of kernel density plots (ii) and scatter plots - 3D and 2D (iii). The age model based on two age groups promised sufficient separation to be used for classification and was therefore additionally analysed via random forest (iv), using 70 % of samples for model building and 30 % for testing (1200 trees). The random forest result is presented in two bars stating the correct classification percentage, including SEM value and the range of achieved accuracies in 10 runs (min and max), and the percentage of misclassified test samples. Samples numbers used for model in panel a: 0-24 h (108), 25-48 h (55), 49-72 h (23), 73-96 h (40). Sample numbers used for model in panel b: 0-24 h (65), 49-96 (63); sample numbers from 0-24 h were reduced for random forest analysis. Separation in panel a were based on 100 (OMB) and 130 PCs (R). Separation in panel b were based on 60 (OMB) and 65 PCs (R).

**Supplemental Figure 28:**
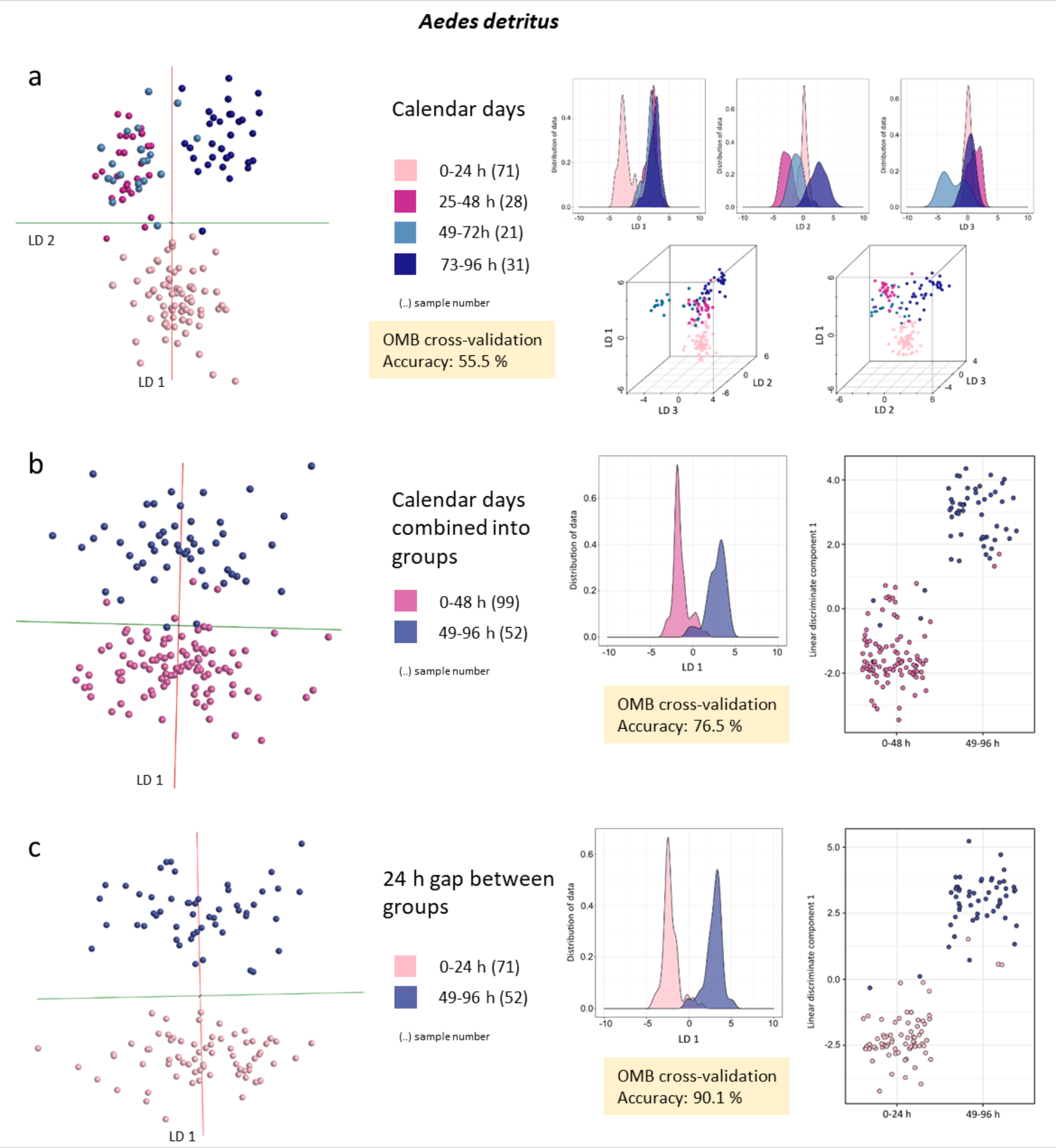
Original and improved age models including only Aedes detritus specimens. The original age model (a) comprises four consecutive age groups demonstrating separation of calendar days. Due to the continuous nature of these classes, separation accuracy is low. Combination of groups (b) reduces the overall class overlap in the model, subsequently improving separation efficiency. Introduction of a 24 h gap between age groups (c) helps to enhance the difference between mosquitoes of different ages even further. All results are based one PC-LD analysis, depicted in form of OMB models and kernel density and scatter plots produced in R (from left to right). The correct classification rates, achieved through ‘Leave 20 % out’ cross-validation in OMB, are highlighted in yellow for each model. The number of samples per class are listed in brackets after the age information.

**Supplemental Figure 29:**
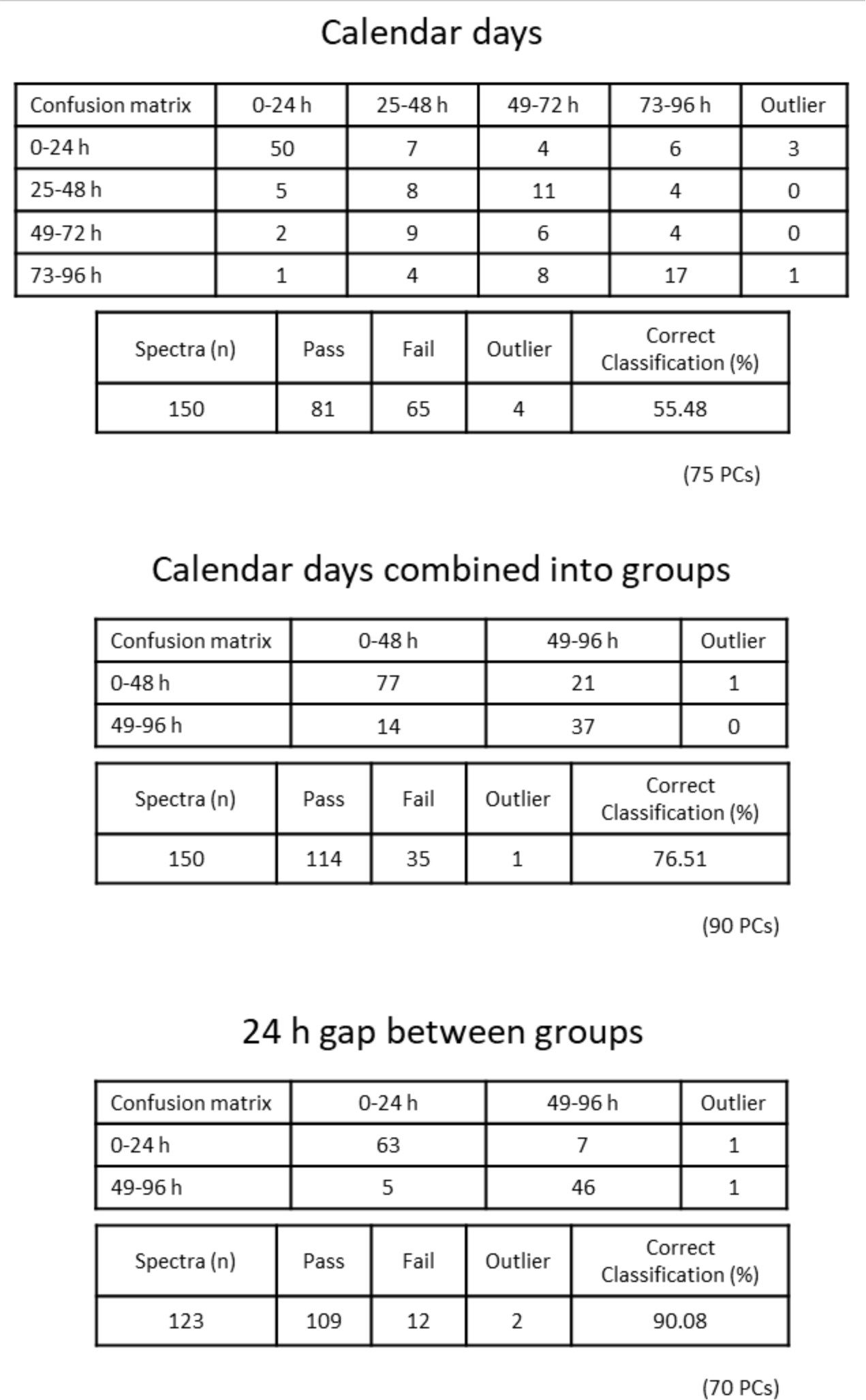
The three Aedes detritus age models were tested via cross-validation in OMB using the option ‘Leave out 20 %’ and a standard deviation of 5. The number of principal components used for model building are given in brackets underneath the tables. One sample each was left out from the first two models as 20 % of 151 samples results in a fractional number that is rounded to the nearest integer.

**Supplemental Figure 30:**
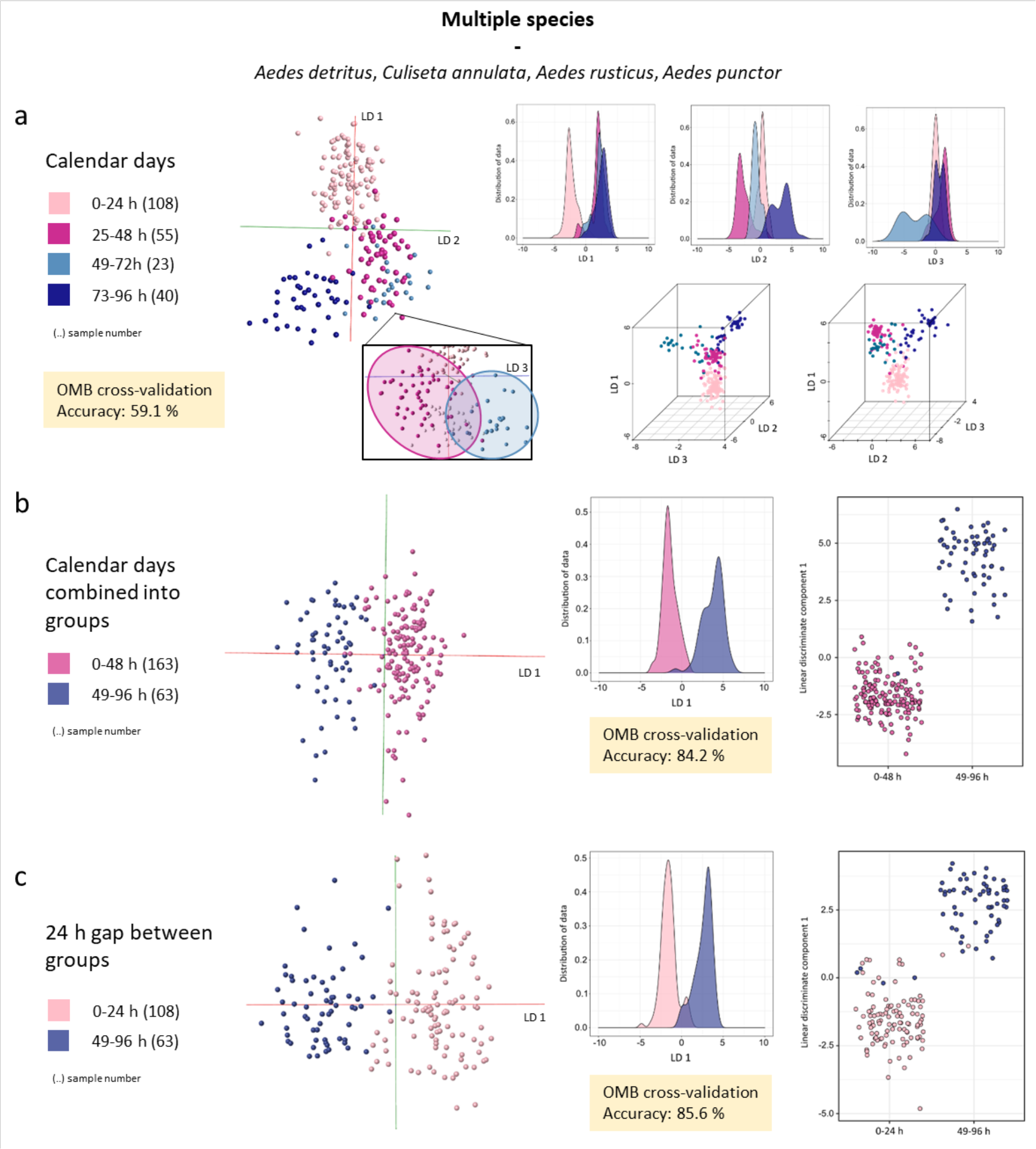
The original and improved age models including specimens from four species: Aedes detritus, Culiseta annulata, Aedes rusticus and Aedes punctor. The original age model (a) comprises four consecutive age groups demonstrating separation of calendar days. Due to the continuous nature of these classes, separation accuracy is low. Combination of groups (b) reduces the overall class overlap in the model, subsequently improving separation efficiency. Introduction of a 24 h gap between age groups (c) helps to enhance the difference between mosquitoes of different ages even further. All results are based one PC-LD analysis, depicted in form of OMB models and kernel density and scatter plots produced in R (from left to right). The correct classification rates, achieved through ‘Leave 20 % out’ cross-validation in OMB, are highlighted in yellow for each model. The number of samples per class are listed in brackets after the age information.

**Supplemental Figure 31:**
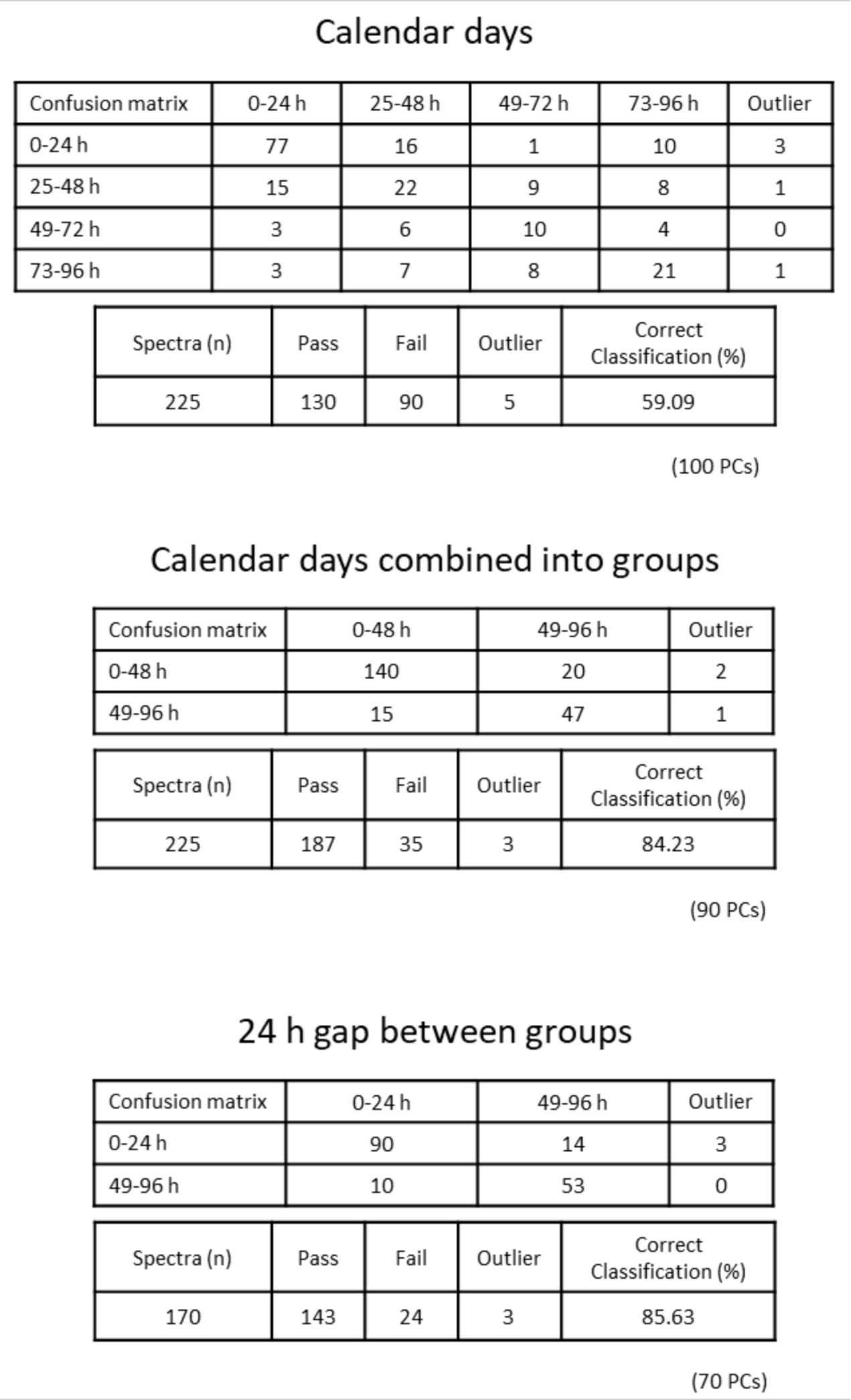
The three multi-species age models were tested via cross-validation in OMB using the option ‘Leave out 20 %’ and a standard deviation of 5. The number of principal components used for model building are given in brackets underneath the tables. One sample each was left out from all models as 20 % of 226 and 171 samples results in fractional numbers that are rounded to the nearest integer.

**Supplemental Figure 32:**
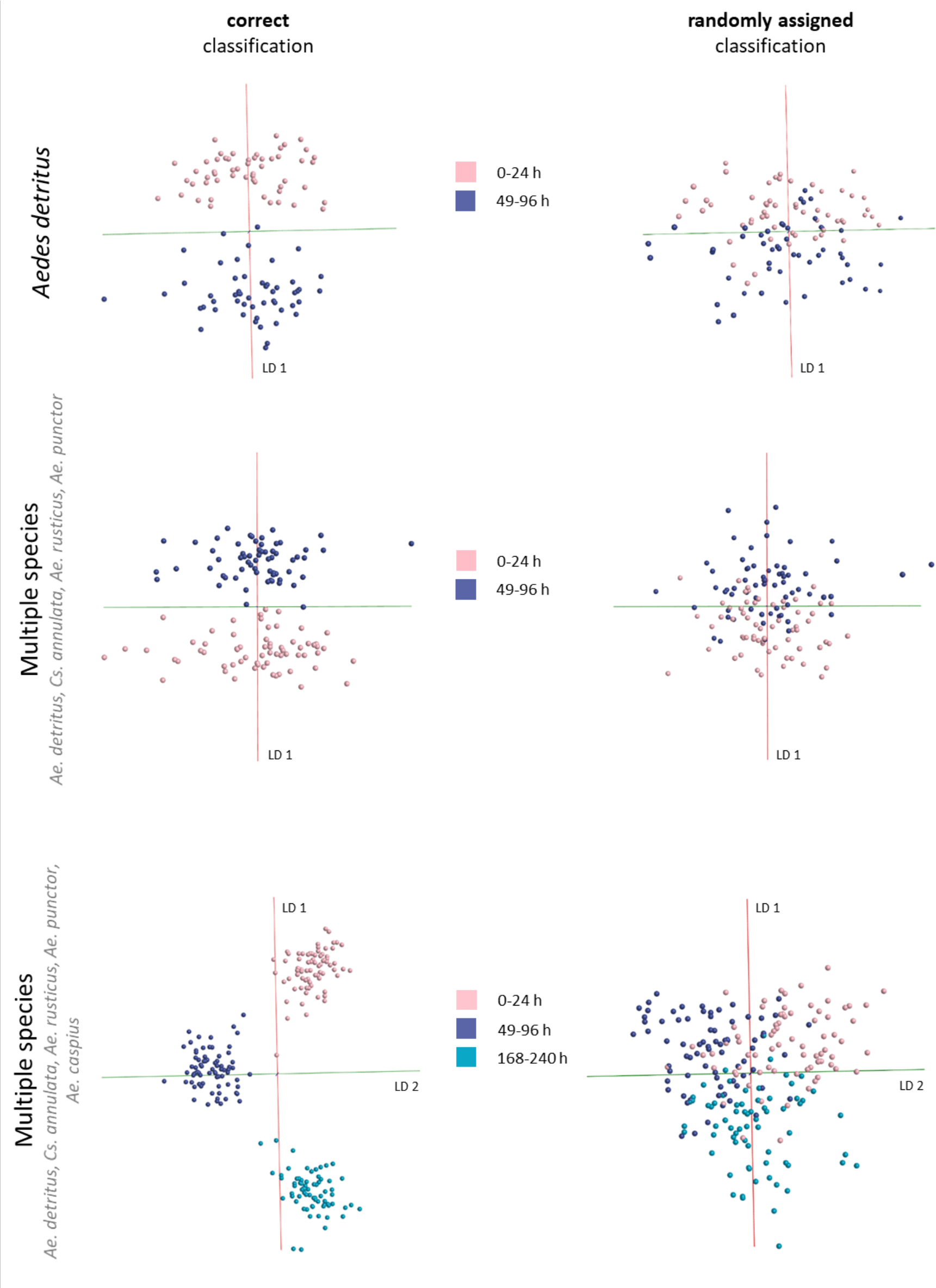
All age models with a 24 h gap between age classes, one Aedes detritus and two multi-species ones, were rebuilt using randomly assigned classifications. The PC-LDA based models built with correct (left) and randomly assigned classifications (right) are listed for comparison. Randomly assigned classifications lead to a considerably worse separation, with individual samples being scattered and classes overlapping. The number of principal components and other settings used for model building were identical for both approaches.

**Supplemental Figure 33:**
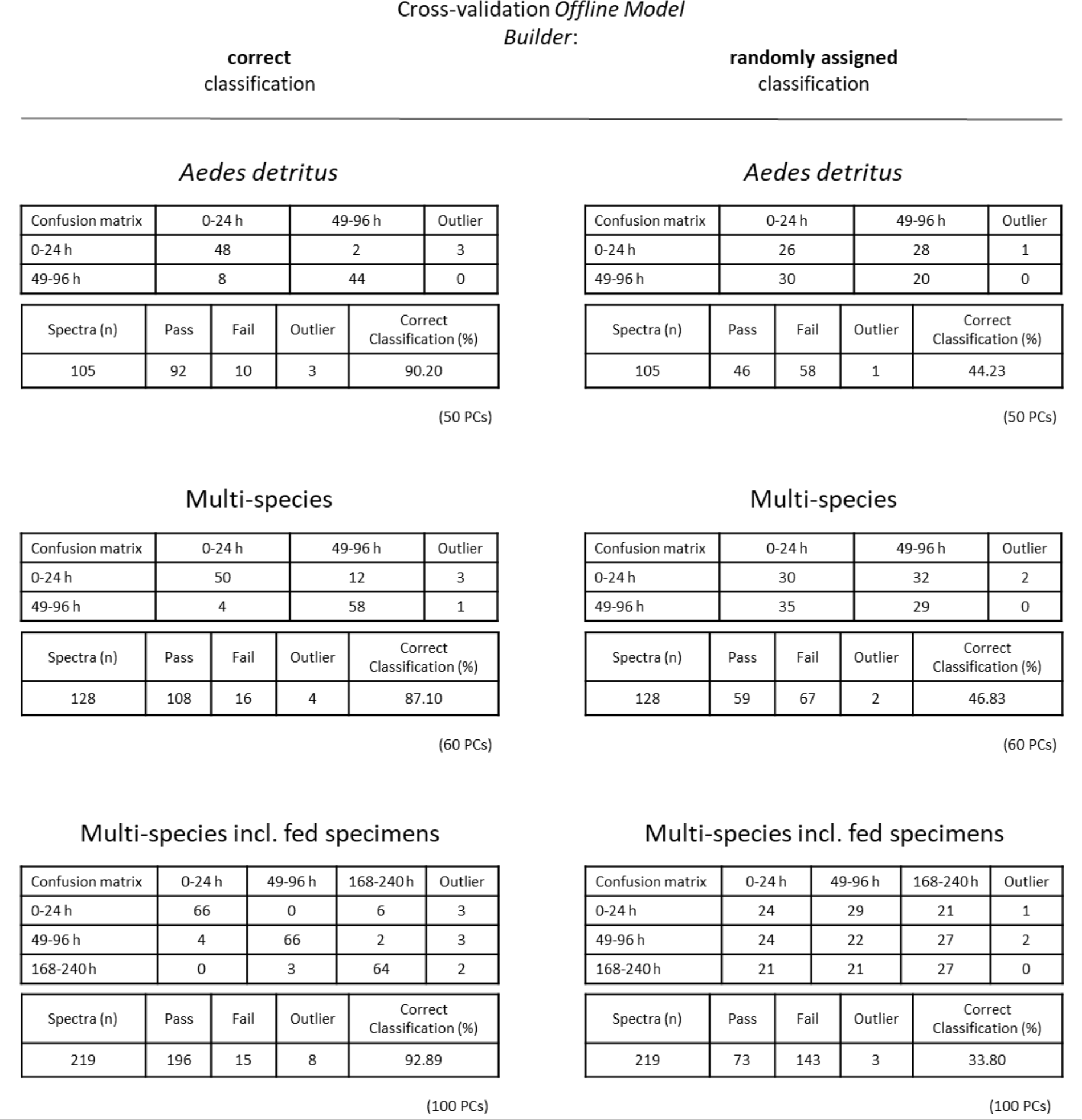
The three main age models, with correct and randomly assigned classifications, were tested via cross-validation in OMB using the option ‘Leave out 20 %’ and a standard deviation of 5. The number of principal components used for model building are given in brackets underneath the tables. Two samples from the Aedes detritus age model were left out as 20 % of 107 samples results in a fractional number that is rounded to the nearest integer.

**Supplemental Figure 34:**
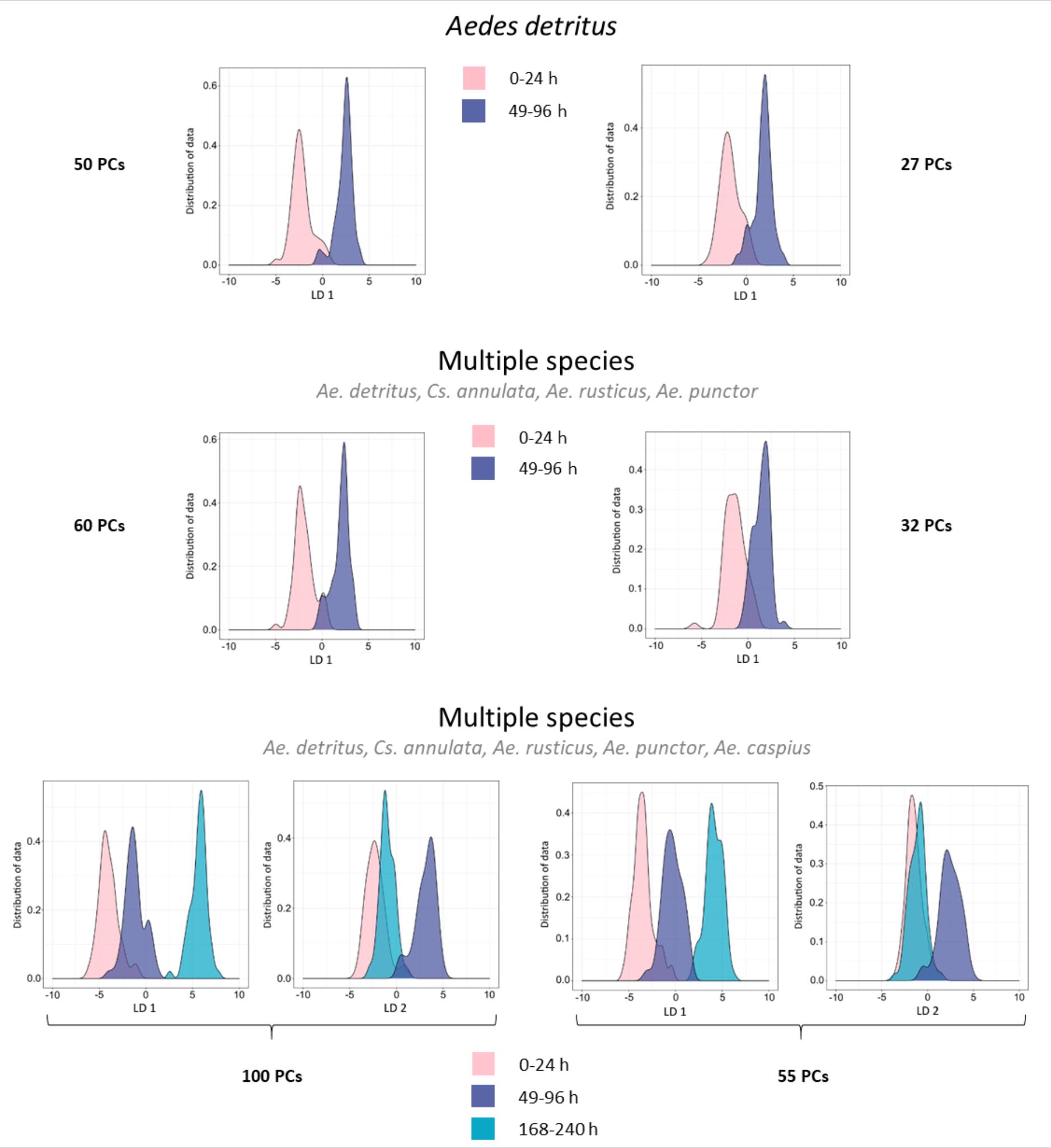
The PC-LDA separation achieved for the three main age models is presented here using the maximum number of principal components possible before overfitting (left side), as well as using only a quarter of possible PCs (right side).

**Supplemental Figure 35:**
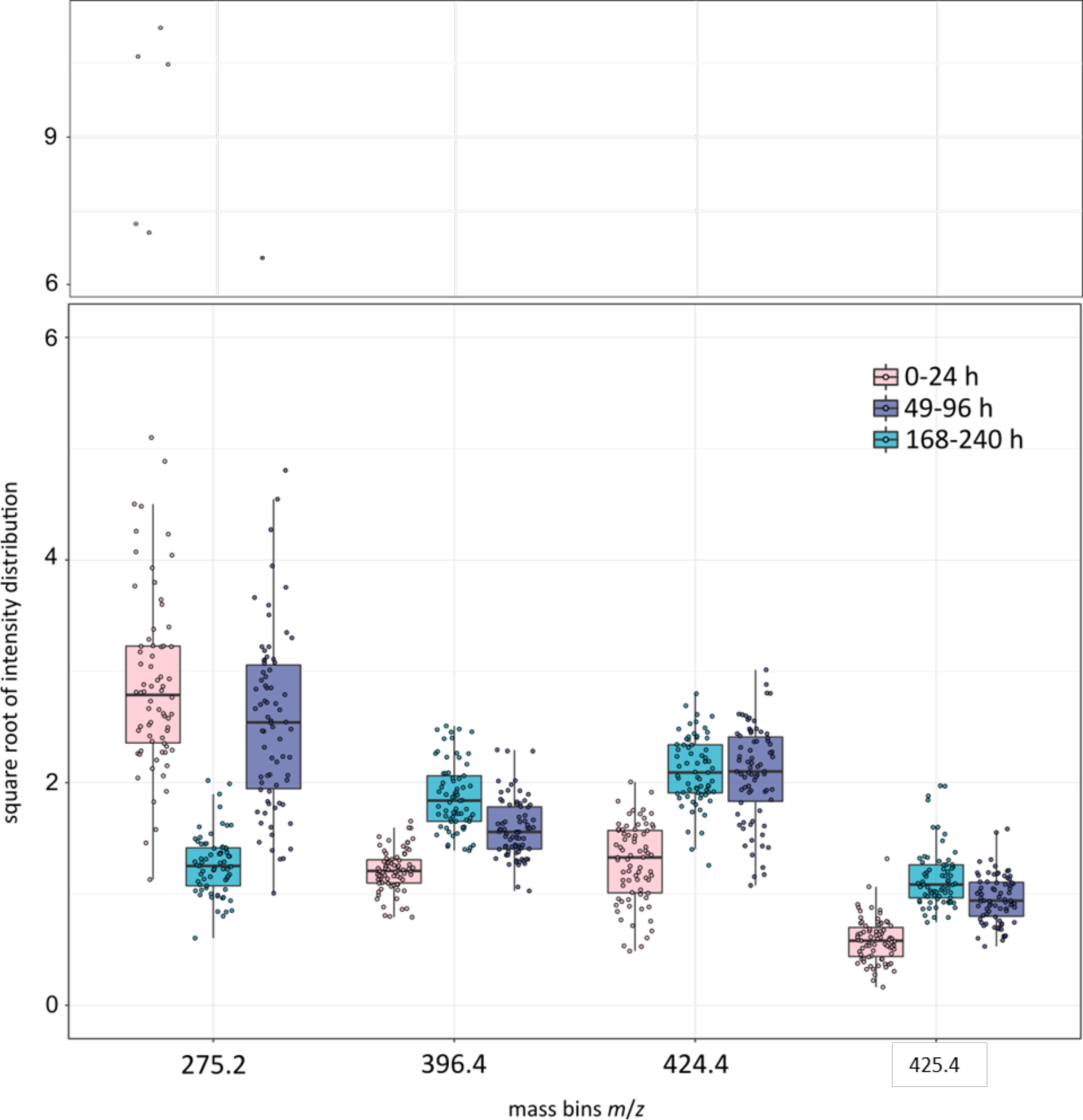
After performing random forest analysis (repeated 10 times) on the age model (Figure 5.20) the R package ‘randomForestExplainer’ was used to determine the ion bins driving the separation process using a Top 10 approach. Four variables were identified as important in all 10 random forest runs. The intensities of all 219 samples were plotted for these bins in a boxplot diagram. A second panel with compacted y-axis is placed on top to show separated values for bin m/z 275.2

**Supplemental Figure 36:**
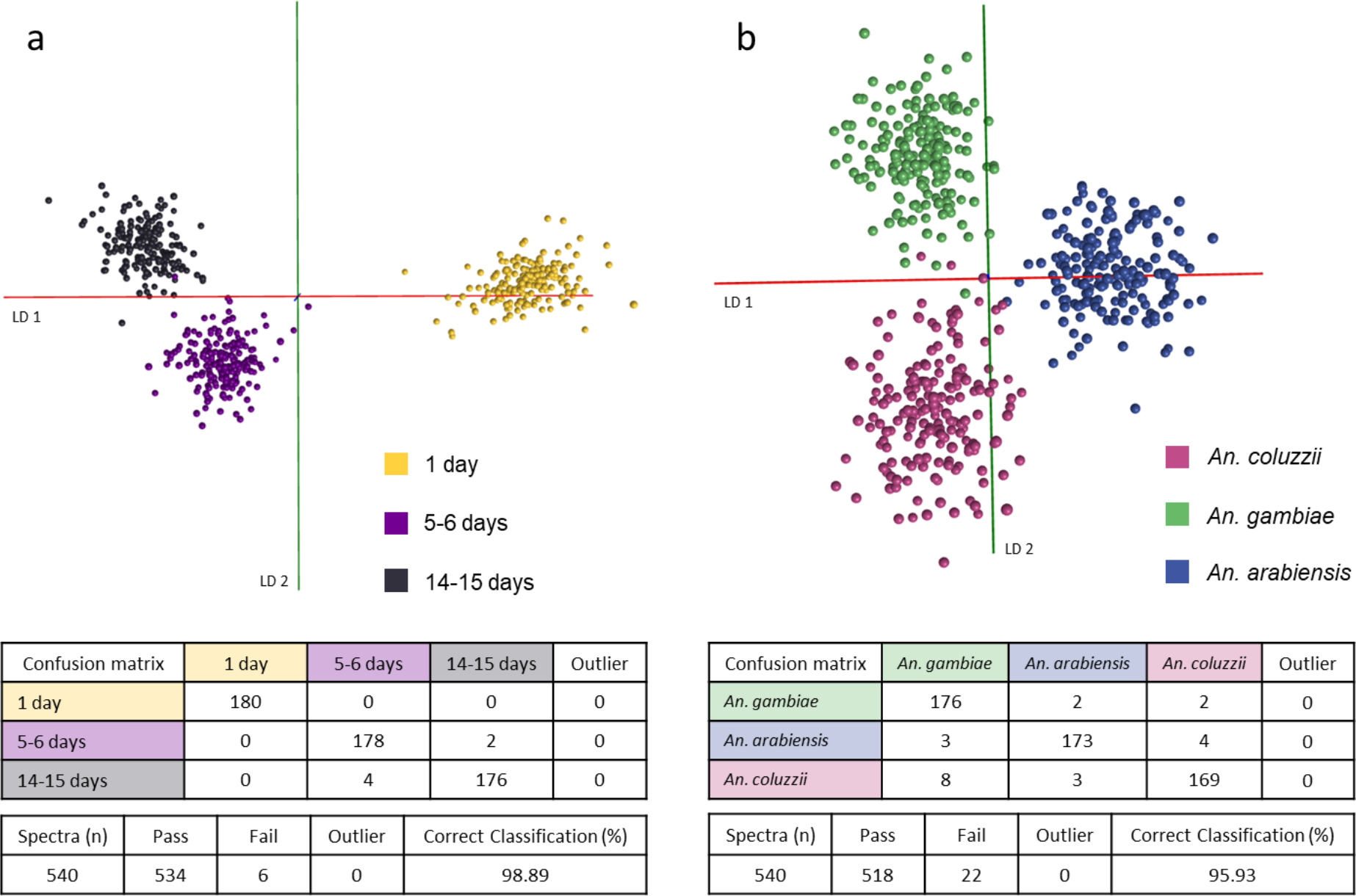
Models separating age groups (1 day, 5-6 days, 14-15 days; 180 samples each) and species classes (An. coluzzii, An. gambiae, An. arabiensis; 180 samples each) were re-built in Offline Model Builder using a bin size of 1 m/z. Models were cross-validated in OMB (‘Leave 20 % out’, standard deviation 5).

**Supplemental Figure 37:**
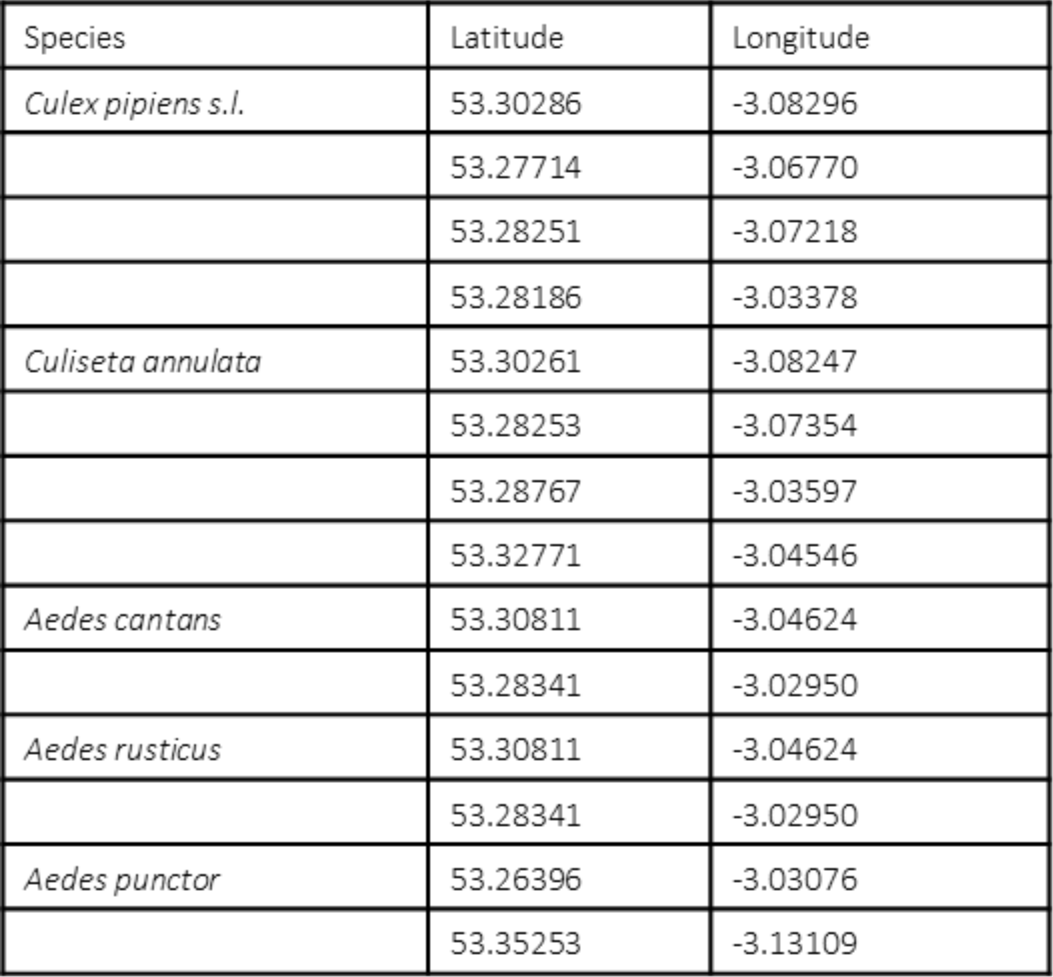
Coordinates of the locations where immature mosquito specimens were collected.

