## Supplementary Material for "Rapid identification of mosquito species, sex and age by mass spectrometric analysis"

** Joint corresponding co-authors

**Supplemental Figure 1:**

*The data matrix, obtained after processing and binning the mass spectral data in Offline Model Builder, was used to create averaged mass spectra for all three species. Each mass spectrum represents an average of all samples available for each species (*Anopheles coluzzii *n=54,* Anopheles gambiae *n=59,* Anopheles arabiensis *n=89).*

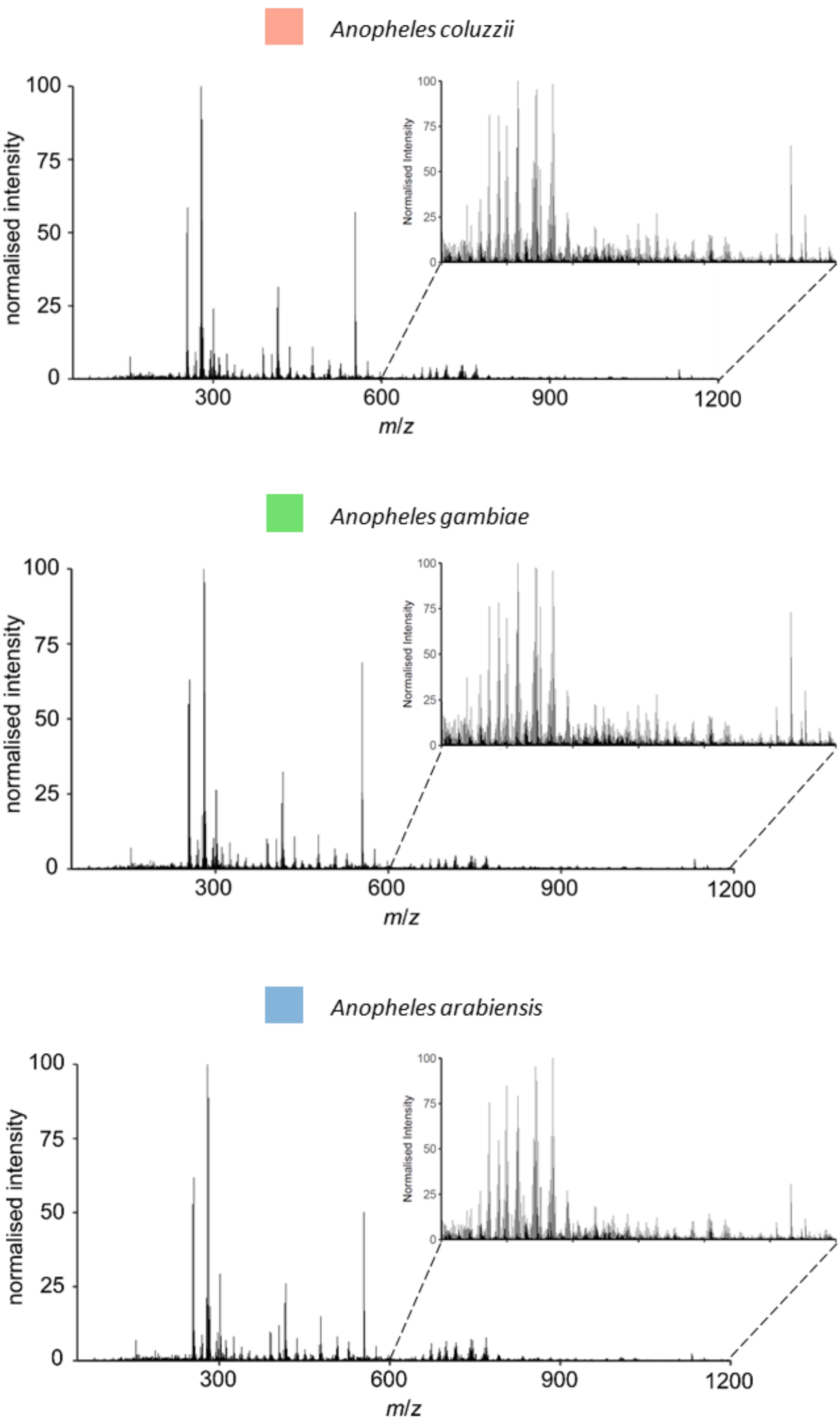

**Supplemental Figure 2:**

*The PC-LDA model separating* An. coluzzii, An. gambiae *and* An. arabiensis*, built in Offline Model Builder using 90 PCs (left), was re-built after randomly assigning classifications to samples (right). The random classification model, also based on 90 PCs, displays no separation of the three species; samples are widely dispersed and groups strongly overlap.*

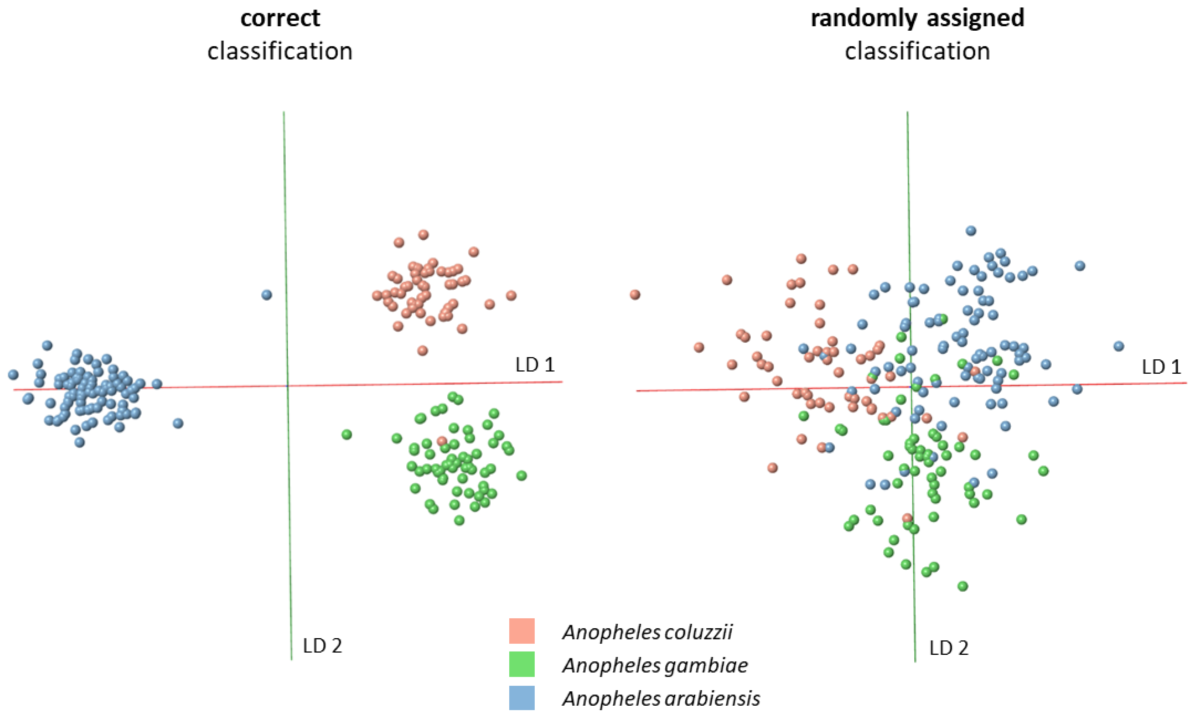

**Supplemental Figure 3:**
*The model separating* An. coluzzii*,* An. gambiae *and* An. arabiensis *was re-built using a lower number of principal components. PC number was decreased to 50, which is ¼ of the maximum number possible. As can be seen in the OMB model (a) as well as the kernel density- and scatter plots (b), reduced variance in the model still resulted in a clear separation of all three species.*

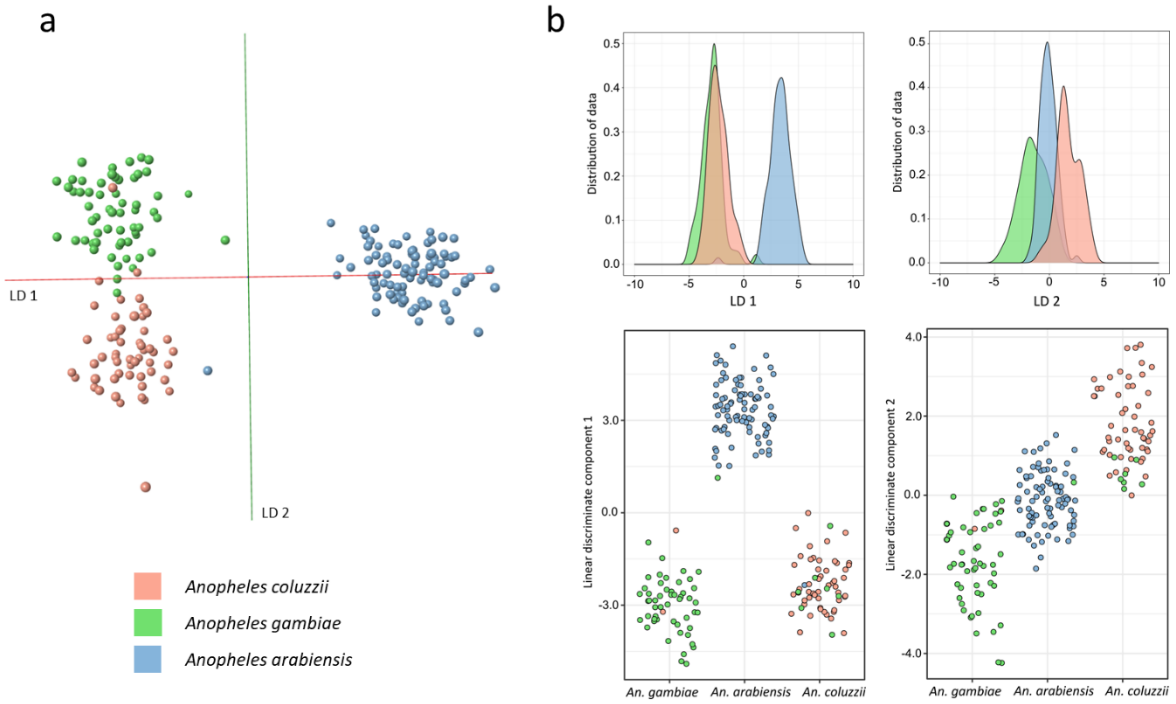

**Supplemental Figure 4:**

*The PCA-LDA based species models (with correct classification, built with 90 and 50 PCs; with randomly assigned classes, built with 90 PCs) were cross-validated within Offline Model Builder using the setting 'Leave 20 % out' and a standard deviation of 5. During cross-validations two samples of the species* An. gambiae *and/or* An. arabiensis *were not tested as 20 % of 202 results in a fractional number.*

Cross-validation *Offline Model Builder*:

**correct** classification

90 PCs

| Confusion matrix | *An. gambiae* | *An. arabiensis* | An. coluzzii | Outlier |
|---|---|---|---|---|
| *An. gambiae* | 53 | 0 | 3 | 1 |
| *An. arabiensis* | 0 | 87 | 1 | 1 |
| *An. coluzzii* | 5 | 0 | 48 | 1 |

| Spectra (n) | Pass | Fail | Outlier | Correct Classification (%) |
|---|---|---|---|---|
| 200 | 188 | 9 | 3 | 95.43 |

50 PCs

| Confusion matrix | *An. gambiae* | *An. arabiensis* | An. coluzzii | Outlier |
|---|---|---|---|---|
| *An. gambiae* | 50 | 1 | 7 | 0 |
| *An. arabiensis* | 0 | 87 | 1 | 0 |
| *An. coluzzii* | 6 | 0 | 48 | 0 |

| Spectra (n) | Pass | Fail | Outlier | Correct Classification (%) |
|---|---|---|---|---|
| 200 | 185 | 15 | 0 | 92.50 |

**randomly assigned** classification

90 PCs

| Confusion matrix | *An. gambiae* | *An. arabiensis* | An. coluzzii | Outlier |
|---|---|---|---|---|
| *An. gambiae* | 16 | 29 | 13 | 0 |
| *An. arabiensis* | 24 | 40 | 23 | 1 |
| *An. coluzzii* | 22 | 17 | 15 | 0 |

| Spectra (n) | Pass | Fail | Outlier | Correct Classification (%) |
|---|---|---|---|---|
| 200 | 71 | 128 | 1 | 35.68 |

**Supplemental Figure 5:**

*Female specimens from the three species* An. arabiensis, An. gambiae *and* An. coluzzii *were equally split into two groups: one group was killed through dehydration and stored at room temperature with desiccant material, the other group was killed by freezing and stored at -20˚C in falcon tubes. Within each group samples were additionally split to be analysed at five different time points: immediately after killing (no storage) and after storage for 1, 2, 4 and 10 weeks. For every combination of storage type and length 5* An. coluzzii*, 5* An. gambiae *and 3* An. arabiensis *mosquitoes were analysed. For both storage conditions (desiccated and frozen) samples from all storage time points were combined to build PC-LDA based species models in OMB (based on 35 and 30 PCs). Both storage types, dry at room temperature (panel a) and frozen at -20˚C (panel b) led to informative REIMS spectra allowing differentiation of species through PC-LD analysis. Following successful separation in individual models, all samples were combined (n=130) for species classification (c). First, PC-LDA was attempted in Offline Model Builder (left) before exporting the data matrix and conducting the analysis in R (right); both were based on 70 PCs. Despite the large amount of variability in the sample set specimens were clustered into their respective species group.*

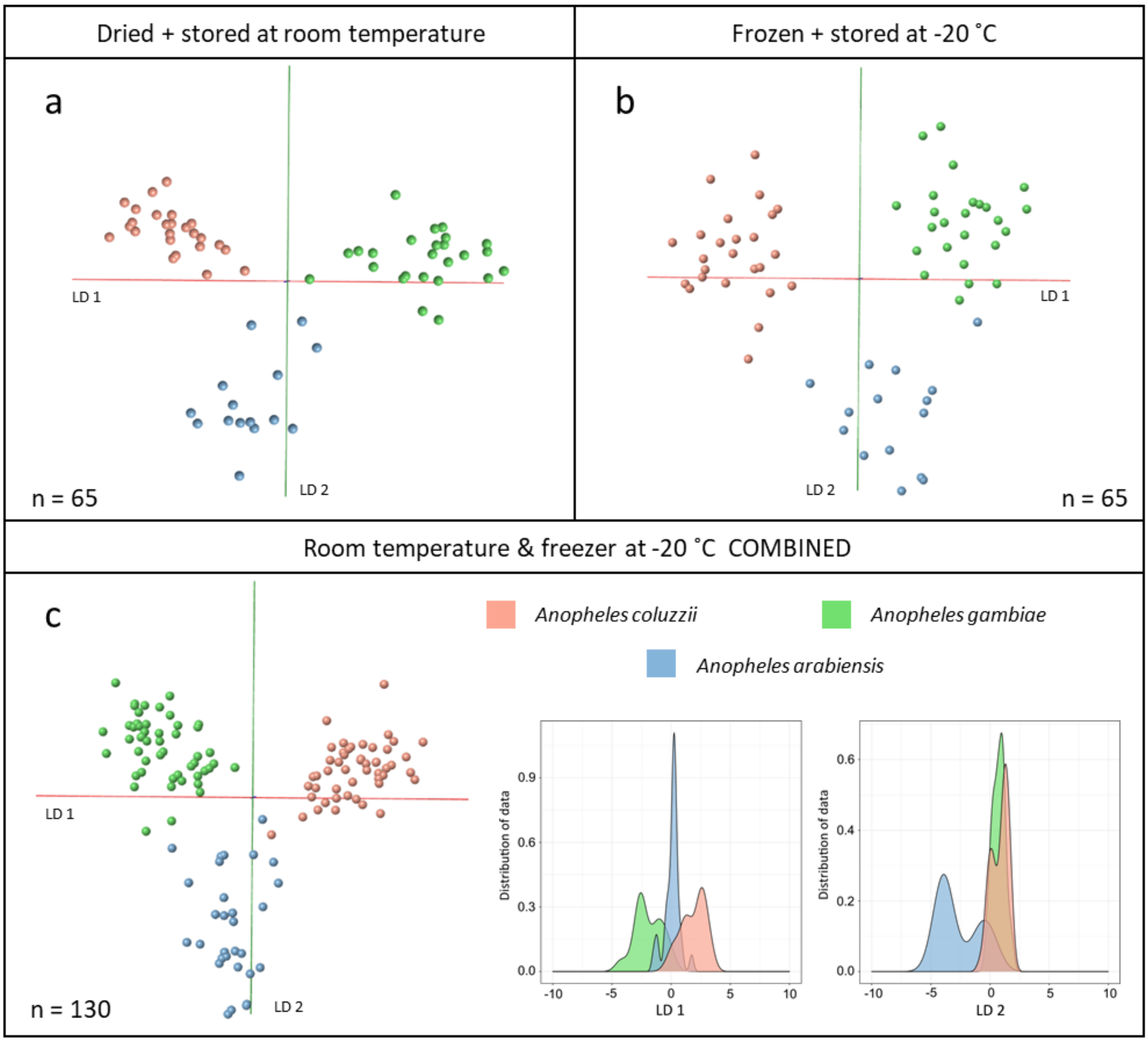

**Supplemental Figure 6:**

*Laboratory raised* Anopheles gambiae *mosquitoes were separated into male (n=54) and female (n=61) classes, based on the sexual dimorphism of their antennae (d), before being analysed through REIMS. Data were analysed in the Offline Model Builder software (PC-LDA) as well as the R environment (PC-LDA, random forest). The results of PC-LDA (conducted in R, based on 60 PCs) are visualised in form of kernel density (a) and scatter plots (b), which both depict good separation of the sexes with only a small amount of samples (5) overlapping. Additionally, random forest analysis was conducted, using a 70 %/30 % split for model training and testing; the analysis was repeated 10 times with different samples in the training and testing category each time (randomly selected). The averaged results are listed as percentages in the confusion matrix (c) with SEM ± and the range of achieved accuracies (min-max) stated for the correct classification percentages. The average number of samples used for testing are listed at the end of the class rows (n=x). PC-LDA models, based on 60 PCs and 29 PCs, were cross-validated within OMB using the setting 'Leave 20 % out' and a standard deviation of 5 (e, upper panel). Furthermore, sample classifications (male, female) were randomly assigned to samples and the model was re-built using the initial principal component number (60 PCs) (e, bottom panel). The resulting separation is noticeably worse with nearly half of the male samples completely overlapping with the female class and a correct classification rate of only 52 %.*

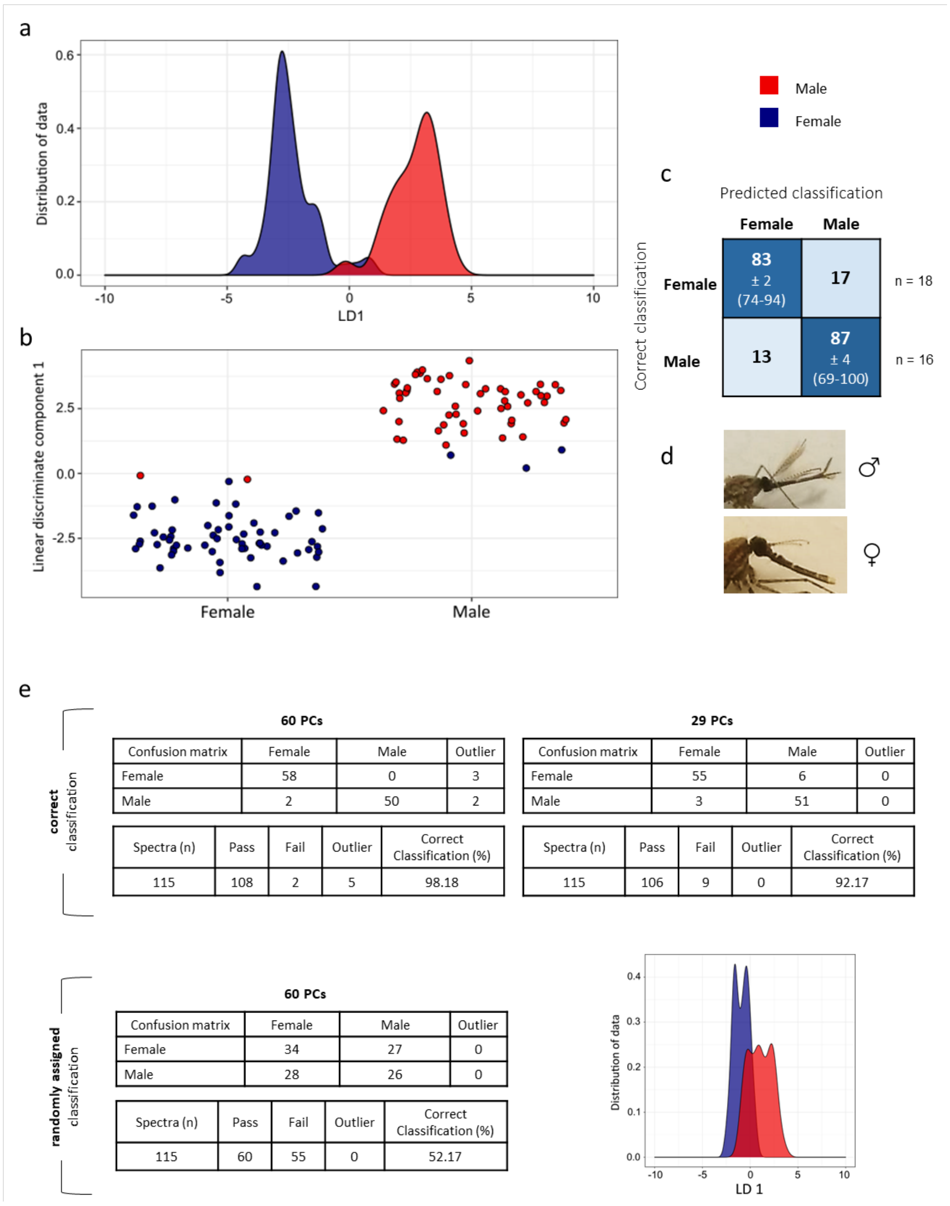


**correct classification — 60 PCs**

| Confusion matrix | Female | Male | Outlier |
|---|---|---|---|
| Female | 58 | 0 | 3 |
| Male | 2 | 50 | 2 |

| Spectra (n) | Pass | Fail | Outlier | Correct Classification (%) |
|---|---|---|---|---|
| 115 | 108 | 2 | 5 | 98.18 |

**correct classification — 29 PCs**

| Confusion matrix | Female | Male | Outlier |
|---|---|---|---|
| Female | 55 | 6 | 0 |
| Male | 3 | 51 | 0 |

| Spectra (n) | Pass | Fail | Outlier | Correct Classification (%) |
|---|---|---|---|---|
| 115 | 106 | 9 | 0 | 92.17 |

**randomly assigned classification — 60 PCs**

| Confusion matrix | Female | Male | Outlier |
|---|---|---|---|
| Female | 34 | 27 | 0 |
| Male | 28 | 26 | 0 |

| Spectra (n) | Pass | Fail | Outlier | Correct Classification (%) |
|---|---|---|---|---|
| 115 | 60 | 55 | 0 | 52.17 |

**Supplemental Figure 7:**

*Cross-validation results for the seven species model built using 100 PCs (panel a). Cross-validation was performed within OMB using the option 'Leave 20 % out' and a standard deviation of 5. Results are listed in form of a confusion matrix containing the numbers of samples which have been either correctly or wrongly classified, as well as the number of outliers per classifications. The summary underneath contains the total number of spectra (samples) used for validation, the number of passed and failed samples, total number of outliers and the calculated correct classification rate (%) of the model. Random forest analysis of the seven species data set was repeated 10 times, using a different set of samples for model training (70 % of data) and testing (30 % of data) each time. The resulting confusion matrices, containing the numbers of correctly and wrongly classified samples, were turned into percentages and averaged over the 10 runs. The averaged correct classification accuracies (in %) plus SEM (±) and the range of achieved accuracies over 10 repeats (min and max) are listed in the coloured cells (panel b). The column on the right (n) states the average number of samples used for testing for each class. In total, the model achieved a classification accuracy of 91 %; meaning 91 out of 100 test samples would be identified correctly.*

a

**Offline Model Builder**

Predicted classification

Correct classification

| Confusion matrix | *Ae. cantans* | *Ae. caspius* | *Ae. punctor* | *Ae. rusticus* | *Cx. pipiens* | *Cs. annulata* | *Ae. detritus* | Outlier |
|---|---|---|---|---|---|---|---|---|
| *Ae. cantans* | 78 | 0 | 0 | 0 | 1 | 0 | 0 | 1 |
| *Ae. caspius* | 0 | 80 | 0 | 0 | 0 | 0 | 0 | 0 |
| *Ae. punctor* | 0 | 0 | 72 | 1 | 0 | 0 | 4 | 3 |
| *Ae. rusticus* | 0 | 0 | 0 | 79 | 0 | 0 | 1 | 0 |
| *Cx. pipiens* | 1 | 0 | 0 | 0 | 79 | 0 | 0 | 0 |
| *Cs. annulata* | 0 | 0 | 0 | 1 | 0 | 77 | 0 | 2 |
| *Ae. detritus* | 0 | 1 | 1 | 0 | 0 | 0 | 77 | 1 |

| Spectra (n) | Pass | Fail | Outlier | Correct Classification (%) |
|---|---|---|---|---|
| 560 | 542 | 11 | 7 | 98 |

b

**Random Forest**

Predicted classification

Correct classification

| Confusion matrix | *Cx. pipiens* | *Cs. annulata* | *Ae. caspius* | *Ae. cantans* | *Ae. punctor* | *Ae. rusticus* | *Ae. detritus* | n |
|---|---|---|---|---|---|---|---|---|
| *Cx. pipiens* | **98** ± 0.6 (95-100) | 1 | 0 | 1 | 0 | 0 | 0 | 26 |
| *Cs. annulata* | 6 | **94** ± 2.3 (78-100) | 0 | 0 | 0 | 0 | 0 | 24 |
| *Ae. caspius* | 2 | 1 | **93** ± 1.8 (81-100) | 0 | 0 | 0 | 3 | 24 |
| *Ae. cantans* | 0 | 0 | 0 | **87** ± 1.9 (75-96) | 0 | 0 | 12 | 24 |
| *Ae. punctor* | 3 | 0 | 0 | 3 | **85** ± 3.9 (57-100) | 0 | 9 | 23 |
| *Ae. rusticus* | 0 | 0 | 0 | 9 | 0 | **87** ± 2.3 (74-100) | 4 | 25 |
| *Ae. detritus* | 0 | 0 | 0 | 3 | 2 | 1 | **93** ± 1.6 (83-100) | 22 |

Model accuracy: 91 % ± 1.0

**Supplemental Figure 8:**

*Comparison of the PC-LDA based 7-species model built with correct sample classifications (left) and randomly assigned classifications (right). Both models were built with the same settings in Offline Model Builder, using 100 principal components.*

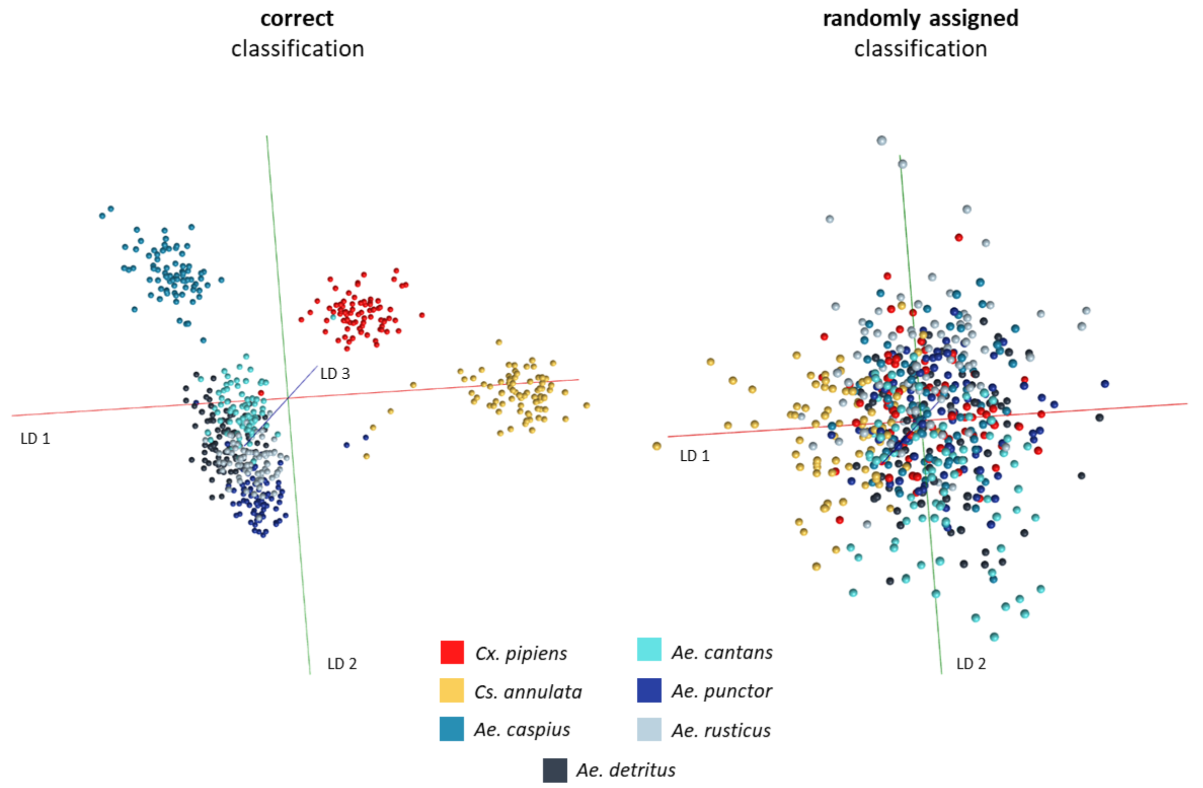

**Supplemental Figure 9:**

*Separation of male (66) and female (70)* Aedes detritus *specimens based on principal component-linear discriminant analysis, displayed in form of a smoothed histogram (a, left) and a scatterplot (a, right) using 80 principal components. The separation was further examined through cross-validation (c) in OMB ('Leave out 20%', standard deviation of 5) as well as random forest analysis (b) in R (70 % training/30 % testing, repeated 10 times). The cross-validated OMB model was based on 70 PCs, one sample was left out during validation as 20 % of 136 samples results in a fractional number that was rounded to the nearest integer. The results of the random forest analysis are depicted in a confusion matrix containing information about the percentages of samples, which had been either correctly or wrongly classified, including the standard error of the mean (±) and the range of accuracies achieved (min and max) for the correct classifications. The average number of samples (n) tested from each class is listed on the left-hand side of the table and the average model accuracy (plus SEM) underneath. Additionally, the PC-LDA based model was re-built with randomly assigned classifications (d, left) and again tested via cross-validation in OMB (d, right), which resulted in the expected decrease of classification accuracy and confirmed that separation (with correct classifications) is based on sex related variance.*

*Aedes detritus*

a

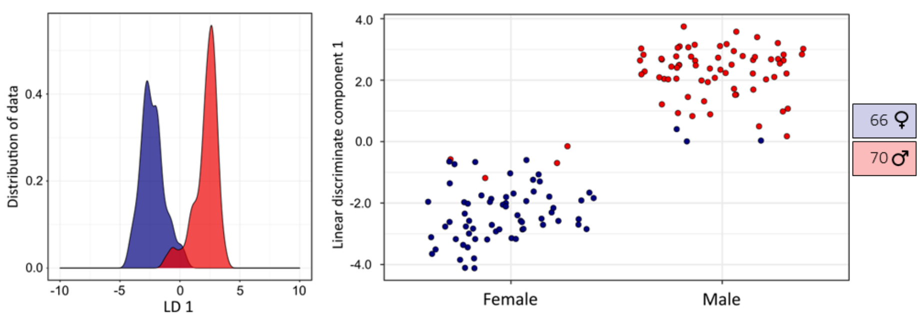


b

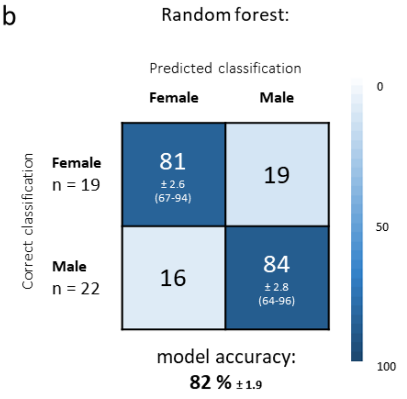


c Cross-validation *Offline Model Builder*:

| Confusion matrix | Female | Male | Outlier |
|---|---|---|---|
| Female | 52 | 14 | 0 |
| Male | 8 | 59 | 2 |

| Spectra (n) | Pass | Fail | Outlier | Correct Classification (%) |
|---|---|---|---|---|
| 135 | 111 | 22 | 2 | 83.46 |

d

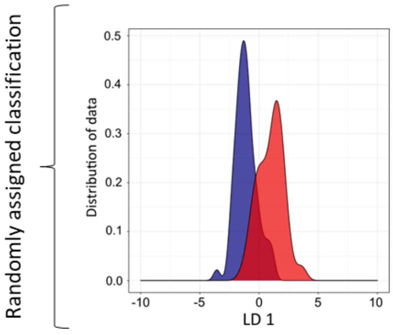


Cross-validation *Offline Model Builder*:

| Confusion matrix | Female | Male | Outlier |
|---|---|---|---|
| Female | 29 | 33 | 1 |
| Male | 38 | 34 | 0 |

| Spectra (n) | Pass | Fail | Outlier | Correct Classification (%) |
|---|---|---|---|---|
| 135 | 63 | 71 | 1 | 47.01 |

**Supplemental Figure 10:**

*Male (60) and female (60) specimens were equally selected from four different mosquito species (*Aedes detritus, Aedes punctor, Aedes rusticus, Aedes cantans*) to test for species independent separation of sexes. The separation is based on principal component-linear discriminant analysis (70 principal components) and displayed in form of a smoothed histogram (a, left) and a scatterplot (a, right). The separation was further examined through cross-validation (c) in OMB ('Leave out 20%', standard deviation of 5) as well as random forest analysis (b) in R (70 % training/30 % testing, repeated 10 times). The cross-validated OMB model was also based on 70 PCs. The results of the random forest analysis are depicted in a confusion matrix containing information about the percentages of samples, which had been either correctly or wrongly classified, including the standard error of the mean (±) and the range of accuracies achieved (min and max) for the correct classifications. The average number of samples (n) tested from each class is listed on the left-hand side of the table and the average model accuracy (plus SEM) underneath. Additionally, the PC-LDA based model was re-built with randomly assigned classifications (d, left) and again tested via cross-validation in OMB (d, right), which confirmed that separation (with correct classifications) is based on sex related variance, despite including multiple species.*

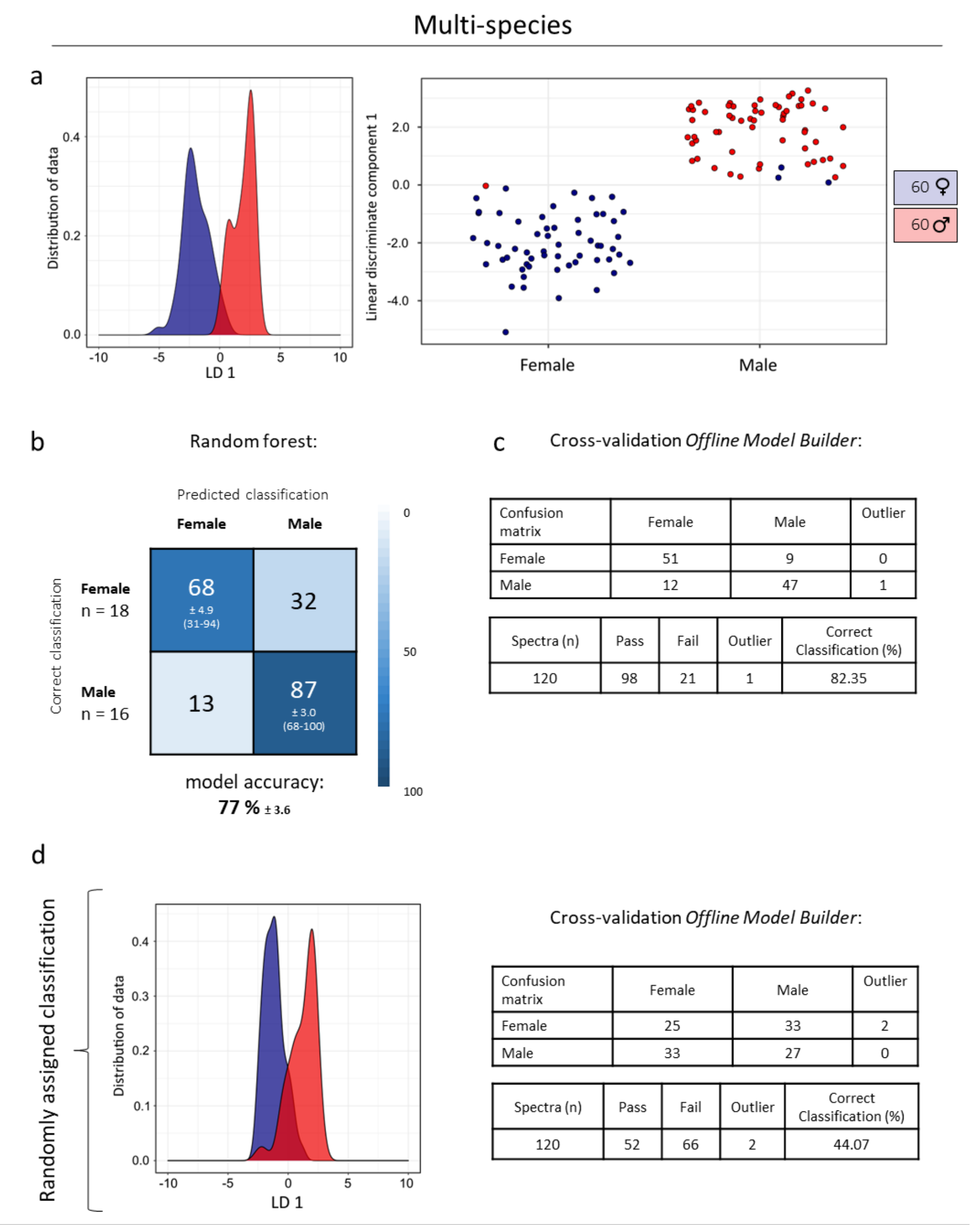

**Supplemental Figure 11:**

*Identifications results of samples (raised) analysed in the same year as samples used for model building, listed for each species. The percentage of correctly identified samples and the probability that the identification is correct are highlighted in yellow for easier comparison.*

| UNKNOWNS 2019 I | *Aedes cantans* (n = 3) | | *Culiseta annulata* (n = 22) | | *Aedes detritus* (n = 16) | | *Aedes punctor* (n = 11) | | *Aedes rusticus* (n = 4) | |
|---|---|---|---|---|---|---|---|---|---|---|
| | correct | wrong | correct | wrong | correct | wrong | correct | wrong | correct | wrong |
| Percentage of all samples [%] | 100 | 0 | 100 | 0 | 93.8 | 6.3 | 90.9 | 9.1 | 75 | 25 |
| Probability of correctness > 80 % | 100 | 0 | 100 | 0 | 100 | 100 | 100 | 100 | 100 | 0 |
| Probability of correctness < 80 % | 0 | 0 | 0 | 0 | 0 | 0 | 0 | 0 | 0 | 100 |
| Average probability of correctness | 96.6 | 0 | 96.7 | 0 | 98.8 | 99.2 | 99.1 | 80.4 | 98.8 | 77.4 |
| Number of outliers (StDev > 10) | 0 | 0 | 0 | 0 | 0 | 0 | 0 | 0 | 0 | 0 |

| UNKNOWNS 2019 II | *Aedes cantans* (n = 6) | | *Culiseta annulata* (n = 13) | | *Aedes detritus* (n = 21) | | *Aedes punctor* (n = 11) | | *Aedes rusticus* (n = 14) | |
|---|---|---|---|---|---|---|---|---|---|---|
| | correct | wrong | correct | wrong | correct | wrong | correct | wrong | correct | wrong |
| Percentage of all samples [%] | 100 | 0 | 92.3 | 7.7 | 95.2 | 4.8 | 90.9 | 100 | 100 | 0 |
| Probability of correctness > 80 % | 83.3 | 0 | 83.3 | 0 | 95 | 100 | 100 | 100 | 100 | 0 |
| Probability of correctness < 80 % | 16.7 | 0 | 16.7 | 100 | 5 | 0 | 0 | 0 | 0 | 0 |
| Average probability of correctness | 91.3 | 0 | 96.7 | 76.1 | 97.8 | 99 | 99.1 | 98.4 | 98.8 | 0 |
| Number of outliers (StDev > 10) | 0 | 0 | 0 | 0 | 0 | 0 | 0 | 0 | 0 | 0 |

| UNKNOWNS 2019 III | *Aedes caspius* (n = 18) | | *Culiseta annulata* (n = 3) | | *Aedes detritus* (n = 6) | | *Culex pipiens* (n = 39) | |
|---|---|---|---|---|---|---|---|---|
| | correct | wrong | correct | wrong | correct | wrong | correct | wrong |
| Percentage of all samples [%] | 100 | 0 | 33.3 | 66.7 | 50 | 50 | 97.4 | 2.6 |
| Probability of correctness > 80 % | 100 | 0 | 100 | 0 | 66.7 | 100 | 92.1 | 100 |
| Probability of correctness < 80 % | 0 | 0 | 0 | 100 | 33.3 | 0 | 7.9 | 0 |
| Average probability of correctness | 98.5 | 0 | 97.4 | 74 | 85.6 | 89.8 | 94.8 | 90.1 |
| Number of outliers (StDev > 10) | 0 | 0 | 0 | 0 | 0 | 0 | 0 | 0 |

I … samples were analysed at the same time as samples used for model building
II … samples were analysed within weeks of model samples
III … samples were analysed > 2 months later than model samples

**Supplemental Figure 12:**

*The seven species PC-LDA model built in Offline Model Builder with 100 PCs and exported to the Recognition software, was used to identify wild mosquitoes caught in traps (TRAPPED). Samples were categorised depending on their time point of analysis : samples which had been analysed at the same time as samples used for model building (I), samples analysed on other days than model samples, but same time period (II) and samples which were analysed over two months later than the last samples included in the model building (a). The number of tested samples is stated in brackets. The percentage of correctly identified samples and the likelihood that the identification is correct are highlighted in yellow for easier comparison. A more detailed view of the identification results and number of samples per species is listed under b.*

a

| TRAPPED | **I** (n = 38 ) | | **II** (n = 66 ) | | **III** (n = 144) | |
|---|---|---|---|---|---|---|
| | correct | wrong | correct | wrong | correct | wrong |
| Percentage of all samples [%] | 86.8 | 13.2 | 90.9 | 9.1 | 52.1 | 47.9 |
| Probability of correctness > 80 % | 93.9 | 40 | 95 | 100 | 84 | 66.7 |
| Probability of correctness < 80 % | 6.1 | 60 | 5 | 0 | 16 | 33.3 |
| Average probability of correctness | 94.2 | 82 | 96 | 90.6 | 90 | 82.9 |
| Number of outliers (StDev > 10) | 0 | 0 | 0 | 0 | 0 | 0 |

| | |
|---|---|
| Total number of samples (n) | 248 |
| Percentage of correctly identified samples [%] | 67.7 |
| Average probability of correctness [%] | 93.4 |

b

| TRAPPED I | *Aedes detritus* (n = 28) | | *Culiseta annulata* (n = 9) | | *Aedes caspius* (n = 1) | |
|---|---|---|---|---|---|---|
| | correct | wrong | correct | wrong | correct | wrong |
| Percentage of all samples [%] | 89.3 | 10.7 | 88.9 | 11.1 | 0 | 100 |
| Probability of correctness > 80 % | 96 | 66.7 | 87.5 | 0 | 0 | 0 |
| Probability of correctness < 80 % | 4 | 33.3 | 12.5 | 100 | 0 | 100 |
| Average probability of correctness | 96.2 | 89.4 | 88.2 | 66.05 | 0 | 75.82 |
| Number of outliers (StDev > 10) | 0 | 0 | 0 | 0 | 0 | 0 |

| TRAPPED II | *Aedes detritus* (n = 64) | | *Culiseta annulata* (n = 2) | |
|---|---|---|---|---|
| | correct | wrong | correct | wrong |
| Percentage of all samples [%] | 90.6 | 9.4 | 100 | 0 |
| Probability of correctness > 80 % | 94.8 | 100 | 100 | 0 |
| Probability of correctness < 80 % | 5.2 | 0 | 0 | 0 |
| Average probability of correctness | 96.1 | 90.6 | 94.5 | 0 |
| Number of outliers (StDev > 10) | 0 | 0 | 0 | 0 |

| TRAPPED III | *Culex pipiens* (n = 63) | | *Culiseta annulata* (n = 18) | | *Aedes detritus* (n = 64) | | *Aedes cantans* (n = 1) | |
|---|---|---|---|---|---|---|---|---|
| | correct | wrong | correct | wrong | correct | wrong | correct | wrong |
| Percentage of all samples [%] | 23.8 | 76.2 | 55.6 | 44.4 | 67.2 | 18.8 | 0 | 100 |
| Probability of correctness > 80 % | 66.7 | 64.6 | 90 | 50 | 86 | 83.3 | 0 | 100 |
| Probability of correctness < 80 % | 33.3 | 35.4 | 10 | 50 | 14 | 16.7 | 0 | 0 |
| Average probability of correctness | 82.6 | 81.9 | 91.7 | 79.7 | 93.3 | 89.1 | 0 | 83.24 |
| Number of outliers (StDev > 10) | 0 | 0 | 0 | 0 | 0 | 0 | 0 | 0 |

I ... samples were analysed at the same time as samples used for model building
II ... samples were analysed within weeks of model samples
III ... samples were analysed > 2 months later than model samples

**Supplemental Figure 13:**

*Larvae were collected from three pools for REIMS analysis as well as identification purposes. For species identification larvae were raised to adults; all larvae emerged as the expected species (a). The difference between the larvae of* Aedes detritus *and* Aedes punctor *is adequate to provide separation even when using unsupervised methods such as PCA. The individual differences are represented by the principal components 1 and 2. The variance in component 3, however, supports a clear clustering of samples into their respective species (b). To test separation the PC-LDA model was also built with classifications randomly assigned to samples. A comparison of the larval species model with and without correctly assigned classes can be seen in panel c.*

a

| Expected Species | Collection pool | Number of larvae kept for identification | Number of mosquitoes identified as expected species | Collection time points |
|---|---|---|---|---|
| *Aedes detritus* | PS | 162 | 162 | 2 weeks before, during and after sampling period |
| *Aedes detritus* | PP | 65 | 65 | 2 weeks before and after sampling period |
| *Aedes punctor* | Burton Woods | 86 | 86 | Test samples were kept from each sampled batch (4) |

b

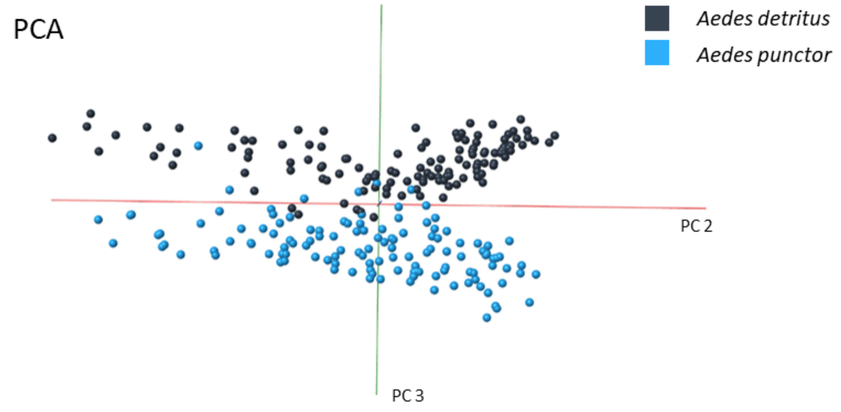


c

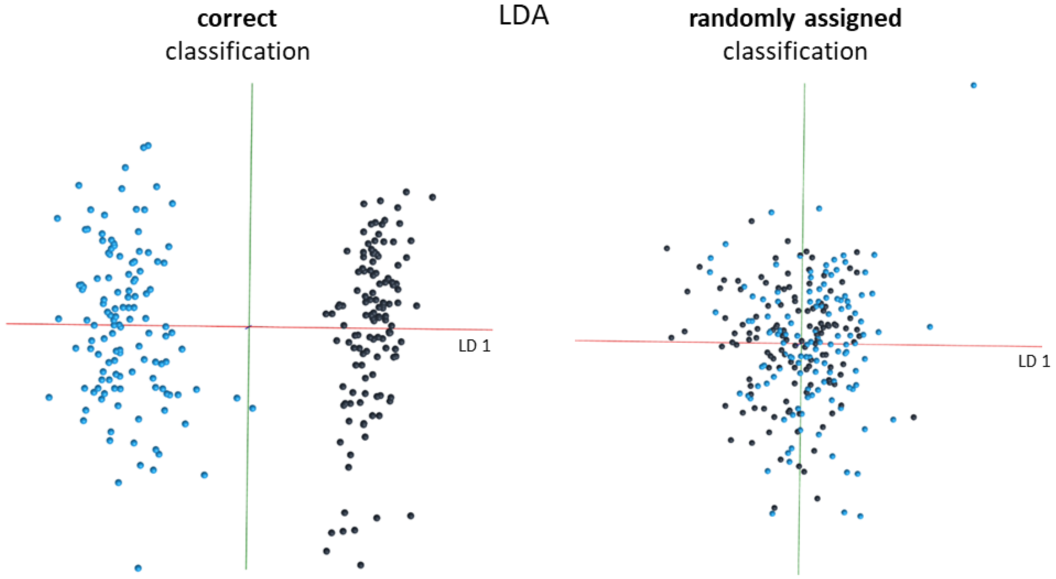

**Supplemental Figure 14:**

*Age groups were separated by PC-LD analysis and visualized using OMB (i) as well as R , in form of 3D models (using different linear discriminant combinations) (ii) and kernel density plots for each LD) using only a quarter of principal components possible. The difference between classes in model a (based on 56 PCs) decreased with the lower PC number. This is especially noticeable between groups 2 and 3, which now strongly overlap and groups 4 and 5, where samples are clustered only loosely without clear group boundaries. The young groups in model b (based on 44 PCs), moved closer to each other due to the reduction in PC numbers, however, separation is still very distinct. The main portion of the older sample classes (12+13 days) are now completely overlaid.*

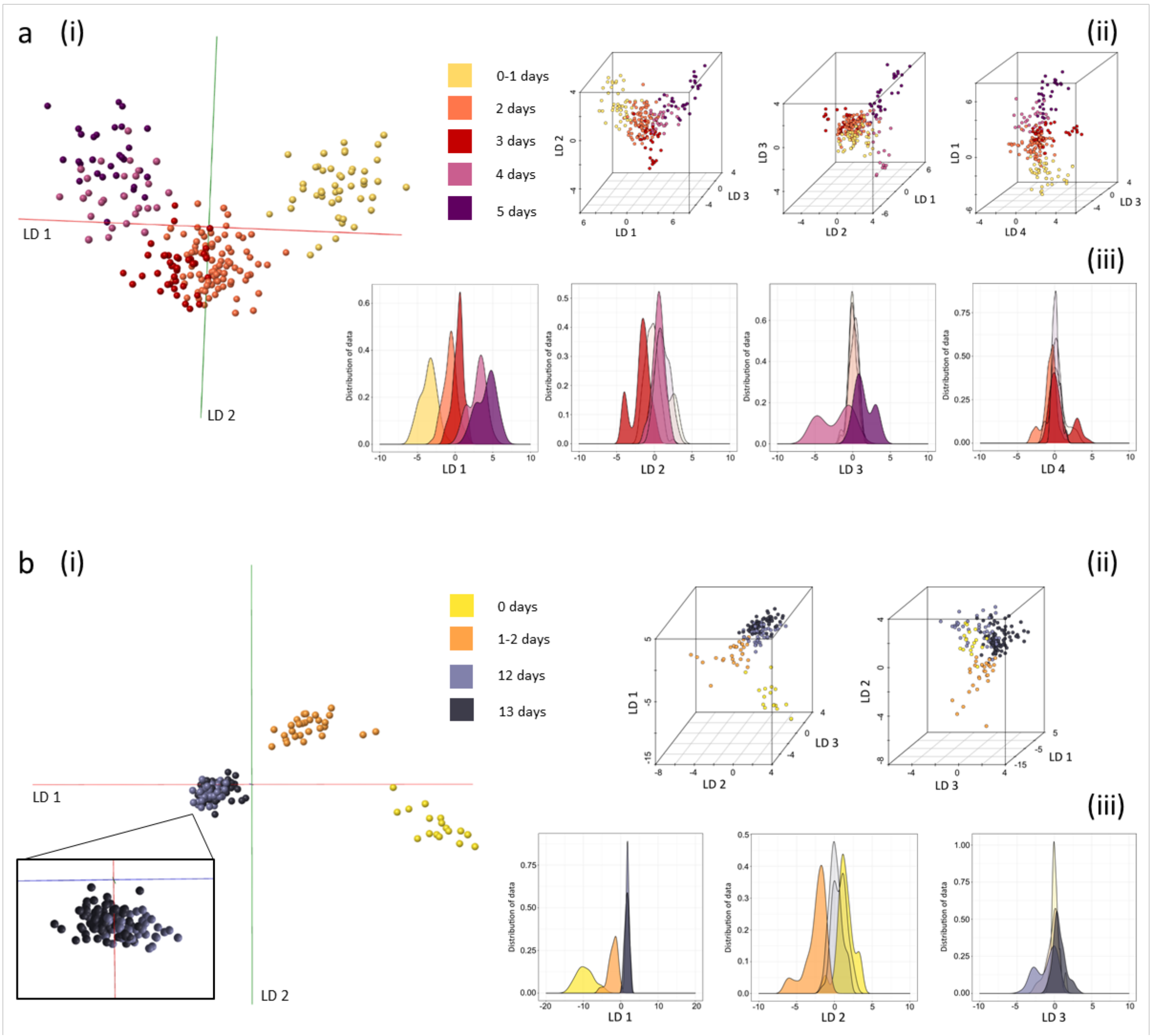

**Supplemental Figure 15:**

*To test the separation principle of model a (based on 100 PCs) and model b (based on 88 PCs) in Figure 6, classifications were randomly assigned to samples before rebuilding the models in Offline Model Builder. The original separations (left panel) can be directly compared to the randomly assigned classification models (right panel). For both models the separation following the randomisation is significantly worse with samples from the same class clustering only very loosely compared to previous grouping and significant overlap of groups.*

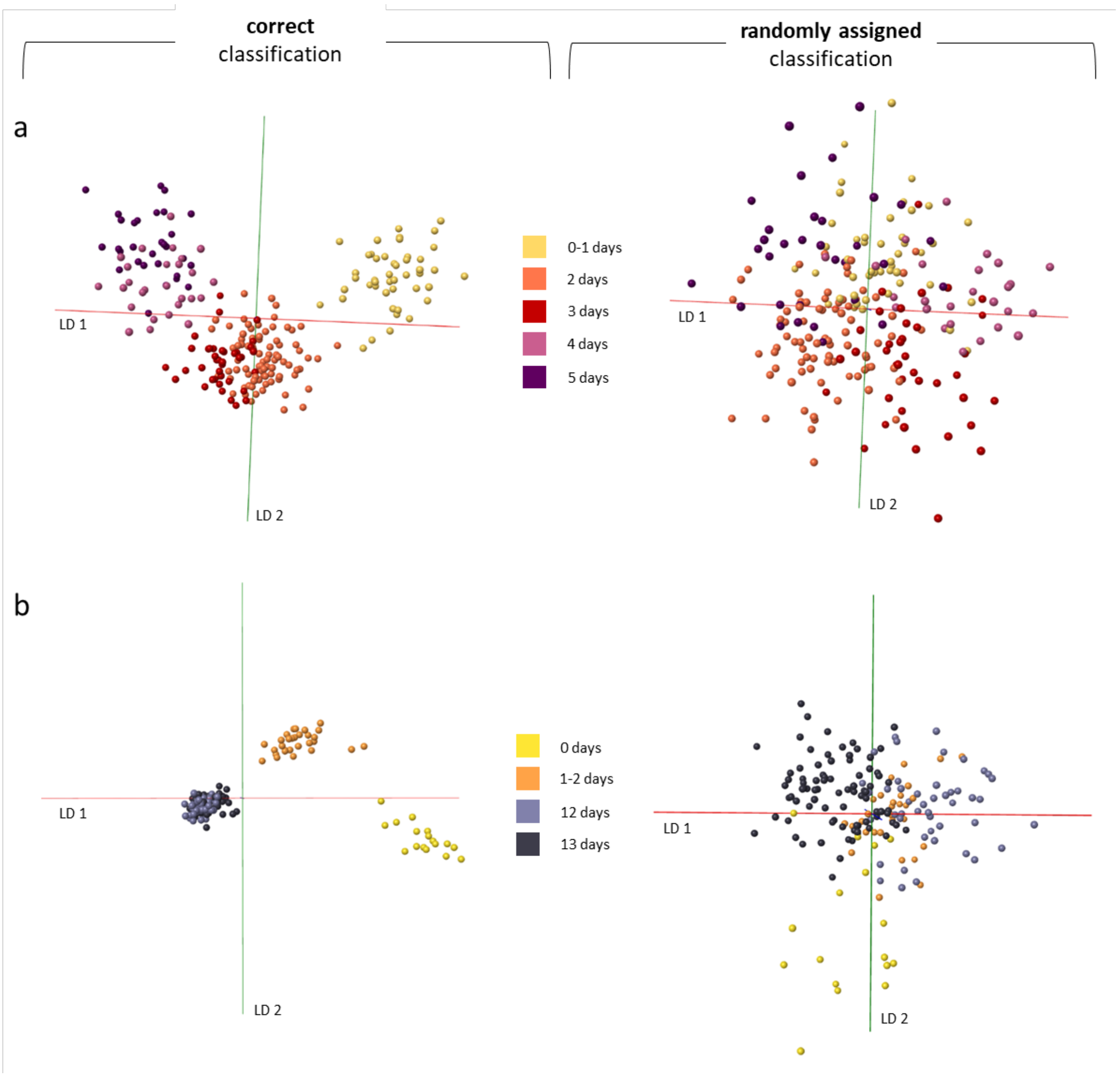

**Supplemental Figure 16:**

*The data matrix, obtained after processing and binning the mass spectral data in Offline Model Builder, was used to create averaged mass spectra for all age classes from 0-5 days. Each mass spectrum represents an average of all samples available for each age group: 0-1 day (n=47), 2 days (n=84), 3 days (n=39), 4 days (n=27), 5 days (n=30).*

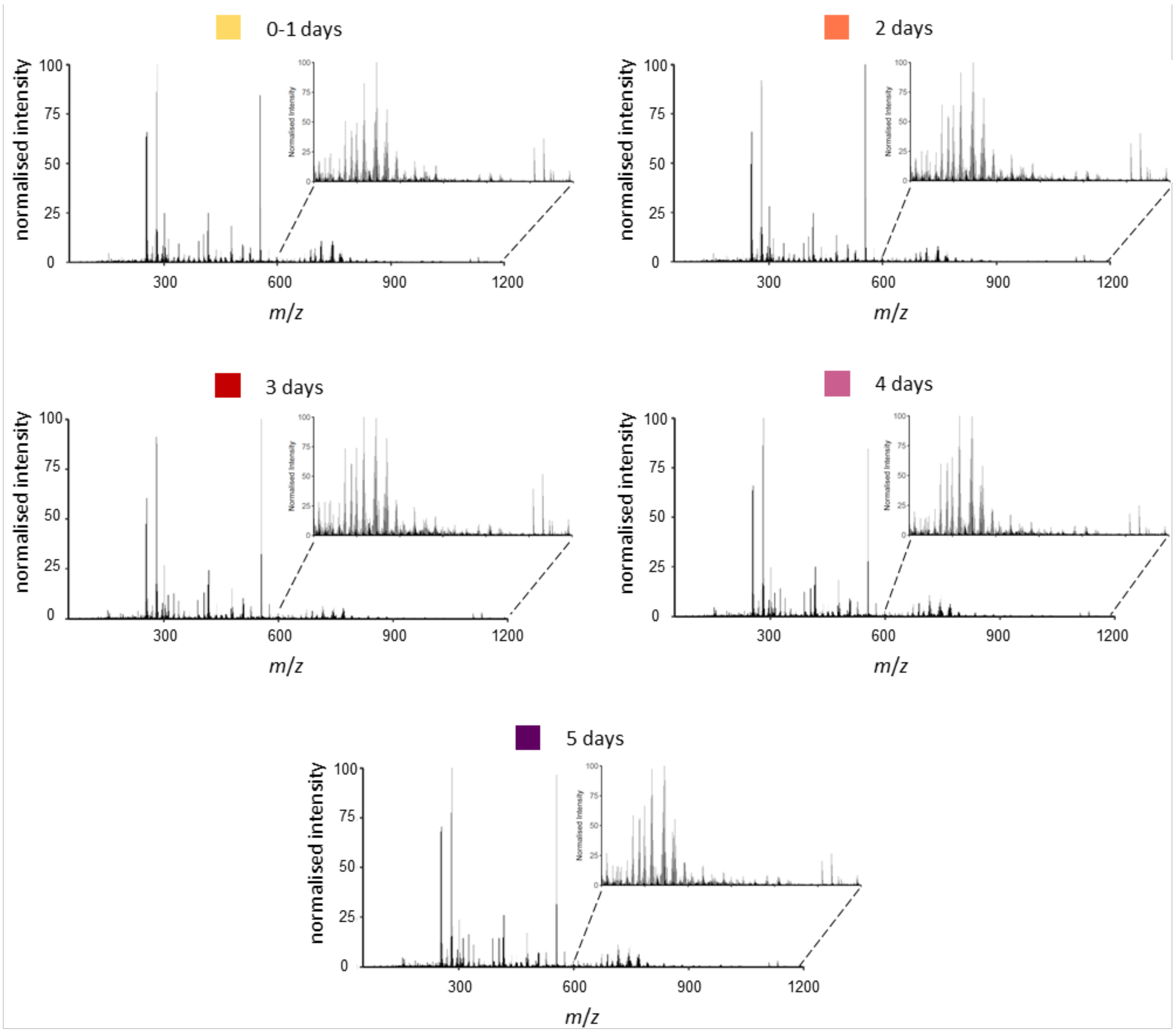

**Supplemental Figure 17:**

*To improve separation of the individual age classes for both models some age groups were combined into one class. For the model in panel a the 2 and 3 day old mosquitoes, as well as the 4 and 5 day old specimens, were combined into one group each, reducing the overall number of classes from 5 to 3 (panel a). As mosquitoes which have just emerged and 1 day old mosquitoes can be readily distinguished, only the 12 and 13 day old mosquitoes were combined into one group for the model in panel b. As with the previous age models, PC-LD analysis was conducted first in Offline Model Builder (i) to extract the data matrix, before repeating analysis in R to visualise separation results through kernel density histograms (ii) and 2D scatter plots (iii). Principal component numbers were the same as used for the previous model (model a: 100 PCs, model b: 88 and 85 PCs) to solely observe the effect of class reduction. For both models all age groups are now separated along linear discriminant one; LD 2 merely contributes additional variance to increase separation of the younger groups. There are now distinct gaps between all age classes.*

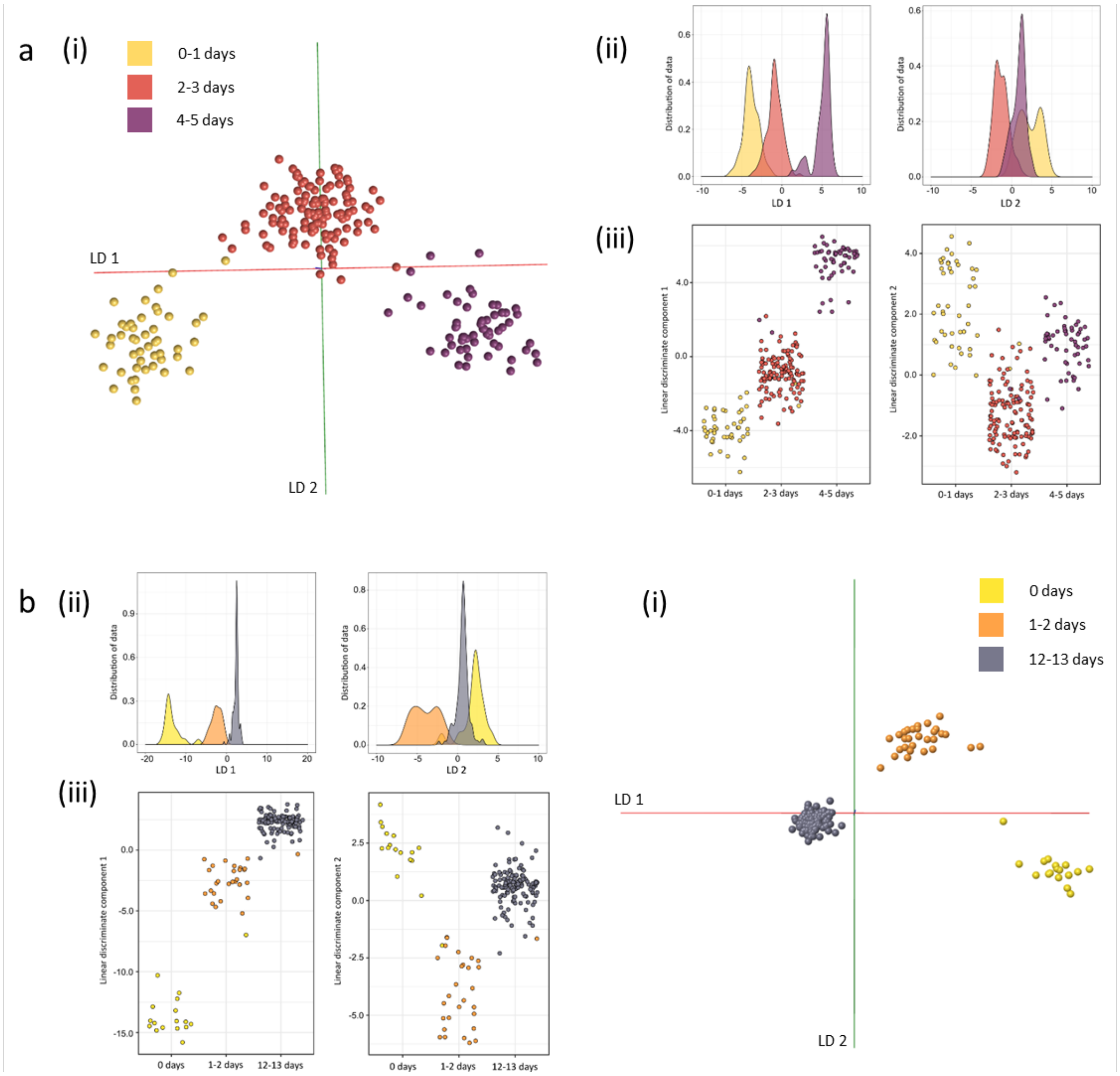

**Supplemental Figure 18:**

*The PCA-LDA based age models, one comprising 5 classes (0-5 days) and the other 4 classes (0-13 days), were cross-validated within Offline Model Builder using the setting 'Leave 20 % out' and a standard deviation of 5 (top panel, original models)). The models comprised of combined age classes were also cross-validated using the same settings (bottom panel, improved models). Combining the age classes clearly improved separation accuracy from 79 to 93 % (for the model on the left) and 74 to 100 % (for the model on the right). Some samples were not tested as 20 % of the total sample number resulted in a fractional number.*

Cross-validation *Offline Model Builder*:

ORIGINAL MODELS

| Confusion matrix | 0-1 day | 2 days | 3 days | 4 days | 5 days | Outlier |
|---|---|---|---|---|---|---|
| 0-1 day | 43 | 2 | 0 | 0 | 0 | 1 |
| 2 days | 1 | 73 | 8 | 1 | 0 | 0 |
| 3 days | 0 | 12 | 25 | 1 | 1 | 0 |
| 4 days | 0 | 0 | 5 | 17 | 5 | 0 |
| 5 days | 0 | 0 | 2 | 9 | 16 | 3 |

| Spectra (n) | Pass | Fail | Outlier | Correct Classification (%) |
|---|---|---|---|---|
| 225 | 174 | 47 | 4 | 78.73 |

| Confusion matrix | 0 days | 1-2 days | 12 days | 13 days | Outlier |
|---|---|---|---|---|---|
| 0 days | 14 | 0 | 0 | 0 | 3 |
| 1-2 days | 0 | 28 | 0 | 0 | 1 |
| 12 days | 0 | 0 | 26 | 20 | 0 |
| 13 days | 0 | 0 | 24 | 57 | 2 |

| Spectra (n) | Pass | Fail | Outlier | Correct Classification (%) |
|---|---|---|---|---|
| 175 | 125 | 44 | 6 | 73.96 |

IMPROVED MODELS

| Confusion matrix | 0-1 day | 2-3 days | 4-5 days | Outlier |
|---|---|---|---|---|
| 0-1 day | 43 | 2 | 0 | 2 |
| 2-3 days | 1 | 115 | 5 | 0 |
| 4-5 days | 0 | 8 | 49 | 0 |

| Spectra (n) | Pass | Fail | Outlier | Correct Classification (%) |
|---|---|---|---|---|
| 225 | 207 | 16 | 2 | 92.83 |

| Confusion matrix | 0 days | 1-2 days | 12-13 days | Outlier |
|---|---|---|---|---|
| 0 days | 15 | 0 | 0 | 2 |
| 1-2 days | 0 | 27 | 0 | 2 |
| 12-13 days | 0 | 0 | 129 | 0 |

| Spectra (n) | Pass | Fail | Outlier | Correct Classification (%) |
|---|---|---|---|---|
| 175 | 171 | 0 | 4 | 100 |

**Supplemental Figure 19:**

*The Anopheles species model and age model (both based on 100 PCs) were cross-validated within Offline Model Builder using the setting 'Leave 20 % out' and a standard deviation of 5.*

Cross-validation *Offline Model Builder*:

| Confusion matrix | An. gambiae | An. arabiensis | An. coluzzii | Outlier |
|---|---|---|---|---|
| An. gambiae | 177 | 1 | 2 | 1 |
| An. arabiensis | 1 | 177 | 2 | 1 |
| An. coluzzii | 4 | 2 | 174 | 1 |

| Spectra (n) | Pass | Fail | Outlier | Correct Classification (%) |
|---|---|---|---|---|
| 540 | 528 | 12 | 0 | 97.78 |

| Confusion matrix | 1 day | 5-6 days | 14-15 days | Outlier |
|---|---|---|---|---|
| 1 day | 180 | 0 | 0 | 0 |
| 5-6 days | 0 | 178 | 1 | 1 |
| 14-15 days | 0 | 2 | 178 | 0 |

| Spectra (n) | Pass | Fail | Outlier | Correct Classification (%) |
|---|---|---|---|---|
| 540 | 536 | 3 | 1 | 99.44 |

**Supplemental Figure 20:**

*The PCA-LDA models separating* Anopheles *mosquitoes by species and age were re-built in R using a lower number of principal components. The separation depicted in the kernel density histograms and scatter plots is based on 135 PCs (¼ of max) for both models.*

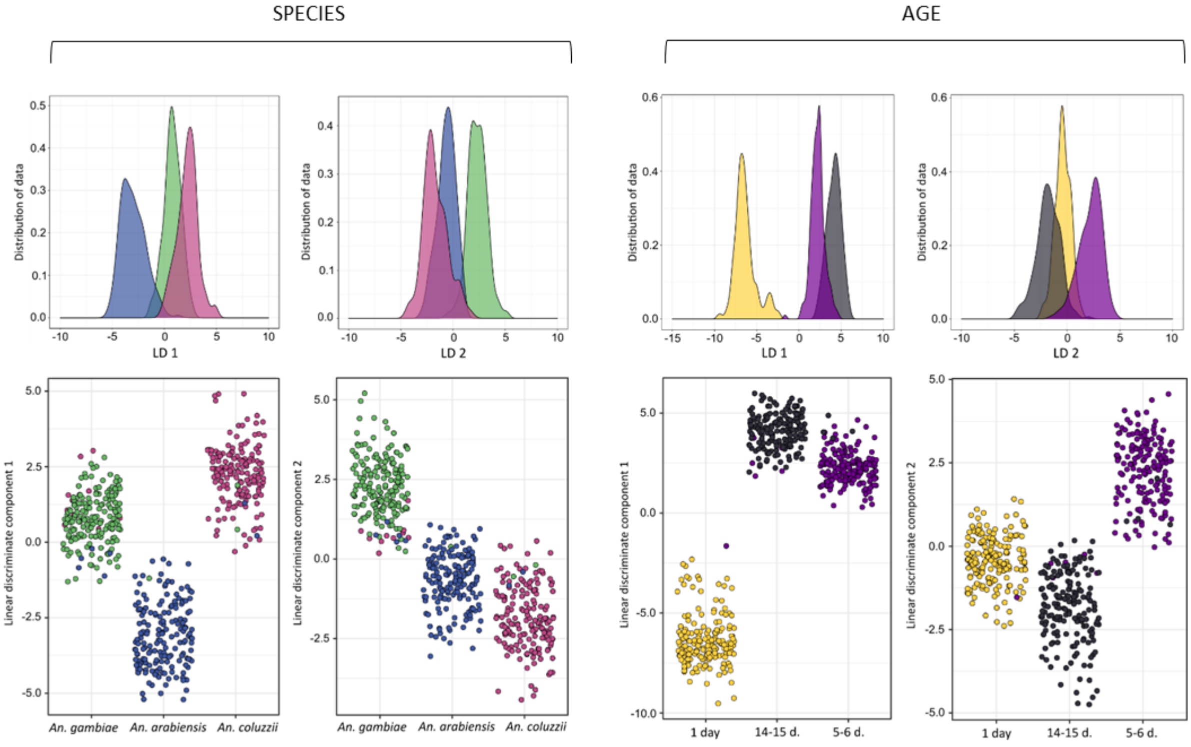

**Supplemental Figure 21:**

*A list of the variables identified as important for the random forest based separation of the three mosquito species An. gambiae, An. arabiensis and An. coluzzii (a) as well as the three age groups 1 day, 5-6 days and 14-15 days (b). Only the intensities of ion bins, which have been in the Top 10 variables list in at least 7 out of 10 random forest runs, are plotted. Although some of the ion bins had not been identified as very important in every run, they nevertheless play an important role in the separation process. The m/z bins 580.5, 683.5 and 808.5 appear to support the separation of Moz from the other two classes, which was not achieved with the variables identified in every run (100 %). The ion bins which are driving age separation 100 % of the time, however, already provide enough variance to separate all three groups; the other two bins (510.5 and 283.3) merely add further variance to the process.*

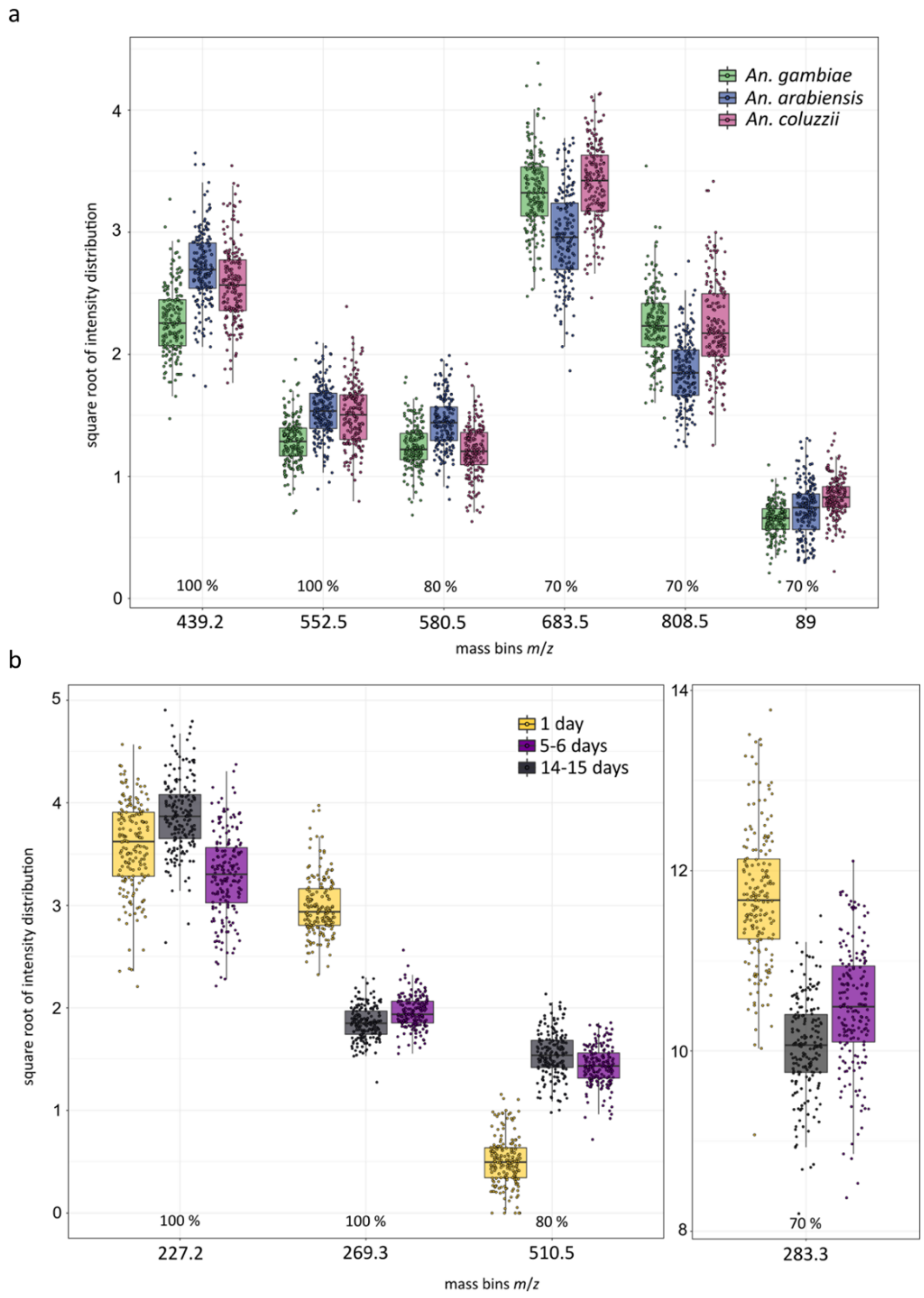

**Supplemental Figure 22:**

*The nine-class species/age model (LDA based on 100 PCs) was cross-validated within Offline Model Builder using the setting 'Leave 20 % out' and a standard deviation of 5.*

Cross-validation *Offline Model Builder*:

| Confusion matrix | *An. gambiae* 1 day | *An. gambiae* 5-6 days | *An. gambiae* 14-15 days | *An. arabiensis* 1 day | *An. arabiensis* 5-6 days | *An. arabiensis* 14-15 days | *An. coluzzii* 1 day | *An. coluzzii* 5-6 days | *An. coluzzii* 14-15 days | Outlier |
|---|---|---|---|---|---|---|---|---|---|---|
| *An. gambiae* 1 day | 60 | 0 | 0 | 0 | 0 | 0 | 0 | 0 | 0 | 0 |
| *An. gambiae* 5-6 days | 0 | 58 | 2 | 0 | 0 | 0 | 0 | 0 | 0 | 0 |
| *An. gambiae* 14-15 days | 0 | 0 | 58 | 0 | 0 | 0 | 0 | 0 | 1 | 1 |
| *An. arabiensis* 1 day | 0 | 0 | 0 | 59 | 0 | 0 | 0 | 0 | 0 | 1 |
| *An. arabiensis* 5-6 days | 0 | 0 | 0 | 0 | 58 | 0 | 0 | 2 | 0 | 0 |
| *An. arabiensis* 14-15 days | 0 | 0 | 0 | 0 | 0 | 60 | 0 | 0 | 0 | 0 |
| *An. coluzzii* 1 day | 1 | 0 | 0 | 0 | 0 | 0 | 57 | 0 | 0 | 2 |
| *An. coluzzii* 5-6 days | 0 | 2 | 0 | 0 | 2 | 0 | 0 | 56 | 0 | 0 |
| *An. coluzzii* 14-15 days | 0 | 0 | 4 | 0 | 0 | 1 | 0 | 0 | 55 | 0 |

| Spectra (n) | Pass | Fail | Outlier | Correct Classification (%) |
|---|---|---|---|---|
| 540 | 521 | 15 | 4 | 97.20 |

**Supplemental Figure 23:**

*The data matrix from the nine-class species/age model was used for random forest analysis, which was repeated 10 times, using different randomly selected training (70 % of the data) and test (30 % of the data) data sets. The confusion matrix contains the mean percentages of correctly identified and misidentified samples for every species as well as the standard error of the mean. The range of classification accuracy achieved for each of the 10 models (lowest and highest percentage) is listed in parentheses below the standard error of the mean. The average number of samples per class used for testing the model are listed on the side (n = x). The overall model accuracy was 79 ± 1.4 % (mean ± SEM).*

Predicted classification

Correct classification

| | *An. coluzzii* 1 day | *An. coluzzii* 5-6 days | *An. coluzzii* 14-15 days | *An. gambiae* 1 day | *An. gambiae* 5-6 days | *An. gambiae* 14-15 days | *An. arabiensis* 1 day | *An. arabiensis* 5-6 days | *An. arabiensis* 14-15 days | n |
|---|---|---|---|---|---|---|---|---|---|---|
| *An. coluzzii* 1 day | **84** ± 4.6 (63-100) | 0 | 0 | 15 | 0 | 0 | 1 | 0 | 0 | 20 |
| *An. coluzzii* 5-6 days | 0 | **71** ± 3.1 (57-92) | 4 | 0 | 9 | 1 | 1 | 13 | 1 | 18 |
| *An. coluzzii* 14-15 days | 0 | 2 | **77** ± 3.7 (60-94) | 0 | 1 | 10 | 0 | 2 | 7 | 18 |
| *An. gambiae* 1 day | 8 | 0 | 0 | **92** ± 1.7 (83-100) | 0 | 0 | 0 | 0 | 0 | 19 |
| *An. gambiae* 5-6 days | 0 | 3 | 3 | 0 | **75** ± 4.4 (47-100) | 12 | 1 | 5 | 0 | 17 |
| *An. gambiae* 14-15 days | 0 | 1 | 5 | 0 | 11 | **74** ± 2.9 (56-88) | 0 | 4 | 4 | 20 |
| *An. arabiensis* 1 day | 0 | 0 | 0 | 3 | 0 | 0 | **97** ± 1.7 (83-100) | 0 | 0 | 17 |
| *An. arabiensis* 5-6 days | 0 | 11 | 1 | 0 | 7 | 0 | 0 | **76** ± 3.4 (57-94) | 5 | 17 |
| *An. arabiensis* 14-15 days | 0 | 5 | 9 | 0 | 4 | 8 | 0 | 4 | **70** ± 2.7 (62-88) | 17 |

**Model accuracy: 79 %** ± 1.4

**Supplemental Figure 24:**

*After building the nine-class model in Offline Model Builder, the data matrix was exported to repeat PC-LD analysis in R. The kernel density plot (LDA was based on 235 PCs) demonstrates an age related separation along LD1 (a), quite similar to what was observed in the Offline Model Builder result. Again, to simplify visualisation, the 1 day old mosquitoes were removed from the data set and the PC-LDA (180 PCs) separation process further examined using 3D scatter plots (b). While having a similar distribution of variance across the linear discriminants as seen in the Offline Model Builder model, the separation seems less defined and the class '*An. arabiensis *14-15 days' is now separated along LD1, together with the age clusters, instead of LD 2. Nevertheless, separation of groups due to age seems to happen along LD1,* An. arabiensis *is separated along LD1 and LD 2 and to distinguish* An. coluzzii *and* An. gambiae *groups LD 3 is needed. Interestingly, separation of differently aged mosquitoes is easier with* An. coluzzii *than with* An. gambiae *specimens. Plotting the outcomes of PC-LD analysis as a matrix table reveals that the six classes are very well separated (c).*

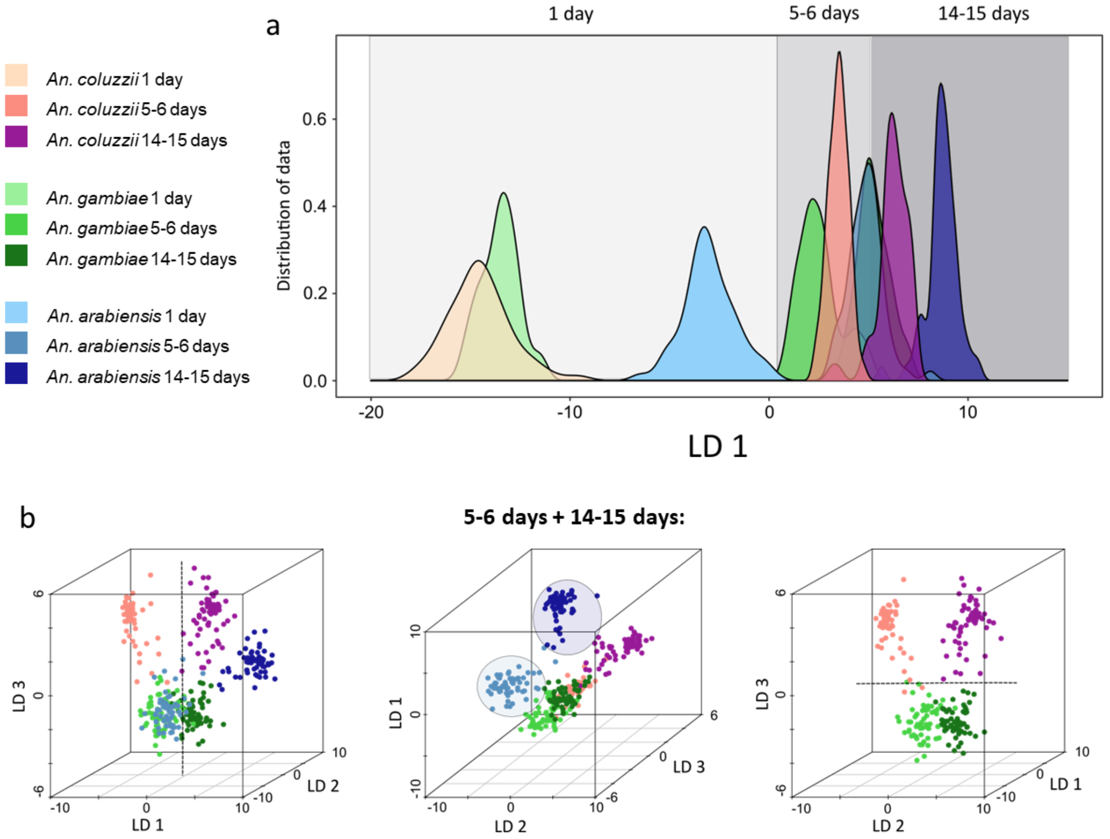


c

| PCA-LDA | *An. gambiae* 5-6 days | *An. gambiae* 14-15 days | *An. arabiensis* 5-6 days | *An. arabiensis* 14-15 days | *An. coluzzii* 5-6 days | *An. coluzzii* 14-15 days | n |
|---|---|---|---|---|---|---|---|
| *An. gambiae* 5-6 days | 58 | 2 | 0 | 0 | 0 | 0 | 60 |
| *An. gambiae* 14-15 days | 0 | 60 | 0 | 0 | 0 | 0 | 60 |
| *An. arabiensis* 5-6 days | 0 | 0 | 60 | 0 | 0 | 0 | 60 |
| *An. arabiensis* 14-15 days | 0 | 0 | 0 | 60 | 0 | 0 | 60 |
| *An. coluzzii* 5-6 days | 4 | 0 | 0 | 0 | 56 | 0 | 60 |
| *An. coluzzii* 14-15 days | 1 | 0 | 0 | 0 | 0 | 59 | 60 |

**Supplemental Figure 25:**

*The top 10 most important variables were collated from ten repeated random forest analyses of the two-factor species/age model. Variables which had been identified as separation drivers in more than half the runs were selected to have their intensities plotted. The first two variables, identified 100 % of the time, m/z 227.2 and 269.3 had also been identified in the age model as important separators. The fact that they have also been identified in the nine-class model, in all 10 runs, confirms their importance for age separation. One of the two main separators of the species model, m/z 439.2, also features in this model's variable list. The other two variables 685.5 and 836.5 have not been identified and seem to be uniquely important for this two-factor model, separating the 1-day old classes of the three species.*

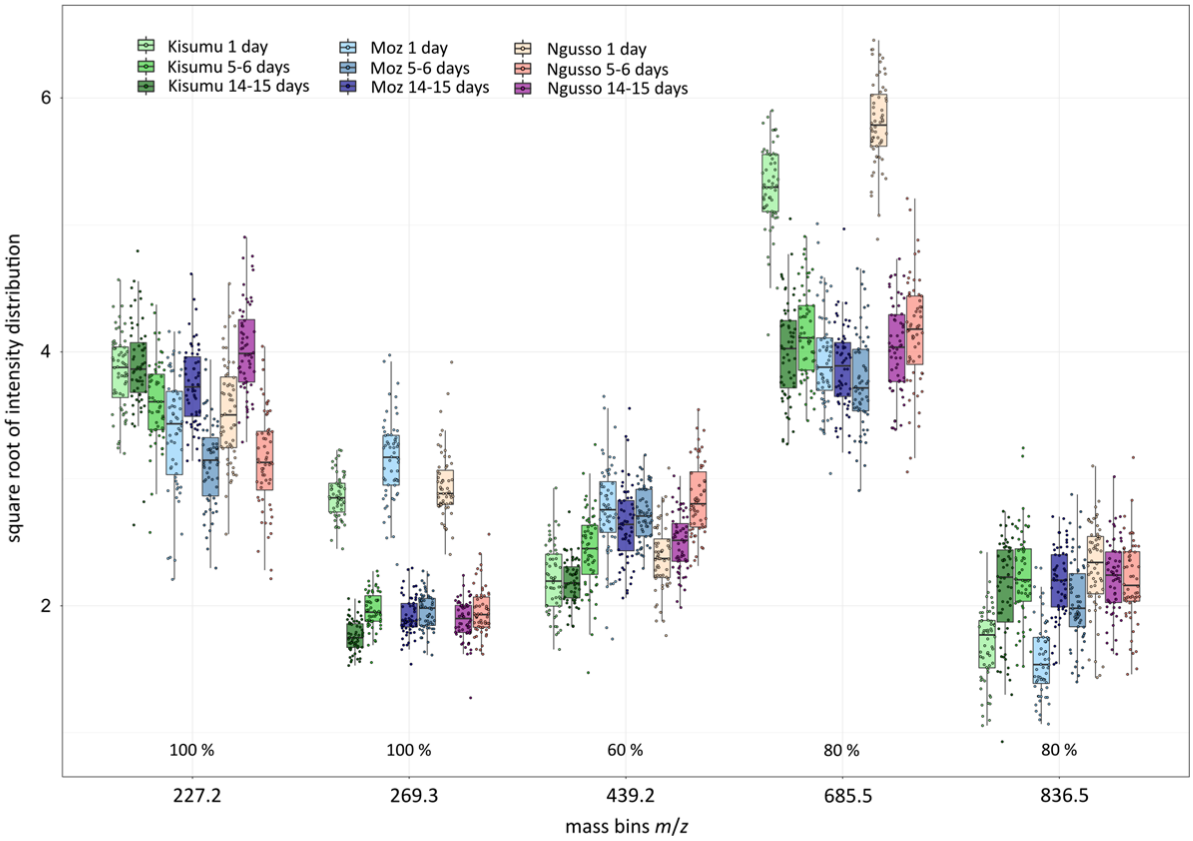

**Supplemental Figure 26:**

*After using 540 Anopheles mosquito specimens, from three species and three age groups each, to build models separating species, age as well as both properties at once, models were re-built with randomly assigned classifications. When re-building the PC-LDA models in Offline Model Builder with classes randomly assigned to samples, separation failed for all three models.*

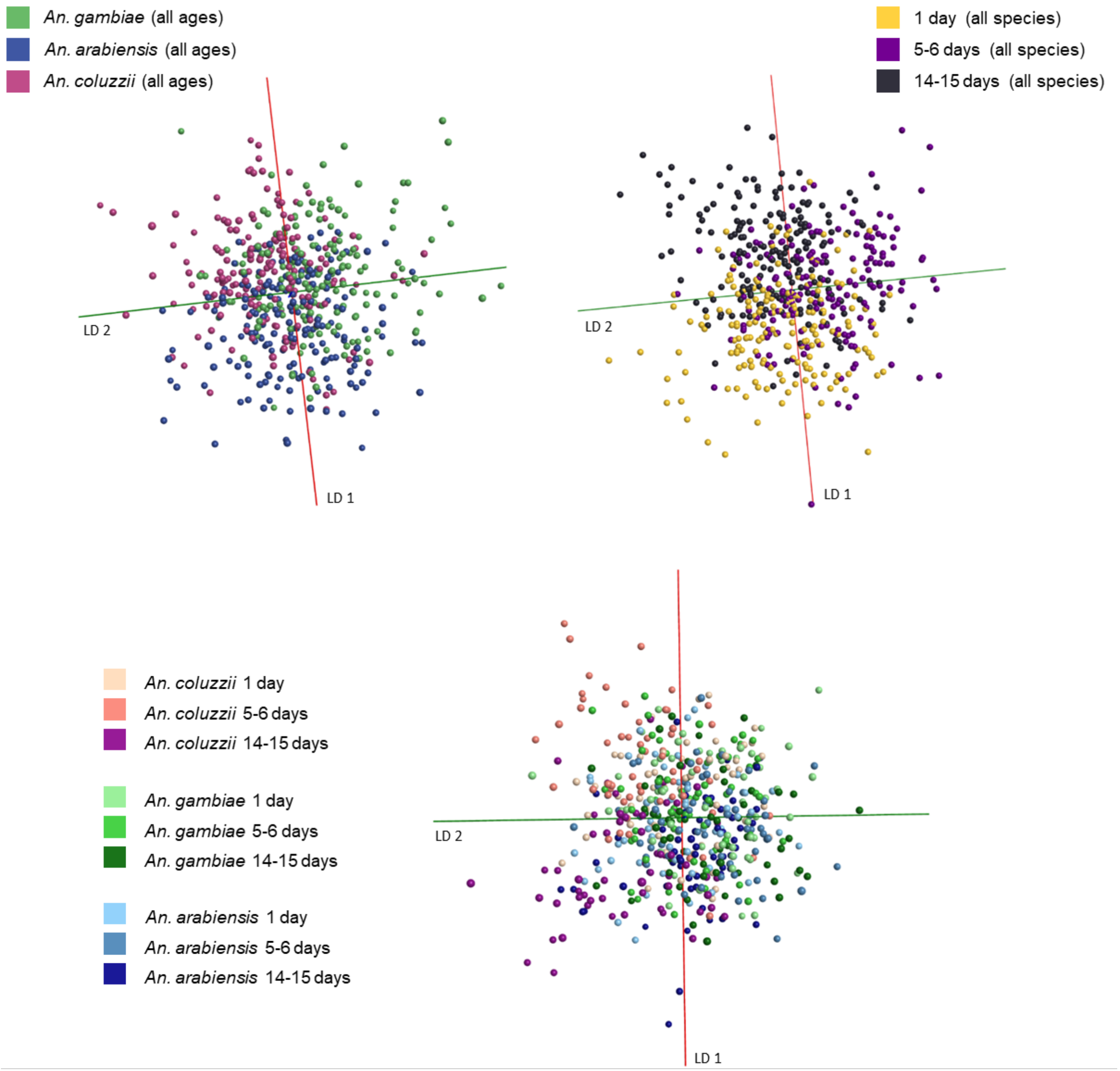

**Supplemental Figure 27:**

*Samples from 4 species (*Aedes detritus, Culiseta annulata, Aedes rusticus, Aedes punctor*) are included in these age models, separating age groups between 0 and 4 days. Separation is demonstrated using four adjacent age groups (panel a), as well as 2 groups separated by a 24 h gap (panel b). First models were built within OMB using PC-LDA (i), before exporting the matrix and conducting PC-LDA in R, depicted in form of kernel density plots (ii) and scatter plots - 3D and 2D (iii). The age model based on two age groups promised sufficient separation to be used for classification and was therefore additionally analysed via random forest (iv), using 70 % of samples for model building and 30 % for testing (1200 trees). The random forest result is presented in two bars stating the correct classification percentage, including SEM value and the range of achieved accuracies in 10 runs (min and max), and the percentage of misclassified test samples. Samples numbers used for model in panel a: 0-24 h (108), 25-48 h (55), 49-72 h (23), 73-96 h (40). Sample numbers used for model in panel b: 0-24 h (65), 49-96 (63); sample numbers from 0-24 h were reduced for random forest analysis. Separation in panel a were based on 100 (OMB) and 130 PCs (R). Separation in panel b were based on 60 (OMB) and 65 PCs (R).*

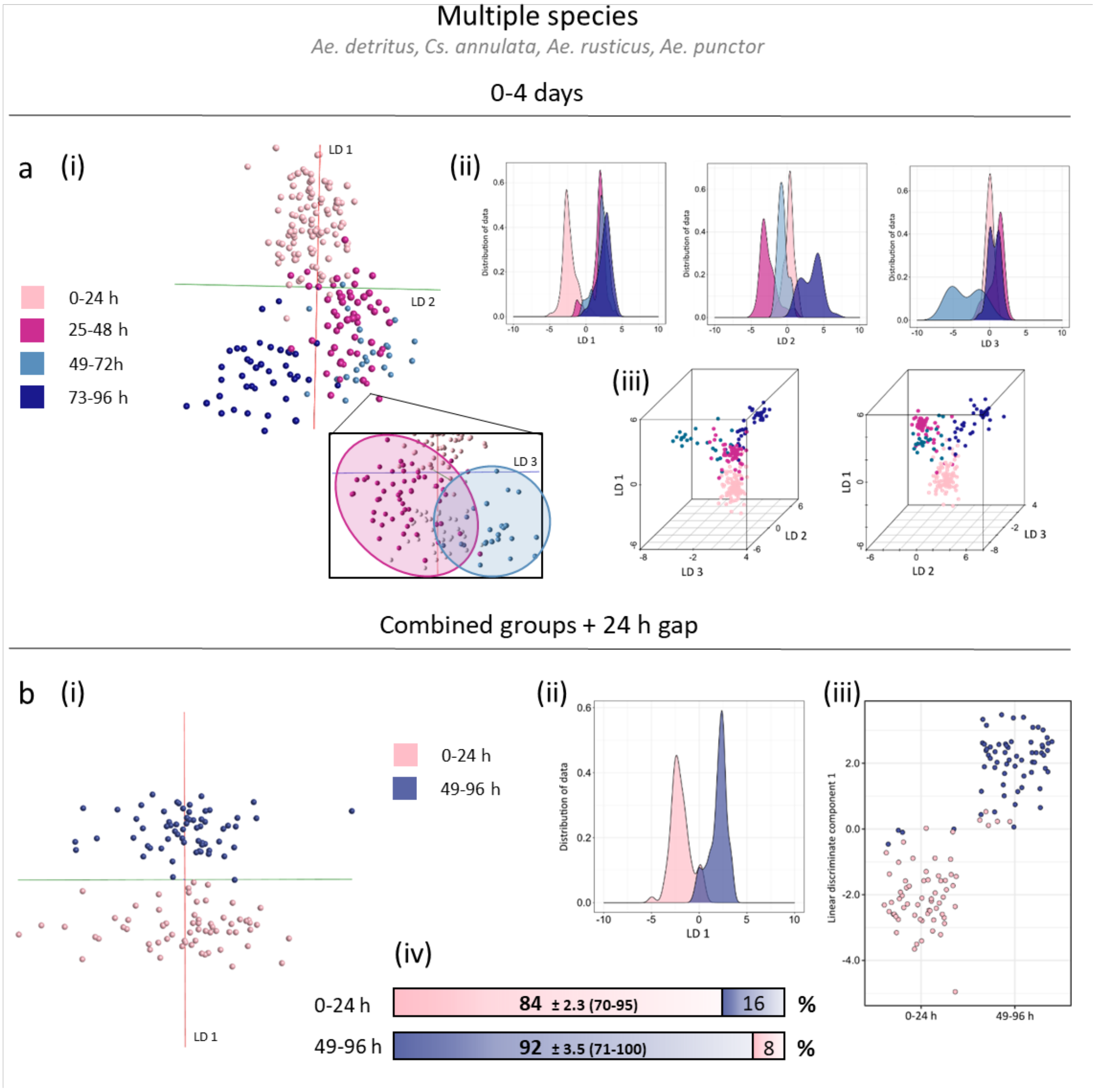

**Supplemental Figure 28:**

*Original and improved age models including only Aedes detritus specimens. The original age model (a) comprises four consecutive age groups demonstrating separation of calendar days. Due to the continuous nature of these classes, separation accuracy is low. Combination of groups (b) reduces the overall class overlap in the model, subsequently improving separation efficiency. Introduction of a 24 h gap between age groups (c) helps to enhance the difference between mosquitoes of different ages even further. All results are based one PC-LD analysis, depicted in form of OMB models and kernel density and scatter plots produced in R (from left to right). The correct classification rates, achieved through 'Leave 20 % out' cross-validation in OMB, are highlighted in yellow for each model. The number of samples per class are listed in brackets after the age information.*

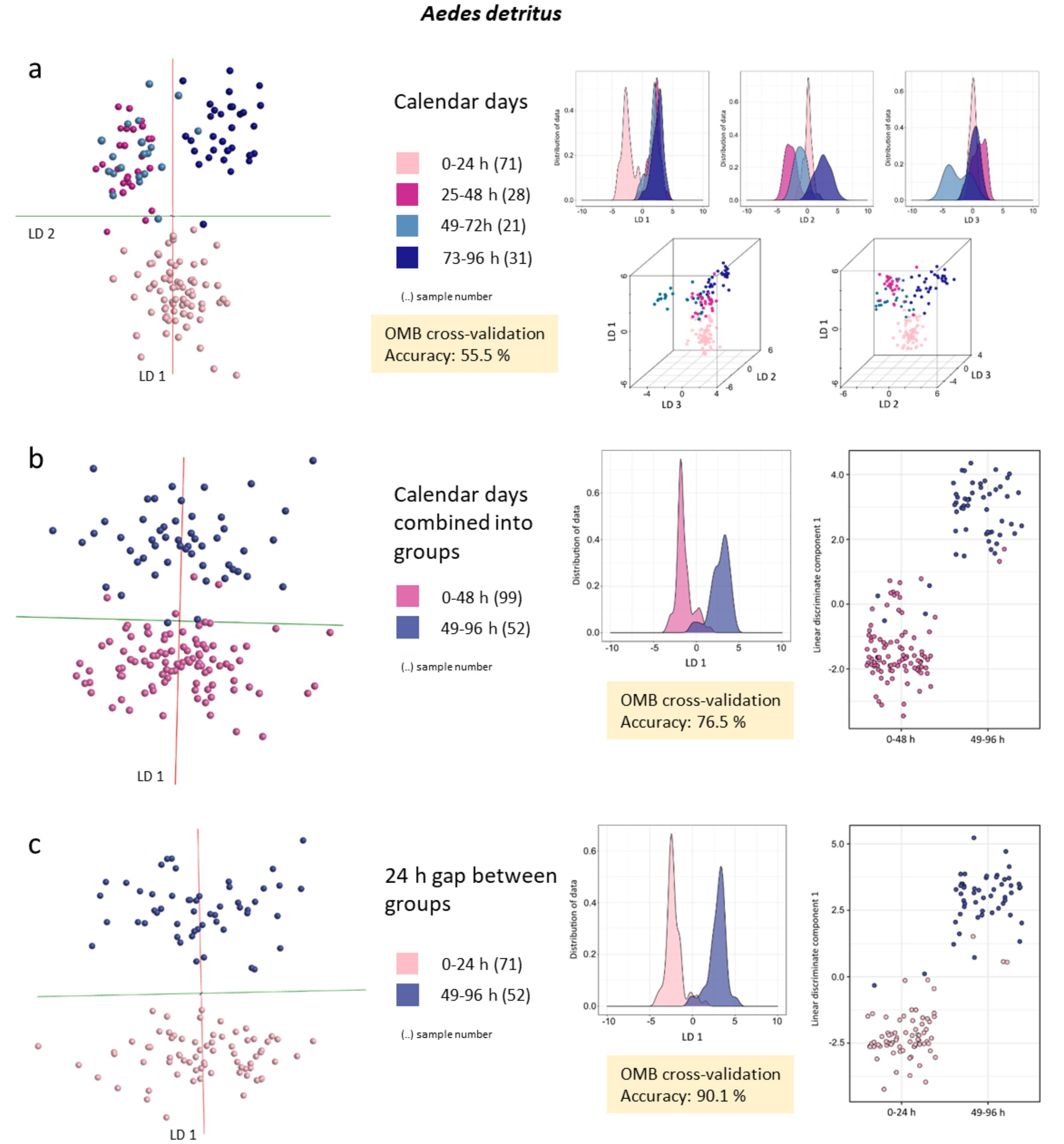

**Supplemental Figure 29:**

*The three Aedes detritus age models were tested via cross-validation in OMB using the option 'Leave out 20 %' and a standard deviation of 5. The number of principal components used for model building are given in brackets underneath the tables. One sample each was left out from the first two models as 20 % of 151 samples results in a fractional number that is rounded to the nearest integer.*

### Calendar days

| Confusion matrix | 0-24 h | 25-48 h | 49-72 h | 73-96 h | Outlier |
|---|---|---|---|---|---|
| 0-24 h | 50 | 7 | 4 | 6 | 3 |
| 25-48 h | 5 | 8 | 11 | 4 | 0 |
| 49-72 h | 2 | 9 | 6 | 4 | 0 |
| 73-96 h | 1 | 4 | 8 | 17 | 1 |

| Spectra (n) | Pass | Fail | Outlier | Correct Classification (%) |
|---|---|---|---|---|
| 150 | 81 | 65 | 4 | 55.48 |

(75 PCs)

### Calendar days combined into groups

| Confusion matrix | 0-48 h | 49-96 h | Outlier |
|---|---|---|---|
| 0-48 h | 77 | 21 | 1 |
| 49-96 h | 14 | 37 | 0 |

| Spectra (n) | Pass | Fail | Outlier | Correct Classification (%) |
|---|---|---|---|---|
| 150 | 114 | 35 | 1 | 76.51 |

(90 PCs)

### 24 h gap between groups

| Confusion matrix | 0-24 h | 49-96 h | Outlier |
|---|---|---|---|
| 0-24 h | 63 | 7 | 1 |
| 49-96 h | 5 | 46 | 1 |

| Spectra (n) | Pass | Fail | Outlier | Correct Classification (%) |
|---|---|---|---|---|
| 123 | 109 | 12 | 2 | 90.08 |

(70 PCs)

**Supplemental Figure 30:**

*The original and improved age models including specimens from four species: Aedes detritus, Culiseta annulata, Aedes rusticus and Aedes punctor. The original age model (a) comprises four consecutive age groups demonstrating separation of calendar days. Due to the continuous nature of these classes, separation accuracy is low. Combination of groups (b) reduces the overall class overlap in the model, subsequently improving separation efficiency. Introduction of a 24 h gap between age groups (c) helps to enhance the difference between mosquitoes of different ages even further. All results are based one PC-LD analysis, depicted in form of OMB models and kernel density and scatter plots produced in R (from left to right). The correct classification rates, achieved through 'Leave 20 % out' cross-validation in OMB, are highlighted in yellow for each model. The number of samples per class are listed in brackets after the age information.*

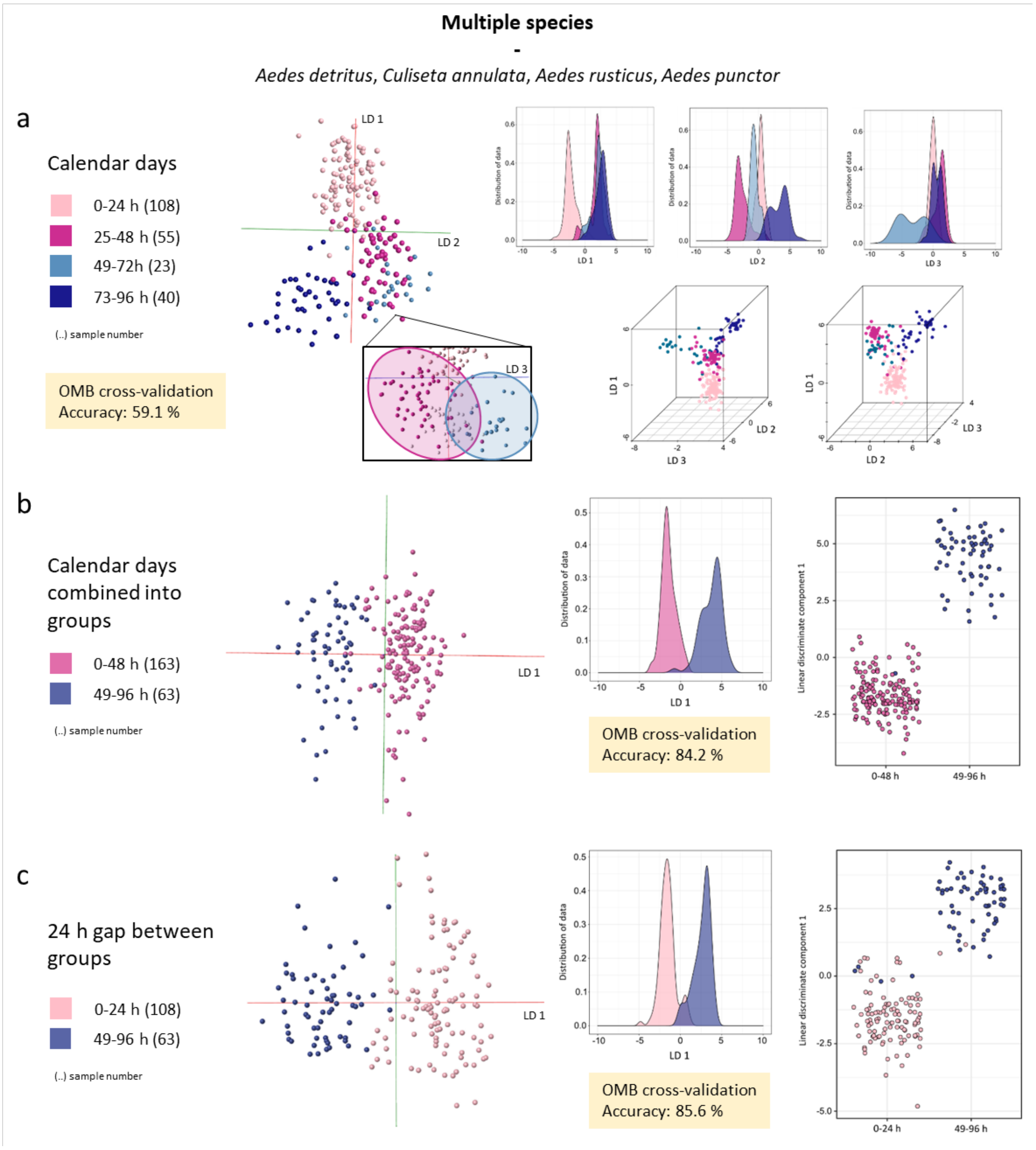

**Supplemental Figure 31:**

*The three multi-species age models were tested via cross-validation in OMB using the option 'Leave out 20 %' and a standard deviation of 5. The number of principal components used for model building are given in brackets underneath the tables. One sample each was left out from all models as 20 % of 226 and 171 samples results in fractional numbers that are rounded to the nearest integer.*

### Calendar days

| Confusion matrix | 0-24 h | 25-48 h | 49-72 h | 73-96 h | Outlier |
|---|---|---|---|---|---|
| 0-24 h | 77 | 16 | 1 | 10 | 3 |
| 25-48 h | 15 | 22 | 9 | 8 | 1 |
| 49-72 h | 3 | 6 | 10 | 4 | 0 |
| 73-96 h | 3 | 7 | 8 | 21 | 1 |

| Spectra (n) | Pass | Fail | Outlier | Correct Classification (%) |
|---|---|---|---|---|
| 225 | 130 | 90 | 5 | 59.09 |

(100 PCs)

### Calendar days combined into groups

| Confusion matrix | 0-48 h | 49-96 h | Outlier |
|---|---|---|---|
| 0-48 h | 140 | 20 | 2 |
| 49-96 h | 15 | 47 | 1 |

| Spectra (n) | Pass | Fail | Outlier | Correct Classification (%) |
|---|---|---|---|---|
| 225 | 187 | 35 | 3 | 84.23 |

(90 PCs)

### 24 h gap between groups

| Confusion matrix | 0-24 h | 49-96 h | Outlier |
|---|---|---|---|
| 0-24 h | 90 | 14 | 3 |
| 49-96 h | 10 | 53 | 0 |

| Spectra (n) | Pass | Fail | Outlier | Correct Classification (%) |
|---|---|---|---|---|
| 170 | 143 | 24 | 3 | 85.63 |

(70 PCs)

**Supplemental Figure 32:**

*All age models with a 24 h gap between age classes, one* Aedes detritus *and two multi-species ones, were rebuilt using randomly assigned classifications. The PC-LDA based models built with correct (left) and randomly assigned classifications (right) are listed for comparison. Randomly assigned classifications lead to a considerably worse separation, with individual samples being scattered and classes overlapping. The number of principal components and other settings used for model building were identical for both approaches.*

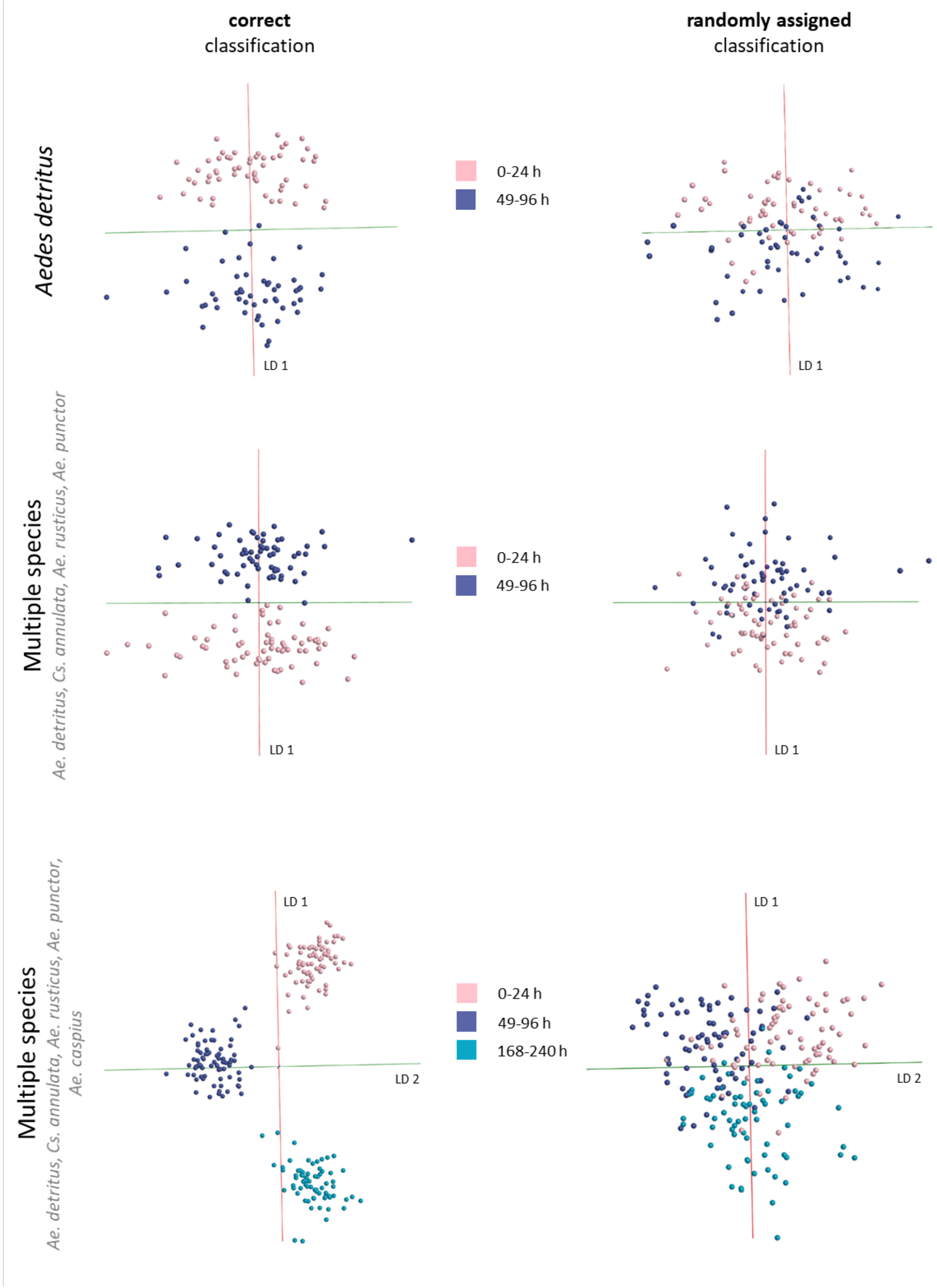

**Supplemental Figure 33:**

*The three main age models, with correct and randomly assigned classifications, were tested via cross-validation in OMB using the option 'Leave out 20 %' and a standard deviation of 5. The number of principal components used for model building are given in brackets underneath the tables. Two samples from the Aedes detritus age model were left out as 20 % of 107 samples results in a fractional number that is rounded to the nearest integer.*

Cross-validation *Offline Model Builder*:

**correct** classification

#### *Aedes detritus*

| Confusion matrix | 0-24 h | 49-96 h | Outlier |
|---|---|---|---|
| 0-24 h | 48 | 2 | 3 |
| 49-96 h | 8 | 44 | 0 |

| Spectra (n) | Pass | Fail | Outlier | Correct Classification (%) |
|---|---|---|---|---|
| 105 | 92 | 10 | 3 | 90.20 |

(50 PCs)

#### Multi-species

| Confusion matrix | 0-24 h | 49-96 h | Outlier |
|---|---|---|---|
| 0-24 h | 50 | 12 | 3 |
| 49-96 h | 4 | 58 | 1 |

| Spectra (n) | Pass | Fail | Outlier | Correct Classification (%) |
|---|---|---|---|---|
| 128 | 108 | 16 | 4 | 87.10 |

(60 PCs)

#### Multi-species incl. fed specimens

| Confusion matrix | 0-24 h | 49-96 h | 168-240 h | Outlier |
|---|---|---|---|---|
| 0-24 h | 66 | 0 | 6 | 3 |
| 49-96 h | 4 | 66 | 2 | 3 |
| 168-240 h | 0 | 3 | 64 | 2 |

| Spectra (n) | Pass | Fail | Outlier | Correct Classification (%) |
|---|---|---|---|---|
| 219 | 196 | 15 | 8 | 92.89 |

(100 PCs)

**randomly assigned** classification

#### *Aedes detritus*

| Confusion matrix | 0-24 h | 49-96 h | Outlier |
|---|---|---|---|
| 0-24 h | 26 | 28 | 1 |
| 49-96 h | 30 | 20 | 0 |

| Spectra (n) | Pass | Fail | Outlier | Correct Classification (%) |
|---|---|---|---|---|
| 105 | 46 | 58 | 1 | 44.23 |

(50 PCs)

#### Multi-species

| Confusion matrix | 0-24 h | 49-96 h | Outlier |
|---|---|---|---|
| 0-24 h | 30 | 32 | 2 |
| 49-96 h | 35 | 29 | 0 |

| Spectra (n) | Pass | Fail | Outlier | Correct Classification (%) |
|---|---|---|---|---|
| 128 | 59 | 67 | 2 | 46.83 |

(60 PCs)

#### Multi-species incl. fed specimens

| Confusion matrix | 0-24 h | 49-96 h | 168-240 h | Outlier |
|---|---|---|---|---|
| 0-24 h | 24 | 29 | 21 | 1 |
| 49-96 h | 24 | 22 | 27 | 2 |
| 168-240 h | 21 | 21 | 27 | 0 |

| Spectra (n) | Pass | Fail | Outlier | Correct Classification (%) |
|---|---|---|---|---|
| 219 | 73 | 143 | 3 | 33.80 |

(100 PCs)

**Supplemental Figure 34:**

*The PC-LDA separation achieved for the three main age models is presented here using the maximum number of principal components possible before overfitting (left side), as well as using only a quarter of possible PCs (right side).*

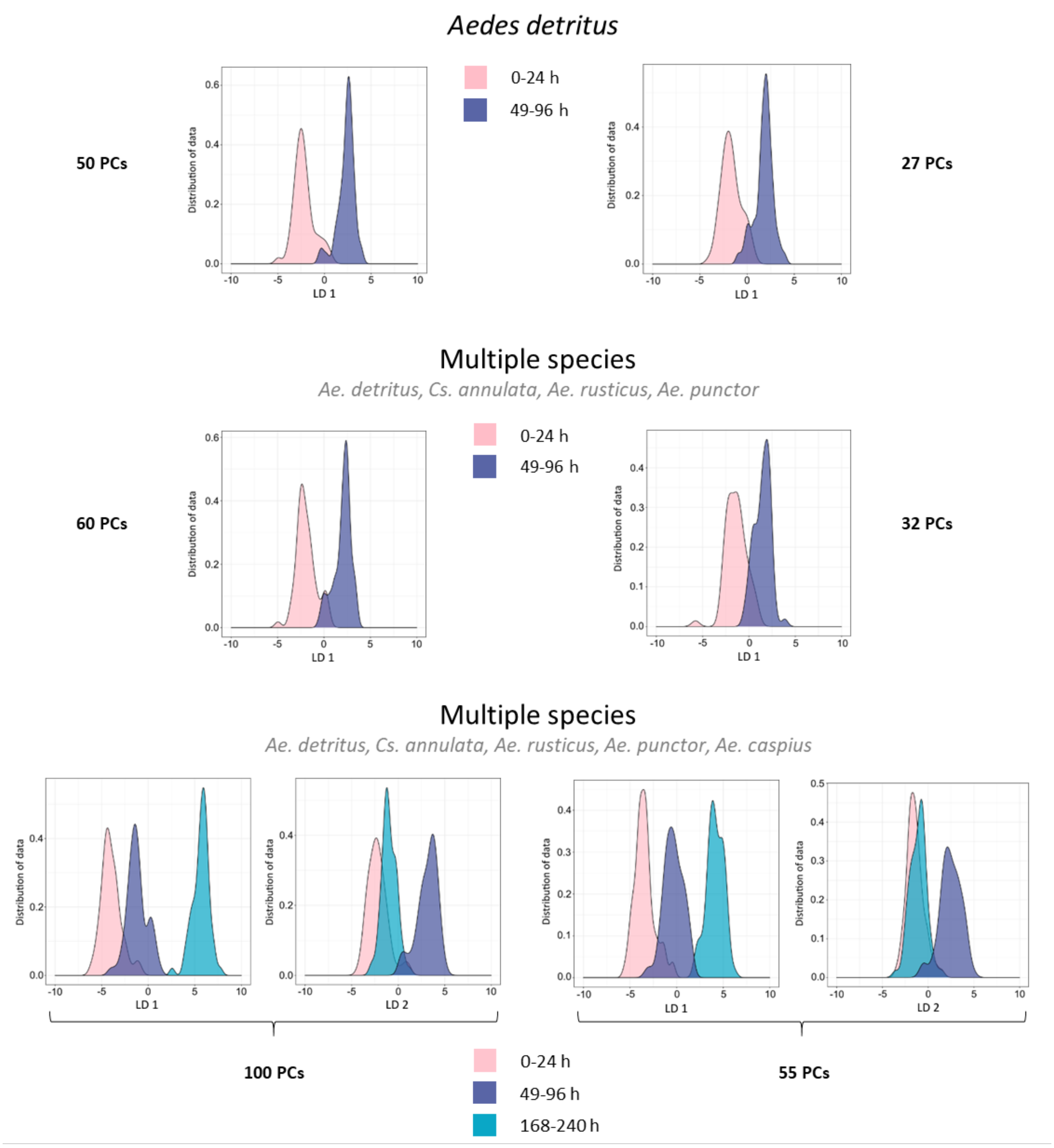

**Supplemental Figure 35:**

*After performing random forest analysis (repeated 10 times) on the age model (Figure 5.20) the R package 'randomForestExplainer' was used to determine the ion bins driving the separation process using a Top 10 approach. Four variables were identified as important in all 10 random forest runs. The intensities of all 219 samples were plotted for these bins in a boxplot diagram. A second panel with compacted y-axis is placed on top to show separated values for bin m/z 275.2*

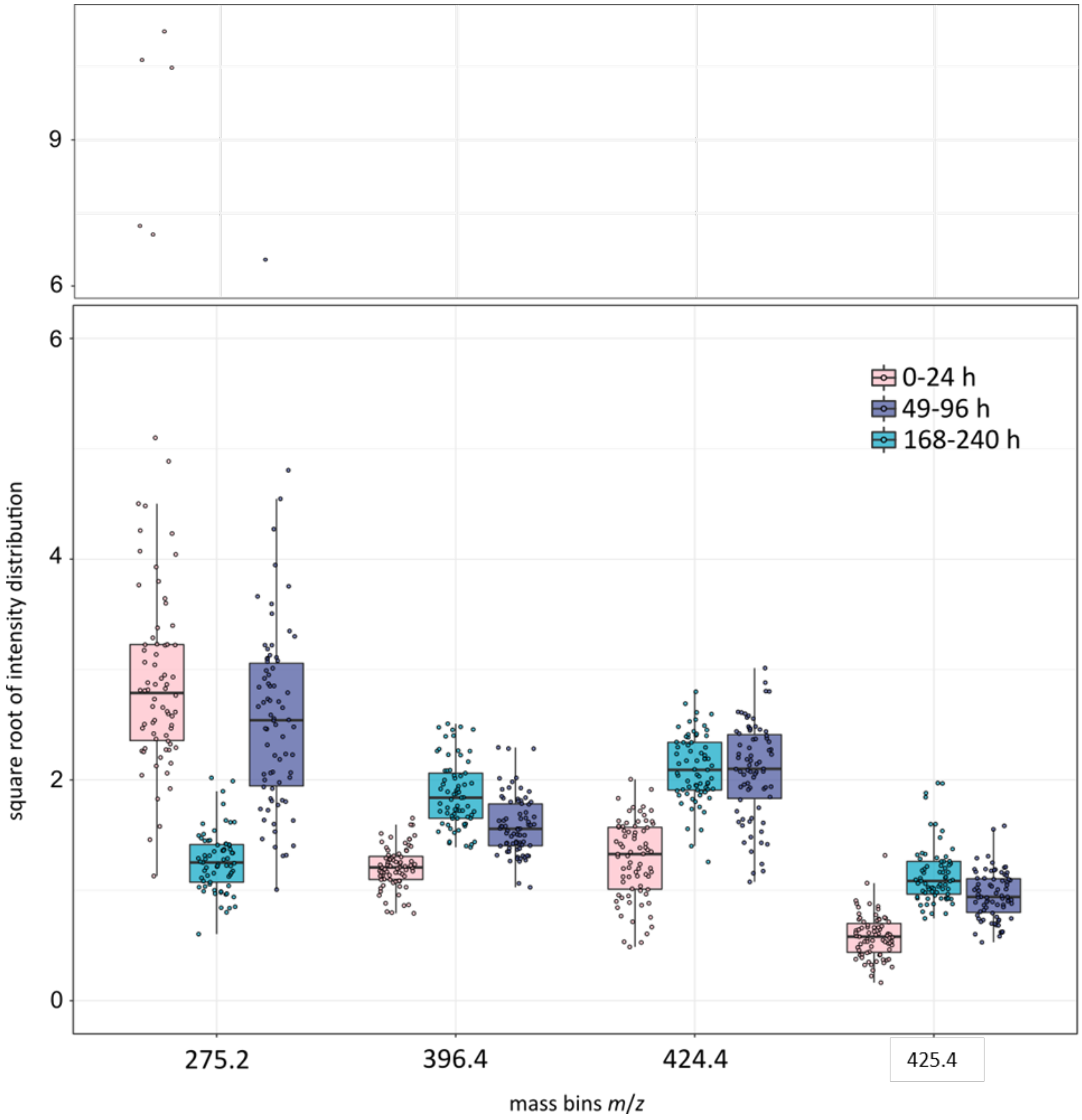

**Supplemental Figure 36:**

*Models separating age groups (1 day, 5-6 days, 14-15 days; 180 samples each) and species classes (*An. coluzzii, An. gambiae, An. arabiensis*; 180 samples each) were re-built in Offline Model Builder using a bin size of 1 m/z. Models were cross-validated in OMB ('Leave 20 % out', standard deviation 5).*

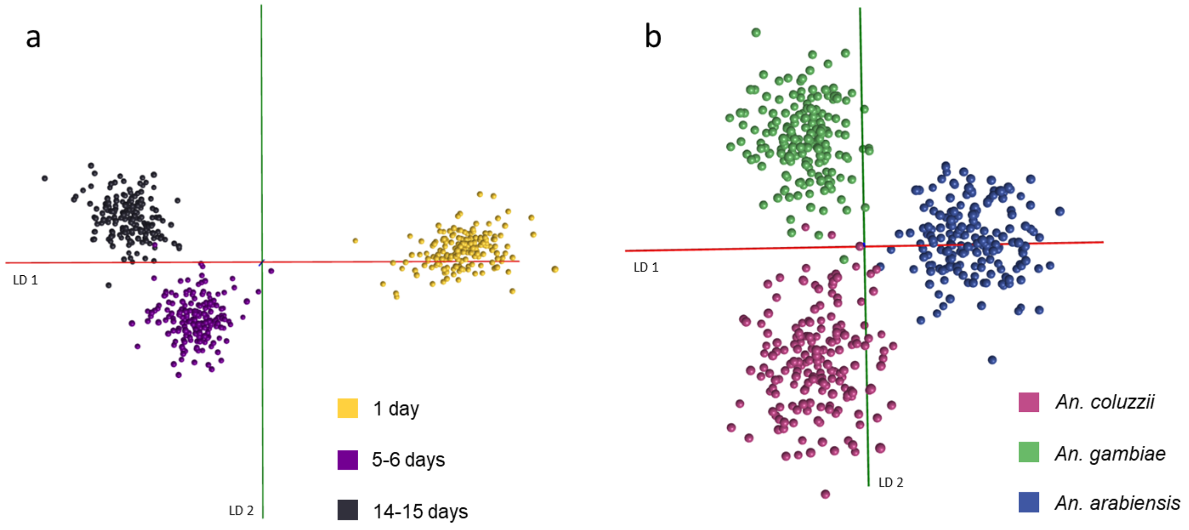


| Confusion matrix | 1 day | 5-6 days | 14-15 days | Outlier |
|---|---|---|---|---|
| 1 day | 180 | 0 | 0 | 0 |
| 5-6 days | 0 | 178 | 2 | 0 |
| 14-15 days | 0 | 4 | 176 | 0 |

| Spectra (n) | Pass | Fail | Outlier | Correct Classification (%) |
|---|---|---|---|---|
| 540 | 534 | 6 | 0 | 98.89 |

| Confusion matrix | *An. gambiae* | *An. arabiensis* | *An. coluzzii* | Outlier |
|---|---|---|---|---|
| *An. gambiae* | 176 | 2 | 2 | 0 |
| *An. arabiensis* | 3 | 173 | 4 | 0 |
| *An. coluzzii* | 8 | 3 | 169 | 0 |

| Spectra (n) | Pass | Fail | Outlier | Correct Classification (%) |
|---|---|---|---|---|
| 540 | 518 | 22 | 0 | 95.93 |

**Supplemental Figure 37:**

*Coordinates of the locations where immature mosquito specimens were collected.*

| Species | Latitude | Longitude |
|---|---|---|
| *Culex pipiens s.l.* | 53.30286 | -3.08296 |
| | 53.27714 | -3.06770 |
| | 53.28251 | -3.07218 |
| | 53.28186 | -3.03378 |
| *Culiseta annulata* | 53.30261 | -3.08247 |
| | 53.28253 | -3.07354 |
| | 53.28767 | -3.03597 |
| | 53.32771 | -3.04546 |
| *Aedes cantans* | 53.30811 | -3.04624 |
| | 53.28341 | -3.02950 |
| *Aedes rusticus* | 53.30811 | -3.04624 |
| | 53.28341 | -3.02950 |
| *Aedes punctor* | 53.26396 | -3.03076 |
| | 53.35253 | -3.13109 |
